# A novel mechanistic framework for precise sequence replacement using reverse transcriptase and diverse CRISPR-Cas systems

**DOI:** 10.1101/2022.12.13.520319

**Authors:** Y. Bill Kim, Elizabeth B. Pierce, Michael Brown, Brenda A. Peterson, Derek Sanford, Justin Fear, David Nicholl, Ellyce San Pedro, Grace M. Reynolds, Joanne E. Hunt, David G. Schwark, Sathya Jali, Nathaniel Graham, Zoe Cesarz, Tracey A. Lincoln Chapman, Joseph M. Watts, Aaron W. Hummel

**Affiliations:** Pairwise, Durham, NC, USA; Inceptor Bio, Morrisville, NC, USA

## Abstract

CRISPR/Cas systems coupled with reverse transcriptase (RT), such as the recently described Prime editing, allow for site-specific replacement of DNA sequences. Despite widespread testing of Prime editing, it is currently only compatible with type II CRISPR/Cas proteins such as *Streptococcus pyogenes* and *Staphylococcus aureus* Cas9. Enabling RT compatibility with other CRISPR/Cas domains, such as type V enzymes with orthogonal protospacer adjacent motif specificities and smaller protein size would expand the range of edits that can be made in therapeutic and industrial applications. We achieve this with a novel mode of DNA editing at CRISPR-targeted sites that reverse transcribes the edit into the target strand DNA (e.g., the complement of the PAM-containing strand), rather than the non-target strand DNA, as in Prime editing. We term this technology <u>R</u>NA encoded <u>D</u>NA <u>r</u>eplacement of <u>a</u>lleles <u>w</u>ith CRISPR (hereafter, REDRAW). We show that REDRAW extends the utility of RT-mediated editing beyond type II to include multiple type V CRISPR domains. REDRAW features a broad (8-10 bases) targeting window, at which all types of substitutions, insertions and deletions are possible. REDRAW combines the advantages of type V CRISPR domains with the extensive range of genetic variation enabled by RT-mediated, templated sequence replacement strategies.

## INTRODUCTION

Advances in genome editing allow researchers to modify the genomes of living organisms with increasing ease. While programmable site-specific nucleases such as zinc-finger nucleases (ZFNs), transcription activator-like effector nucleases (TALENs), and CRISPR/Cas9 allow facile disruption of genes through the generation of insertions or deletions (indels) arising from error-prone DNA repair by non-homologous end-joining (NHEJ) or double-stranded DNA breaks (DSBs), precise editing using nuclease technologies traditionally requires the use of homology-directed repair (HDR). HDR is active only during certain stages of the cell cycle and has variable efficacy across species and cell types. It is especially difficult to use for genome editing in plant systems, hampering the ability to generate precisely edited crops with desired phenotypes^1–3^. In contrast, recently described base editors^4^ and Prime editors^5^ mediate efficient, site-specific DNA sequence changes by localized deamination and reverse transcription, respectively. These precision genome editors have been applied to numerous purposes in research and product development for therapeutics and crop breeding.

Base editors use natural or engineered cytidine or adenosine deaminases to introduce single-nucleotide C to T or A to G transition mutations without relying on cellular DSB-repair pathways. Although useful in therapeutics, agriculture and elsewhere, base editors cannot efficiently cause other types of variation, including transversions, multiple base substitutions, and precise indels, limiting the range of genetic variation that can be created, assessed, and used with a base editing toolbox. Thus, new technologies capable of efficient transversions, multi-base substitutions, and indels can complement a base editing toolbox by extending the range of genetic variation to install the most biologically relevant allele type for any native gene modification project.

Prime editing addressed many of the shortcomings of base editing by using templated reverse-transcription to introduce small-scale point mutations, insertions, and deletions at CRISPR target sites in the genome. Prime editing features a Cas9 H840A nickase fused to Moloney Murine Leukemia Virus (M-MuLV) reverse transcriptase (RT)^5^. After binding, the Prime editor nicks the strand with the protospacer adjacent motif (PAM; hereafter this strand is called the non-target strand, or NTS, because it is the strand not paired with the guide RNA spacer sequence), the nicked DNA is primed by hybridizing to the RNA sequence 3′ of the single guide RNA (sgRNA), and the edit encoded by the RNA template is then reverse transcribed into the nicked DNA strand by the RT^5^. This intricate mode of action requires two biomolecular features of the enzyme-target complex for successful reverse transcription: (1) the extended RNA sequence must be able to hybridize with the R-loop structure formed by the nicked DNA, and (2) the reverse transcriptase must be able to access the hybridized RNA/DNA duplex, which is in close vicinity to the Cas9 ribonucleoprotein (RNP) domains. Perhaps due to architectural constraints of the enzyme-target complex, Prime editing systems have so far been limited to type II CRISPR/Cas proteins, most prominently Cas9 from *Streptococcus pyogenes* and *Staphylococcus aureus*.

The type II CRISPR system can be generally characterized as containing two DNAse domains (an HNH and a non-continuous RuvC), requiring a 3′ G-rich PAM, and creating blunt-end DNA products when used as a nuclease. Type II systems are larger than other CRISPR systems that have single nuclease domains, such as the type V CRISPR systems^6^. Cas12a (formerly Cpf1) is a widely used type V CRISPR protein that contains a continuous RuvC without HNH, recognizes 5′ T-rich PAMs, is usually smaller than Cas9, and generates 5′ overhangs when used as a nuclease on double-stranded DNA. Moreover, Cas12a processes its own CRISPR RNA (crRNA), which facilitates multiplex genome editing by simply expressing several crRNA precursors in a single molecule array using one promoter^6^. Expanding the compatibility of RT-mediated precision genome editing to alternate CRISPR domains would expand the ability to manipulate any genomic sequences, especially in organisms with A/T rich genomes, such as plants.

Here, we significantly extend the capabilities of our base editing toolbox with a reverse transcriptase editing system that enables all types of small, precise edits and is compatible with both type II and type V CRISPR systems. Dubbed REDRAW for <u>R</u>NA encoded <u>D</u>NA <u>r</u>eplacement of <u>a</u>lleles <u>w</u>ith CRISPR (hereafter, simply called REDRAW), this system expands the functionality of reverse-transcriptase-mediated precision genome editing.

## RESULTS

### Precise reverse transcription into the target strand DNA

Prime editing uses a NTS nickase Cas9 (nCas9) fused to an engineered M-MuLV RT penta-mutant (RT(5M)) and a Prime editing guide RNA (pegRNA) comprising a primer-binding site (PBS) and reverse-transcriptase template (RTT) which encodes the desired mutation^5^. The reverse-transcription complex is formed when nCas9 induces a nick on the exposed NTS of the R-loop DNA, which is then hybridized by the complementary sequence in the PBS^5^. The resulting DNA-RNA hybrid primes the reverse transcriptase reaction wherein RT(5M) encodes the desired mutation into the DNA^5^.

We tested whether Cas12a is compatible with the Prime editing architecture (**Supplementary Figure 1a**). A previous study has shown that AsCas12a can be converted into a catalytic NTS nickase by mutating an arginine in position 1226 into an alanine^7^. This induces kinetic impairment in the nuclease which drastically slows cleavage of the target strand DNA which normally follows a cut in the NTS DNA. We transfected HEK293T cells with plasmids encoding RT(5M) fused to LbCas12a R1138A (the equivalent of the R1226A mutation in AsCas12a) in both N-terminal and C-terminal fusion orientations along with plasmids encoding the modified crRNA that included typical PBS and RTT extension lengths used in Prime editing. However, no precise edit was detected via high-throughput sequencing at two different loci, despite extensive testing of PBS lengths ranging from 9-79 bases and RTT lengths ranging from 8-80 bases paired with either N-terminal or C-terminal RT fusion and recruitment of RT(5M) to Cas12a R1138A with the SunTag antibody system^8^ (**Supplementary Figure 1b,c**). Providing a nearby nick on the target strand to promote productive edit incorporation, either by adding another crRNA or by adding a Cas9 nickase and a guide RNA, did not enable editing (**Supplementary Figure 1c**). We hypothesized that the R-loop structure generated by Cas12a, in contrast to Cas9, is not a preferred substrate for reverse transcriptase. This is consistent with previous literature that demonstrated variable, and often reduced, accessibility of single-stranded DNA (ssDNA) in the R-loop of Cas12a base editors by cytidine deaminase^9^.

To overcome this challenge, we envisioned that the RT(5M) may be able to access the target strand DNA and polymerize it upon providing a DSB with an RNA template complementary to the target strand (**Figure 1a**). RT-mediated polymerization would result in elongation of the target strand DNA with the edit encoded in the RNA template followed by permanent incorporation through DNA repair. Transfection of HEK293T cells with plasmids encoding nuclease active LbCas12a, independently expressed RT(5M), and a crRNA extended with an RNA sequence complementary to the target strand DNA adjacent to the DSB resulted in the generation of precise edits, with efficiencies of up to 0.8% of total reads depending on the combination of PBS and RTT sequences used (**Figure 1b**). The presence of RNA template complementary to the target strand of DNA, as well as a functional RT, was necessary for precise editing. We refer to this novel mechanism of reverse transcription into the target-strand DNA as REDRAW (<u>R</u>NA encoded <u>D</u>NA <u>r</u>eplacement of <u>a</u>llele <u>w</u>ith CRISPR). crRNAs that contain extended segments of PBS and RTT with target strand complementarity are referred to as <u>t</u>arget-<u>a</u>llele <u>g</u>uide RNAs (tagRNAs; **Supplementary Figure 2a**).

**Figure 1.**
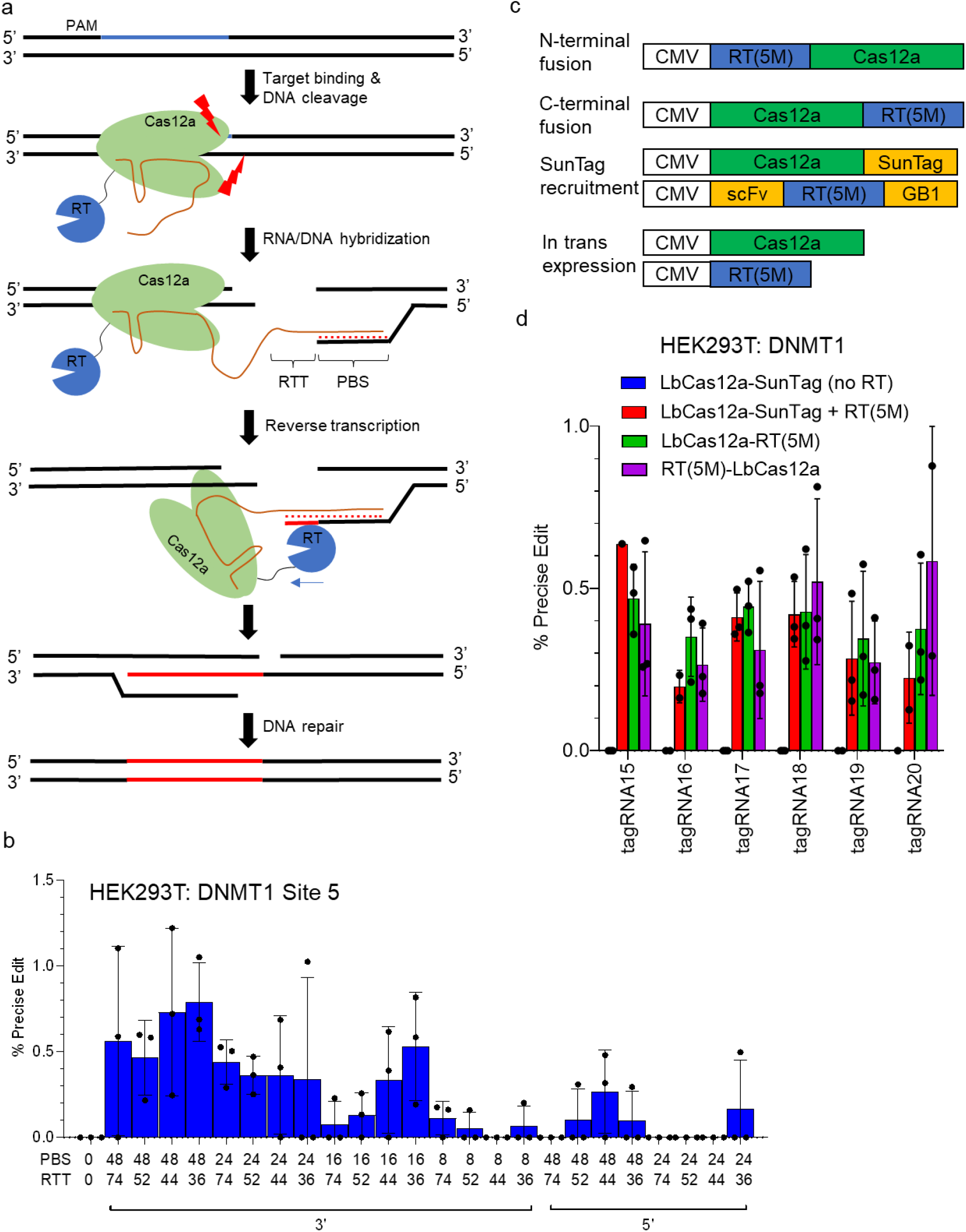
Development of REDRAW. (a) REDRAW editing strategy, which involves Cas12a-mediated DSB generation followed by target-strand reverse transcription of DNA. (b) Optimization of RTT and PBS lengths compatible with REDRAW. Varying lengths of PBS and RTT were attached to either 5′ or 3′ end of crRNA. All tagRNAs were expressed from a U6 promoter. (c) REDRAW architectures tested. RT(5M) was fused to the N-terminus or C-terminus of Cas12a, as well as recruited to Cas12a via an antibody system known as SunTag (8 copies of GCN4 epitopes attached to Cas12a and co-expressed RT(5M) fused to GCN4 scFv). (d) REDRAW activity using various REDRAW architectures.

As a preliminary optimization step, we explored the range of PBS and RTT lengths and orientations for tagRNAs compatible with REDRAW. LbCas12a is known to process its own crRNA by cleaving the RNA 5′ to the pseudoknotted hairpin that forms the crRNA scaffold^6^. To enable testing of both 5′ and 3′ fusion orientations for PBS and RTT attachments, LbCas12a with a mutation that abolishes its RNA-processing function^10^ was used in the initial experiments. Across 2 different target sites in HEK293T cells, we simultaneously varied PBS from 8-48 bases and RTT from 36-72 bases on either the 5’ or 3’ end of the crRNA. We identified that PBS of 48 bases offered the most efficient editing, while various RTT lengths were well tolerated. REDRAW accommodated PBS and RTT extensions both 5′ and 3′ of the crRNA, although 3′ extension resulted in more robust activity (**Figure 1b, Supplementary Figure 2b**).

To identify the most effective protein architecture we explored different strategies to deliver the RT(5M) to the target (**Figure 1c**). We fused RT(5M) to LbCas12a in N-terminal and C-terminal orientations and recruited it to the target with the SunTag antibody system by adding eight GCN4 epitope tags to the Cas12a C-terminus and an anti-GCN4 antibody^8^ to the RT(5M) N-terminus. Both fusion and recruitment approaches yielded similar REDRAW editing efficiencies (**Figure 1d**). For further optimizations we selected the N-terminal fusion of RT(5M) to LbCas12a along with optimal tagRNA extension lengths (PBS ~48bp, RTT ~52-72bp) fused to the 3′ end of crRNA. This configuration is termed REDRAW Editor 1 (RE1).

### Optimizing the protein fusion and tagRNA stability

To further improve REDRAW, we explored biochemical approaches to increase the stability of various nucleic acid components. First, we hypothesized that the stability of the DSB intermediate is important for enabling robust strand invasion by the PBS sequence and RNA-DNA hybridization for reverse-transcription. We envisioned that stabilizing the transiently exposed ssDNA after nuclease cleavage may prevent the sequestration of the intermediate by factors involved in non-homologous end joining (NHEJ), which presumably reduces the efficiency of REDRAW by leading to INDEL byproducts. Therefore, we identified and tested 5 single-stranded DNA (ssDNA) binding proteins by fusing them in either N-terminal or C-terminal orientation to RE1. These were Rad52 from *Homo sapiens*^11^; ssDNA binding protein RecA from *Escherichia coli* (EcRecA) and *Bacillus subtilis* (BsRecA)^12^; ssDNA binding protein from T4 bacteriophage (T4SSB)^13^, and Brex27 peptide motif that was previously shown to recruit RAD51, which stabilizes ssDNA during DSB events^14^. Plasmids encoding various fusion constructs and 2 tagRNAs targeting 2 different endogenous loci were transfected into HEK293T cells, and the precise editing efficiency was compared against RE1 without any fusion. While most fusion constructs did not increase REDRAW efficiency, and some negatively impacted the outcome, fusion of a 36 aa Brex27 peptide resulted in an average 1.8- and 1.3-fold increase in REDRAW efficiency as an N-terminal and C-terminal fusion to RE1, respectively (**Figure 2a, Supplementary Figure 3**).

**Figure 2.**
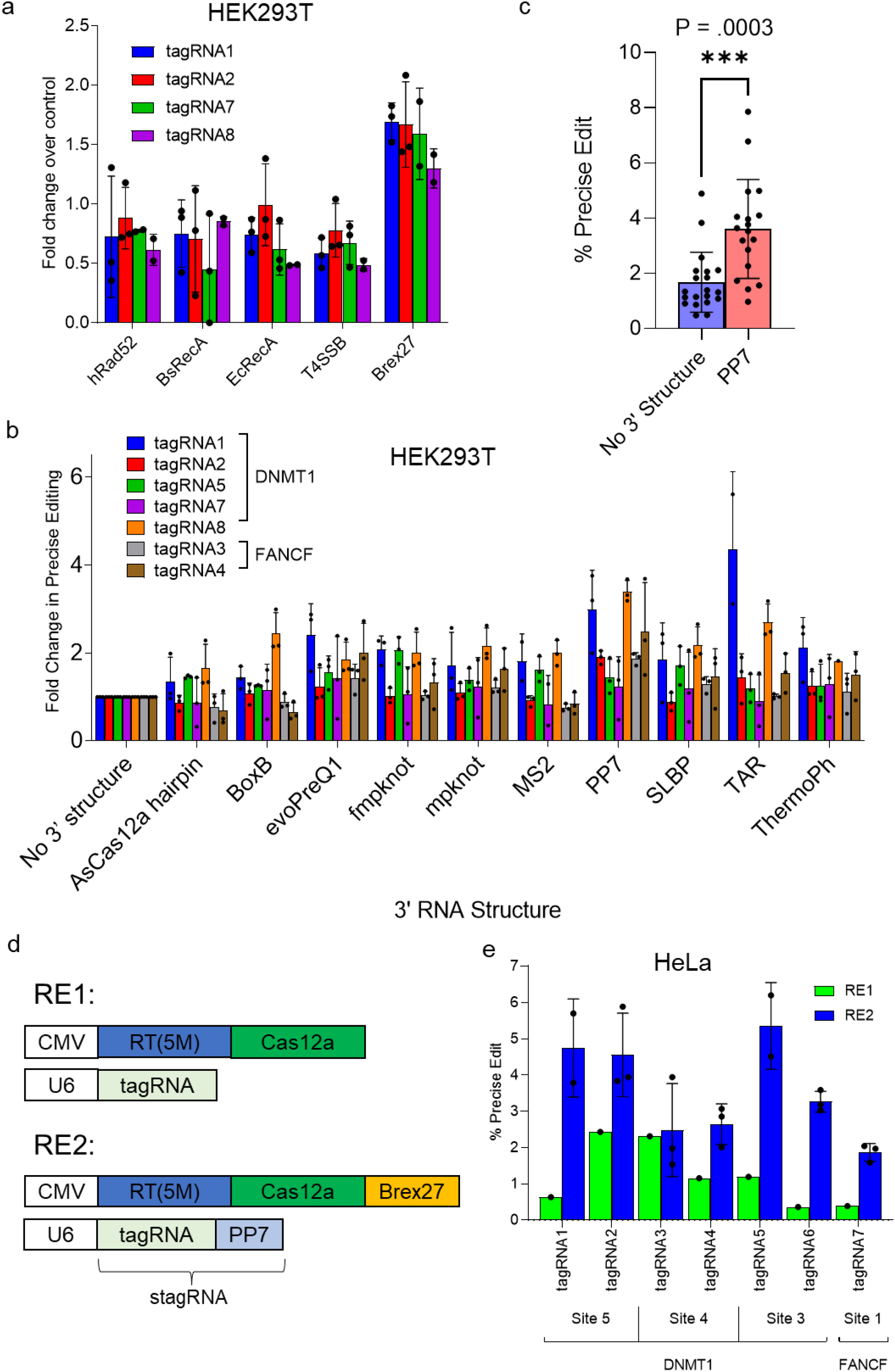
Engineering REDRAW components. (a) Identification of ssDNA binding proteins (ssDNA BP) that improve precise editing efficiency. Five proteins (hRAD52, *Bacillus subtilis* RecA (BsRecA), *Escherichia coli* RecA (EcRecA), ssDNA binding protein from bacteriophage T4 (T4SSB), and BRCA2 subunit Brex27 that recruits and stabilizes RAD51-DNA interaction) were fused to the N-terminus of RT(5M)-LbCas12a and tested with 4 tagRNAs. (b) Screening of structured RNA motifs that improve the precise editing efficiency by reducing tagRNA degradation. RT(5M)-LbCas12a was tested with 8 different tagRNAs containing 3′-attached RNA motifs including AsCas12a crRNA hairpin, BoxB, evoPreQ1, fmpknot, mpknot, MS2, PP7, SLBP, TAR, and ThermoPh. The relative efficiency compared to an unmodified tagRNA was plotted. (c) 3′ attachment of PP7 hairpin led to the most consistent increase in precise editing. (d) Diagram showing the initial REDRAW editor, RE1, and the improved REDRAW editor, RE2, which contains a Brex27 peptide fusion and tagRNA stabilized with PP7 (stagRNA). (e) RE2 mediates 4.5-fold greater precise editing activity on average, compared to RE1 across 8 different stagRNAs in 2 genomic loci in HeLa cells.

Next, we sought to increase the stability of the tagRNA to prevent degradation of the extended sequence that may reduce the amount of full-length tagRNAs in the cell capable of forming a viable substrate for reverse transcription. The use of structured RNA motifs has previously been shown to increase the stability of transcribed RNA^15^. To identify structured RNA motifs that best stabilize the tagRNA without hindering hybridization with the target strand DNA or Cas12a function, we screened a custom curated panel of RNA hairpins and pseudoknots added to the 3′ end of the tagRNA. Among the motifs tested were hairpins such as MS2, PP7, and BoxB, from bacteriophages, three from HIV (ThermoPh^16^, SLBP^17^, TAR^18^), the AsCas12a crRNA hairpin, and the pseudoknots evoPreQ1, mpknot, and fmpknot (a truncated form of mpknot) which have been successfully used to stabilize pegRNAs in Prime editing^15^. Plasmids encoding RE1 along with modified tagRNAs with hairpin extensions were transfected into HEK293T cells and the precise editing was measured using high-throughput sequencing. The bacteriophage PP7 hairpin consistently improved the frequency of precise editing by an average of 2.3-fold (**Figure 2b,c**). The PP7-stabilized tagRNA is hereafter termed stagRNA (stabilized tagRNA). Together the C-terminal fusion of Brex27 to RE1 (RT(5M)-LbCas12a-Brex27) coupled with the use of the stagRNA results in a 4.5-fold average increase in precise editing efficiency compared to RE1, and this configuration is hereafter referred to as RE2 (**Figure 2d,e**). In summary, RE2 improves REDRAW editing with a protein fusion optimized to stabilize the ssDNA intermediate of the DSB, and with a stabilized tagRNA.

### REDRAW enables a range of DNA modifications in multiple cell types

The REDRAW mechanism installs edits on the PAM distal side of the DSB. In theory, the desired edits can be installed anywhere near the target site by adjusting the length and composition of the RTT sequence and the position of the mutations therein. To assess this theory, we comprehensively tiled two-base transversion edits from 32 base pairs upstream of the PAM through the full length of the protospacer sequence, while keeping the total length of RTT constant. We screened the corresponding stagRNAs by measuring the quantity of genomic products with desired changes in HEK293T cells. Precise edits could be incorporated in a continuous 25 base pair window ranging from 1 base pair upstream of the PAM (position −5; the first position of the protospacer is 1) to position 21 near the end of the protospacer sequence, with best efficiency obtained when the desired mutation was placed near the midpoint of the protospacer (positions 9-15; **Figure 3a**). The absence of precise editing outside the protospacer and PAM may be the result of repetitive binding and cutting by Cas12a when REDRAW editing does not disrupt the target sequence, resulting in INDEL mutations that sequester these edit types from the pool of precision outcomes. However, this does not explain the gradual decrease in efficiency as the edit position is moved from 12 to −5. More investigation is necessary to fully understand the constraints that define the REDRAW editing window.

**Figure 3.**
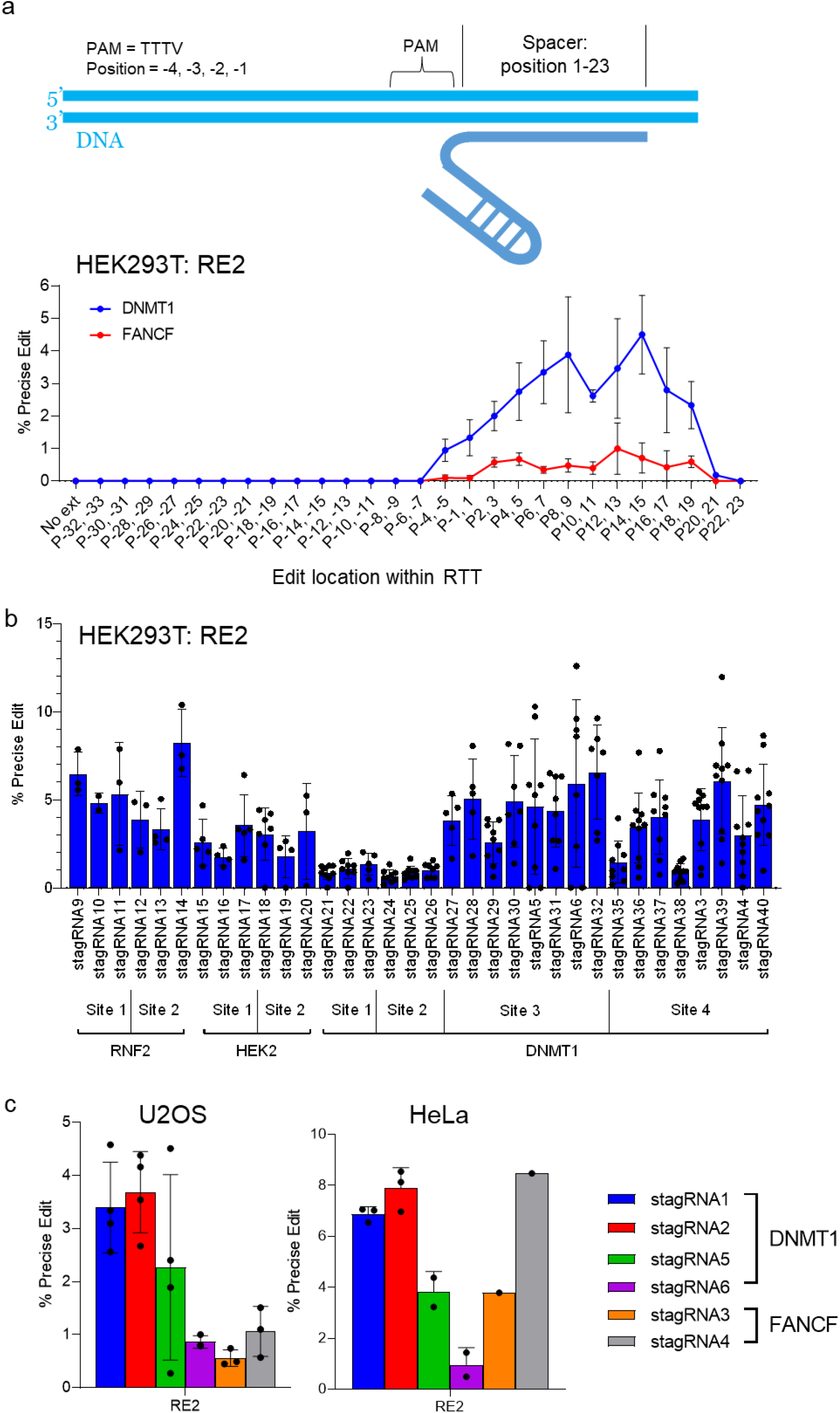
Characterization of the REDRAW targeting window and activity across genomic sites and cell types. (a) Determining the optimal targeting window of RE2. Double transversion mutations were tiled within the RTT ranging from position −32 to 23. At both sites tested, optimal editing was observed when the RTT encoded edits within position 9-15. (b) RE2 was tested across a wide range of genomic contexts at RNF2, HEK2, and DNMT1 loci with several stagRNAs in HEK293T cells. Full stagRNA information can be found in **Supplementary Note 1**. (c) RE2 is active in cell types other than HEK293T, including U2OS and HeLa cells.

To understand the activity of RE2 at different genomic contexts we tested 34 different stagRNAs encoding various edits targeted to three different loci (all stagRNA sequences used in this study can be found in **Supplementary Note 1**), detecting up to 8% frequency of precise editing by high throughput sequencing (**Figure 3b**). Encouraged by the robust activity of RE2 in HEK293T cells and by the improved efficiency of RE2 over RE1 in HeLa cells (**Figure 2e**), we expanded our investigation of REDRAW in other cell types. We compared RE2 in HeLa and U2OS cells by transfecting RE2-expressing plasmids together with six different stagRNAs targeted against two genomic loci. We observed maximal REDRAW editing of 3.6% and 8% frequency in U2OS and HeLa cells, respectively (**Figure 3c**). Importantly, lipofectamine transfection was used exclusively throughout this study. We expect REDRAW editing efficiency to improve with nucleofection due to its superior delivery frequencies across cell types. Taken together, RE2 can create precise DNA edits in various genomic sites in multiple human cell lines.

RT editors are most valuable to therapeutic and agricultural genome editing toolboxes for their ability to extend beyond the editing range provided by base editors. For this reason, we were especially interested in assessing REDRAW for its ability to install various single- and multi-base precise edits including transversions, insertions and deletions. We started by encoding the RTT with transversion mutations targeted to positions 12 through 15 of the protospacer, tiling each of the four positions as single, double, triple, and quadruple mutations at two genomic loci. All edits were possible at both loci, with maximal editing up to 6.6% (**Figure 4a**). We next compared the efficiency of 2-nucleotide transition or transversion edits against 3-base deletions at six different loci. At each locus, the installation frequency was similar for the transition, transversion and deletion edits (**Figure 4b**).

**Figure 4.**
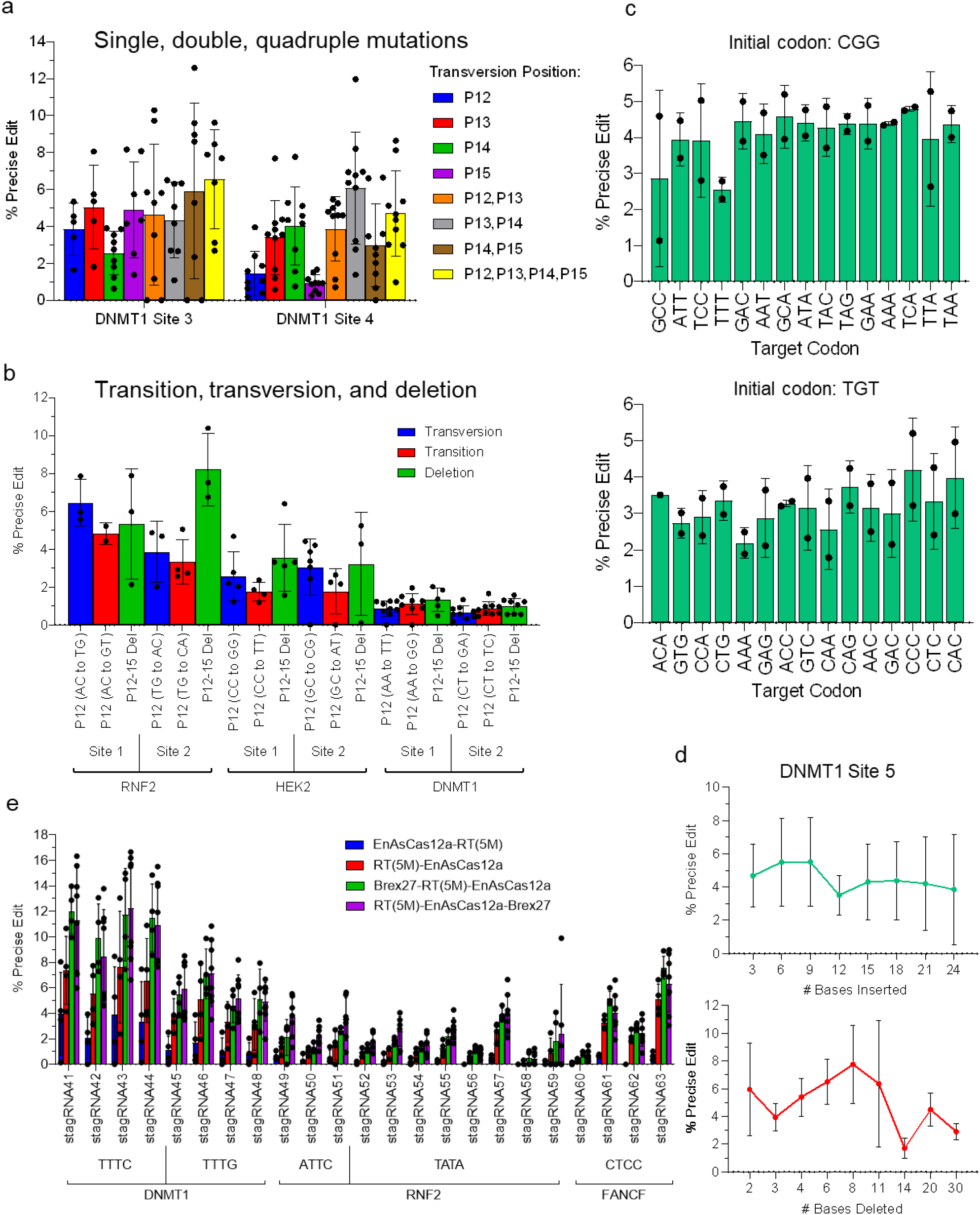
Mutational scope of REDRAW editors. (a) RE2 can generate single, double, and quadruple mutations with similar efficiency. (b) Transition, transversion, and deletion are introduced with similar efficiencies. (c) RE2 can introduce all combinations of nucleotide mutations at the same 3 bases with similar efficiencies at 2 different genomic loci. (d) RE2 can create defined deletions and insertions of varying lengths. (e) EnAsCas12a REDRAW provides expanded PAM accessibility.

When engineering crop traits for maximum performance, there is considerable value in making and screening allelic series to identify the optimal phenotype. We were thus interested in assessing REDRAW for its potential to install all possible amino acid substitutions at a particular codon. As a proxy for systematic substitution of all possible codons, we instead designed stagRNAs encoding all possible triplet mutations in which all three bases were different from the original base at each position. These were tested in combination with RE2 at 2 different target sites in HEK293T cells, generating relatively consistent editing frequencies of 4.1% and 3.2%, respectively (**Figure 4c**). This indicates REDRAW enables predictable installation of diverse 3-nucleotide motifs and forecasts its suitability for generating allelic series of amino acid substitutions in trait engineering.

The ability to precisely insert or delete specific sequences is an additional extension beyond the capabilities of base editors and is therefore highly desirable. To assess the ability of REDRAW to install insertions of various lengths, stagRNAs were designed with insertions ranging from 3 to 24 bases at two genomic loci. After transfecting plasmids encoding these stagRNAs and RE2, we observed similar insertion efficiency for the entire range (**Figure 4d and Supplementary Figure 4**), suggesting REDRAW can efficiently install insertions larger than 24 bases. We next assessed REDRAW for the introduction of precise deletions of various lengths. stagRNAs encoding deletions ranging from 2 to 30 bases at two genomic loci were transfected with RE2, resulting in detectable deletions across the tested range. However, there was a clear trend of decreasing efficiency with longer deletions, although there were locus-specific differences in when this penalty occurred (**Figure 4d and Supplementary Figure 4**). We speculate the reduced efficiency of longer deletions could be due to inefficient DNA repair when homology between the NTS and the edited target strand is distant from the DSB site.

In summary, REDRAW can effectively install a wide variety of biologically relevant, precise edits at multiple loci. Importantly, this includes single-base transversions, multi-base substitutions, and precise indels, providing a significant extension to the capabilities of base editors and defining a genome editing toolbox that can deliver all types of small, precise edits.

### Use of Cas12a enzymes with altered PAM specificities improves the targeting range of REDRAW

Biologically relevant genome editing for therapy and crop improvement requires installation of specific alleles at exact locations. Non-canonical PAM recognition is therefore useful for positioning REDRAW editors at the optimal location. RR-Cas12a variants recognize TYCV and wild type TTTV PAM sequences^19^ and EnAsCas12a uses VTTV, TTTT, TTCN, and TATV PAM sequences^20^.

To determine the compatibility of Cas12a PAM variants with REDRAW, we first tested EnAsCas12a, which is an engineered AsCas12a with expanded PAM access and robust activity in human cells^20^. We tested both the N-terminal and C-terminal fusions of RT(5M) to EnAsCas12a, with and without the Brex27 peptide used in RE2. These proteins were co-expressed with 23 stagRNAs across 3 genomic loci in HEK293T cells. Like LbCas12a-based REDRAW, the N-terminal RT fusion was 4.0-fold more efficient than the C-terminal fusion and was preferred to the C-terminal fusion orientation with all 23 stagRNAs assessed (**Figure 4e**). Appending a C-terminal Brex27 peptide further boosted efficiency, as RT(5M)-EnAsCas12a-Brex27 resulted in up to 12.2% precise double transversion editing at both TTTV and non-canonical PAM sites. To further expand target accessibility, we created RR-RE2 by replacing wildtype LbCas12a in RE2 with an LbCas12a containing a double-arginine (RR) mutation that expands its PAM access^19^. RR-RE2 delivered precise editing up to 3.4% in HEK293T cells (**Supplementary Figure 5**), confirming that Cas12a variants with altered PAM recognition and improved activity are compatible with REDRAW. Taken together, the targeting scope of REDRAW is substantially increased by using RR-RE2 and EnAs-RE2, increasing the probability that a suitable PAM site will be available to position the optimal REDRAW editing window directly over the target base(s).

### Concurrent nicking of non-target strand DNA increases REDRAW efficiency

During REDRAW editing, the exposed 3′ end of the target strand DNA is extended by reverse transcription, incorporating the desired edit encoded in the RNA template into the DNA. Subsequently, there exists a mismatched intermediate where the newly synthesized target strand DNA imperfectly pairs with the native sequence on the NTS. This intermediate is prone to recognition and repair by the DNA mismatch repair (MMR)^21^ pathway, which can lead to conversion of the edit into the NTS, a favorable outcome for REDRAW efficiency. However, MMR can also convert the target strand edit back to the original sequence, thereby reversing the REDRAW edit before it is permanently incorporated into the target site. In eukaryotic cells, mismatch detection is sensitized by the presence of a nick in the DNA strand. Mismatched bases in the nicked strand are more likely to be replaced^21^ than the mismatched bases opposite them in the unnicked strand. This is the basis for the significantly improved editing outcomes observed when nicking of the unedited strand was added to tools that generate MMR substrates as intermediates, as with the 3^rd^ generation cytosine base editor (BE3)^4^ and Prime editor (PE3)^5^. These observations are consistent with a REDRAW model where nicking the NTS promotes MMR-mediated removal of the wild type bases complementary to the reverse transcribed target strand DNA that encode the desired mutation.

We therefore sought to increase the efficiency of REDRAW during resolution of the partially edited intermediate by causing a nick in the NTS to encourage MMR with the edited target strand as the template (**Figure 5a**). Unlike Prime editing, which uses Cas9 as a ssDNA nickase, creating a secondary nick with REDRAW is not as simple as adding a second crRNA because REDRAW uses a nuclease active Cas12a. However, the presence of strategically placed mismatches within the crRNA spacer has been previously shown to attenuate the kinetics of DNA cleavage in Cas12a, resulting in a NTS nick with wildtype Cas12a^22^. We therefore hypothesized this strategy could enable NTS nicking by the Cas12a nuclease in REDRAW as well. We first confirmed the mismatched crRNA mediated Cas12a nicking behavior *in vitro* by incubating purified Cas12a nucleases with supercoiled plasmid DNA and visualizing the migration pattern on an agarose gel (**Supplementary Figure 6a**). As expected, placement of mismatches at positions 12-15 led LbCas12a to behave as a nickase. Combining mismatches in these positions into double, and in some cases triple, mutations led to even greater nicking activity *in vitro* (**Supplementary Figure 6a**).

**Figure 5.**
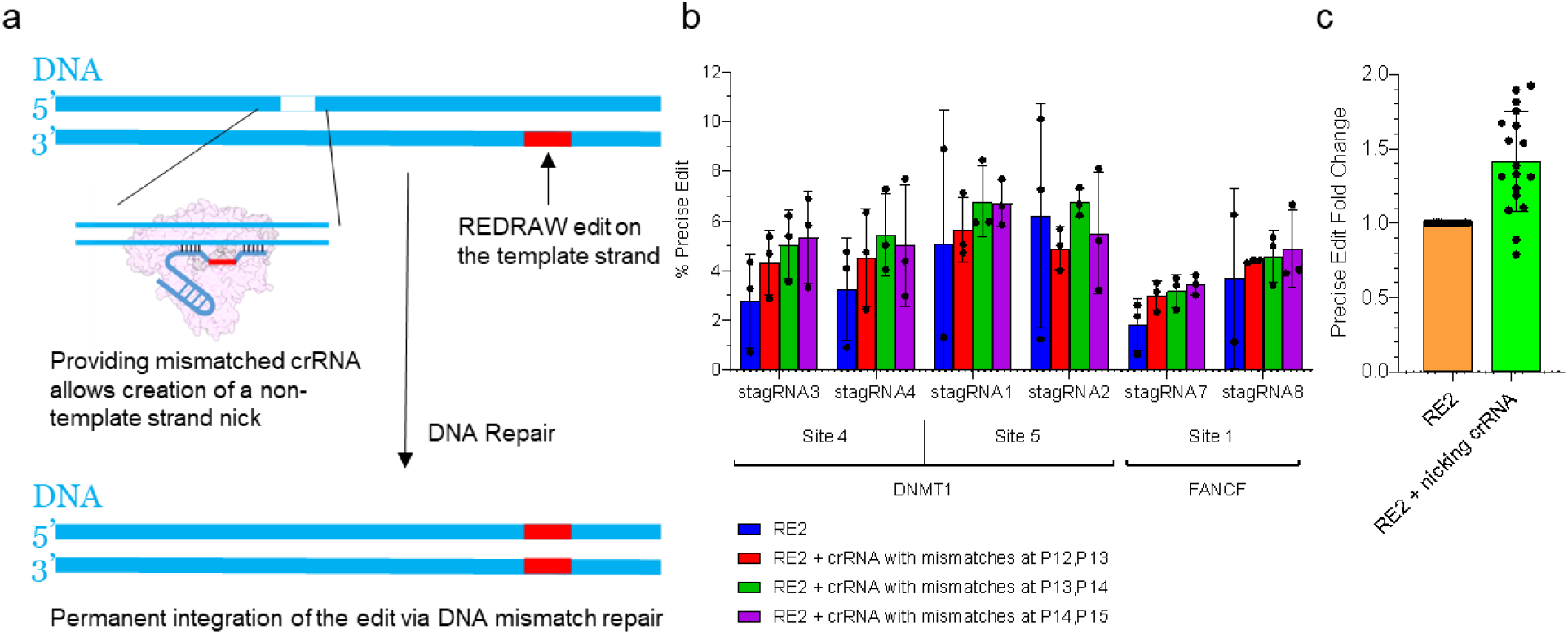
Non-target strand nicks enhance REDRAW efficiency. (a) Schematic of non-target-strand DNA nick generated by a secondary crRNA with mismatches in the spacer region. Nicking is expected to enhance the permanent integration of REDRAW edit through DNA mismatch repair. (b) Precise editing using 6 stagRNAs in 3 genomic loci in the presence or absence of secondary nicking crRNAs. (c) The use of mismatched crRNAs in conjunction with RE2 led to 1.4-fold increase in editing efficiency.

To verify whether providing a mismatched crRNA during REDRAW increases the editing efficiency in human cells, we transfected HEK293T cells with RE1 targeting DNMT1 with an additional plasmid expressing crRNAs containing mismatches in the spacer sequence. Combining REDRAW with mismatched crRNAs designed to nick the NTS at position −5 relative to the tagRNA spacer resulted in maximal 5.5- and 8.7-fold increase in precise editing, respectively, across the two tagRNAs tested (**Supplementary Figure 6b**). The use of crRNAs with double mismatches at other bases, or with crRNAs with triple mismatches, caused inconsistent improvement, indicating the best nickase configuration *in vitro* may not be optimal for promoting REDRAW activity in cells and further suggesting target site-dependency for the best nickase design. The use of crRNAs with no mismatches, which would be expected to introduce a second DSB, eliminated precise REDRAW editing (**Supplementary Figure 6b**).

To further evaluate the NTS nicking strategy for improving REDRAW efficiency, RE2 using six different stagRNAs was paired with mismatched crRNAs designed to nick the NTS at position +45, −5, and −14 at DNMT1 Site 5, DNMT1 Site 4, and FANCF Site 1, respectively, in HEK293T cells. This modestly increased REDRAW efficiency at all targets individually (**Figure 5b**), and in aggregate showed a 1.4-fold average improvement over RE2 alone (**Figure 5c**). The use of RE2 with a mismatched crRNA is termed REDRAW Editor 3 (RE3). In summary, a secondary crRNA causing NTS nicking can increase REDRAW editing, presumably by manipulating the MMR process to favor permanent incorporation of the target strand edit over permanent reversal of the partially edited target.

### REDRAW is compatible with distant CRISPR family members

In theory, REDRAW is compatible with any protein that can create DSBs and recruit RT with an RNA template to the target site. To demonstrate REDRAW’s compatibility with other CRISPR proteins we first tested Cas12b, a type V enzyme containing a single, continuous RuvC domain and using both a crRNA and a tracRNA^23^. We replaced Cas12a in RE2 with BhCas12b v4, a Cas12b variant that was engineered for robust activity in human cells^23^, to generate BhCas12b-RE2. TagRNAs were designed for BhCas12b by attaching PBS and RTT sequences and the PP7 end-stabilization motif at the 3’ end of BhCas12b sgRNA. After transfecting plasmids expressing BhCas12b-RE2 and the end-stabilized tagRNA into HEK293T cells, we observed low but detectable editing at several genomic loci, confirming that REDRAW is compatible with Cas12b (**Supplementary Figure 7**).

We next assessed whether the widely used type II enzyme, Cas9, is suitable for REDRAW. We replaced Cas12a in RE2 with nuclease-active SpCas9 to generate SpCas9-RE2. TagRNAs were designed for SpCas9-RE2 by attaching PBS and RTT sequences and the PP7 end-stabilization motif at the 3’ end of SpCas9 sgRNA. After transfecting plasmids expressing SpCas9-RE2 and end-stabilized, Cas9-compatible tagRNAs into HEK293T cells, we observed up to 5.3% precise editing across three different genomic loci (**Figure 6a**). In the same experiment we tested whether stagRNAs can be used with Prime editor 2 (PE2), and whether enhanced pegRNAs (ePegRNAs^15^), developed for use in Prime editing, can be used with SpCas9-RE2. We tested these configurations at three loci in HEK293T cells for their ability to generate double transversion mutations with various forms of tagRNAs stabilized with either PP7 as used in RE2, or evoPreQ1 as used in Prime editing, and ePegRNAs^15^. Consistent with the presumed mechanisms of Prime editing and REDRAW, both tagRNAs and ePegRNAs were compatible with Cas9-RE2, whereas PE2 was only compatible with ePegRNAs. No precise editing was detected in the absence of RT, or when the crRNA did not encode any RNA extensions.

**Figure 6.**
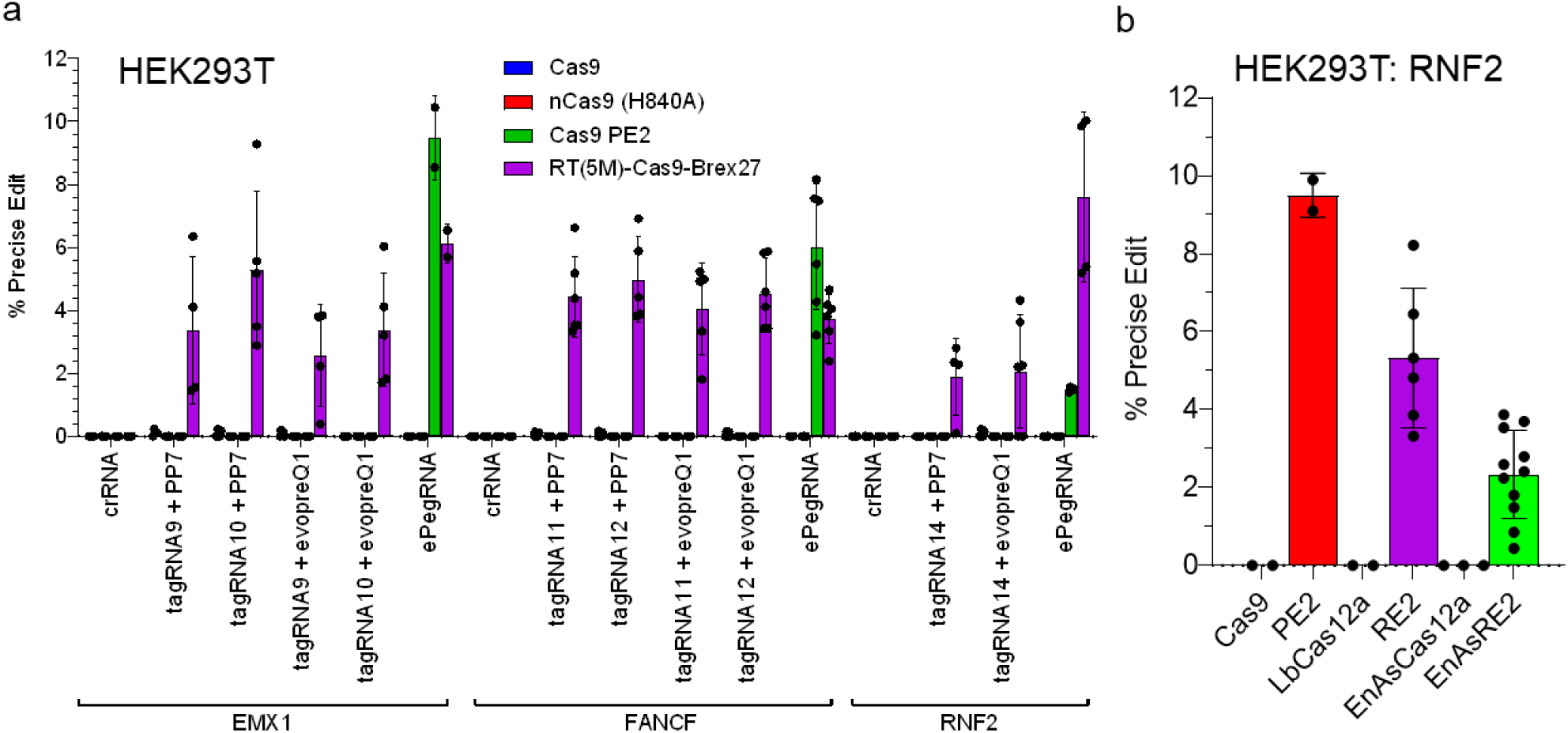
REDRAW with Cas9, and comparison with Prime editing. (a) Cas9, Cas9 (H840A), PE2, and Cas9-RE2 was used with a guide RNA without any RNA extension, tagRNA stabilized with PP7 hairpin, tagRNA stabilized with evoPreQ1, and ePegRNA encoding the same edit at three genomic loci. Cas9-RE2 can utilize tagRNA as well as ePegRNA, but PE2 is only compatible with ePegRNA. Cas9 and Cas9 H840A nickase without RT(5M) cannot mediate precise editing. (b) Side-by-side comparison of REDRAW (RE2 and EnAsRE2) and Prime editing (PE2) targeting the RNF2 genomic locus in HEK293T cells.

Although a comparison of Cas12a-REDRAW and Prime editors is confounded due to their orthogonal PAM specificities and editing windows, we targeted the same genic region (RNF2) using RE2, EnAs-RE2, and PE2 in HEK293T cells and quantified the precise editing efficiency using high-throughput sequencing. Consistent with the published activity profile^5^, PE2 generated precise edits at ~9.5% efficiency, which was slightly higher than that of RE2 (5.3%) and EnAsRE2 (2.3%) (**Figure 6b**).

In summary, REDRAW is compatible with Cas12b and Cas9, and can accommodate various RNA templates. Unlike Prime editors, we expect REDRAW to be compatible with a wide range of CRISPR subtypes because it is agnostic with respect to the accessibility of the R-loop DNA. We similarly predict any site-specific double-stranded nuclease could be used in place of a CRISPR protein.

### REDRAW compares favorably to homology-directed repair

Homology-directed repair (HDR) can introduce precise base substitutions and insertions by repairing a nuclease-induced, DSB via the homologous recombination pathway with edits encoded on a DNA template^24^. We compared the efficiency of REDRAW to that of Cas12a-mediated HDR in HEK293T cells. We transfected RE2 and stagRNAs encoding four different double transversion mutations in parallel with Cas12a nuclease and crRNAs mixed with single-stranded, 100-base DNA oligonucleotides encoding the same edits in both target strand and NTS orientations. RE2 caused up to 5.7% of precise editing, whereas HDR using Cas12a generated maximum editing of 0.9% (**Supplementary Figure 8a**). On average, RE2 was 6.1-fold more efficient than Cas12a-mediated HDR. We observed that REDRAW has a repressive effect on indel frequency, causing about 65% fewer indels at the same target compared to a typical Cas12a nuclease (**Supplementary Figure 9**). Furthermore, the edit-to-indel ratio for RE2 was 9.3- and 10.6-fold greater than that of HDR using target strand and NTS ssDNA donors, respectively, across all four DNA modifications tested (**Supplementary Figure 8b**). This shows REDRAW offers a substantial improvement in efficiency and product purity when compared to Cas12a-mediated HDR methods. However, because REDRAW uses Cas12a as a nuclease, indels still comprise 5-25% of total REDRAW edits, marking indel reduction as an obvious target for future engineering to improve product purity.

### Off-target activity of REDRAW

Previous reports have shown that the Cas12a nuclease exhibits markedly less off-target DNA cleavage than the Cas9 nuclease^25^, suggesting REDRAW may have favorable target specificity. We investigated off-target editing of REDRAW at two loci that were previously identified for Cas12a using GUIDE-Seq^25,26^. We treated HEK293T cells with LbCas12a nuclease, RE2, RE3, RR-RE2 and RR-RE3. We determined the extent of off-target editing induced by these genome editors via high-throughput sequencing of DNA extracted from transfected HEK293T cells. RE2, RE3, RR-RE2, and RR-RE3 generated the intended precise edit at the on-target loci with appropriately designed stagRNAs (**Supplementary Figure 10a,b**). We did not observe any RNA-templated edits encoded by the stagRNAs at the off-target sites, presumably because of the lack of homology in the PBS and RTT regions within those loci. Compared to Cas12a, RE2 resulted in 74.0% and 60.0% reduced indel formation at off-target sites 1 and 2 using identical stagRNAs (**Supplementary Figure 10a**). RE3 led to a decrease in indel formation at the respective off-target sites by 74.8% and 68.7% averaged across both nicking crRNA usages. Similarly, RR-RE2 led to an average 13.5% and 59.0% reduction and RR-RE3 showed an average 46.3% and 48.4% reduction of indels created at off-target sites 1 and 2, respectively, using identical stagRNAs (**Supplementary Figure 10b**). Taken together, the Cas12a REDRAW editors generally show less RNA-guided off-target indel creation than Cas12a alone.

## DISCUSSION

### REDRAW is a novel reverse-transcriptase-based genome editing technology with a distinct mode of action

In REDRAW we have developed a reverse transcriptase editing system that complements base editing by enabling templated installation of single base transversions, multi-base substitutions, and precise indels. By reaching across a DSB to prime the target DNA strand, REDRAW has a flexible architecture and is therefore compatible with both type II and type V CRISPR systems. Thus, by extending beyond the genetic variation that can be introduced by base editors and by overcoming the limited CRISPR system compatibility of Prime editors, REDRAW is a highly functional tool that expands our ability to alter DNA sequences in living systems.

### Systematic optimization of REDRAW results in three generations of editors with increasing efficiency

In this study, REDRAW is extensively developed and characterized using Cas12a to deliver the nuclease and RT activities to the target site. REDRAW editor 1 (RE1) is an N-terminal fusion of RT(5M) with the LbCas12a nuclease. RE1 uses a crRNA extended with a sequence complementary to the PAM distal side of the DSB that enables reverse transcription of the RNA template sequence into the target-strand DNA. RE1 achieves up to 0.8% precise nucleotide modifications in human cells, validating the REDRAW approach and providing a springboard for further optimization.

REDRAW editor 2 (RE2) adds a C-terminal Brex27 peptide fusion to the RT(5M)-LbCas12a protein. Brex27 is a 38 aa peptide that recruits RAD51, a ssDNA binding protein^14^ native to all eukaryotic cells and important for chromosomal stability during DNA repair events. Brex27 increases REDRAW activity, consistent with a model where the DNA ends of the DSB are stabilized by RAD51, potentially extending their availability for REDRAW editing and reducing sequestration towards the unproductive NHEJ pathway. RE2 also includes a PP7 hairpin at the 3′ end of the tagRNA, reducing susceptibility of the extended sequence to 3′-5′ exonucleases. Together, the ssDNA stabilization and structured RNA motif stabilizing the tagRNA resulted in over 10-fold improvement in REDRAW activity, delivering up to 8.2% precise nucleotide modifications in human cells with RE2.

REDRAW editing creates a transitory intermediate with three features that require repair to resolve the target site disruption: 1) the reverse-transcribed sequence on the target strand invades the double helix across the DSB, creating a 5’ flap on the target strand that must be removed by cellular flap endonuclease 1 (FEN1); 2) this results in staggering of the DSB which must be repaired by cellular ligases; 3) the edited sequence contained within the reverse-transcribed bases is non-complementary to the NTS sequence it is paired with, creating a substrate for the cellular DNA mismatch repair pathway (MMR). Resolution of the DNA mismatch could result in the edited sequence of the target strand being corrected back to the native sequence, an outcome that would reverse the REDRAW edit. Thus, manipulation of the MMR pathway to favor incorporation of the edited sequence into the NTS would increase REDRAW efficiency.

The MMR pathway is stimulated by nicked DNA and preferentially repairs mismatches in the nicked strand to be complementary to the undamaged strand. REDRAW Editor 3 (RE3) exploits this tendency by using a secondary crRNA with specific mismatches to create a nick on the NTS DNA. crRNAs with certain mismatches to their target site form a structure that causes the Cas12a nuclease in the REDRAW protein to cut only one strand of the chromosome. In human cells, RE3 increases the efficiency of REDRAW by 1.4-fold compared to RE2.

### REDRAW works in multiple cell lines with a wide variety of CRISPR systems and has lower off-target activity than Cas12a

REDRAW editing is effective in different human cell lines including U2OS and HeLa, suggesting relevance to various *ex vivo* and *in vivo* therapeutic applications. It is also compatible with Cas12a from various species, with engineered variants thereof, with Cas12b and with Cas9 which provides access to a diverse range of PAM sequences and enables optimization of editing efficiency for specific cell types. For example, the highest human cell REDRAW editing (up to 12.2% precise modification) rate observed in this study used EnAs-RE2, which contains the enhanced AsCas12a^20^ that has been engineered for high efficiency in human cells. Replacing Cas12a in RE2 with nuclease-competent Cas12b and Cas9 led to the creation of Cas12b-RE2 and Cas9-RE2, respectively, which can each create precise edits in human cells.

As Cas12a is widely regarded to have less off-targeting than Cas9^25,27^, we expected REDRAW to have similarly low off target activity. In this study, RE2 and RE3 caused no RT-mediated precise sequence changes at two known Cas12a off-target sites. However, REDRAW did generate indels at these sites, albeit at substantially lower frequency than a Cas12a nuclease alone.

### REDRAW compared to Prime editing and homology-directed repair

REDRAW first incorporates the edit on the target strand DNA. This is a significant departure from the Prime editing paradigm, which reverse transcribes from the nicked NTS DNA. During Prime editing, a portion of the pegRNA is hybridized to the nicked non-target strand DNA^5^. The resulting complex is a substrate for RT-mediated DNA polymerization that transfers the RNA-encoded edit into genomic DNA. In contrast, the RNA template of REDRAW is hybridized to the target strand. Consequently, REDRAW does not depend on R-loop structure for successful reverse transcription and should be compatible with any CRISPR domain capable of inducing target strand cleavage. Thus, REDRAW demonstrates a new mode of reverse transcriptase editing with CRISPR systems.

CRISPR proteins, including type V enzymes, with PAMs adjacent to the 5′ end of the protospacer are configured to work better with REDRAW than with Prime editing. During REDRAW, DNA is synthesized towards the PAM located at the 5’ end of the protospacer from the site of DNA cleavage, introducing edits within the protospacer sequence or its PAM. Successful installation of the edit can prevent re-binding and re-cleavage of the target site, as the edited target sequence is no longer complementary to the crRNA. Conversely, extension of the non-target strand, as in Prime editing, would preserve most of a type V protospacer and PAM sequence, leaving the target sequence vulnerable to re-binding and re-cleavage after successful installation of the edit. This would increase the likelihood of further modifications to the correctly edited DNA, thereby destroying the intended allele.

Like Prime editors, REDRAW uses crRNA extended with PBS and RTT sequences required for reverse transcription. However, REDRAW is typically used with a longer range of PBS and RTT. Optimal PBS and RTT lengths are around 48 bp and 52 bp, respectively. REDRAW’s compatibility with relatively long template lengths suggests flexibility in the architecture of the target-bound enzyme and a mechanism that is not prone to steric interference by its components. We speculate this as a reason REDRAW activity is less dependent on the exact lengths of PBS and RTT than Prime editors, which are sensitive to length alterations as small as a single base in both PBS and RTT^5^.

The traditional approach to introduce base transversions, multi-base substitutions, or multi-base insertions into cells is oligo-templated repair via the HDR pathway. HDR requires a complex suite of cellular factors that are most abundant in actively dividing cells. The pathway comprises an imposing web of biological complexity and has been mostly intransigent to decades of efforts to improve its efficiency for genome editing. We found RE2 to be 6.1-fold more efficient, on average, than installation of the same edits by HDR with a Cas12a nuclease. Indel side products arise from NHEJ repair of a DSB and can therefore occur as byproducts of HDR and of REDRAW editing. By this measure, RE2 provides a substantially superior option to HDR with an edit-to-indel ratio up to 10.6-fold better than from oligo-templated HDR installing the same edits. However, indels still comprise 5-25% of the total target site changes arising from REDRAW activity, making further reduction of NHEJ-derived byproducts an important avenue for future research.

We have tested up to 24 base insertions with no decrease in REDRAW editing efficiency. We do not yet know the upper size limit of the reaction, but it is likely dependent on RT processivity, stagRNA integrity, and potentially other factors. For precise gene deletion, insertion, or replacement up to 24 bases and likely longer, REDRAW is an attractive alternative when using Cas12a or other type V CRISPR enzymes.

## Conclusion

REDRAW is a highly functional editing tool capable of installing all types of single and multiple base substitutions and small, precise indels. Its inherent architectural flexibility enables use with a wide variety of CRISPR enzymes including multiple type II and type V systems. We expect its novel mode of action, compatibility with diverse CRISPR/Cas families, and forgiving design principles will enable broad biomedical, biotechnological and agricultural applications.

## METHODS

### Cloning

Plasmids containing REDRAW constructs and guide RNA were either ordered and prepared by Genscript or cloned in house utilizing the Bioxp^™^. For Bioxp cloning fragment inserts were generated by CodexDNA and ran on the Bioxp^™^ for high throughput Gibson assembly. Following Synthesis with the Bioxp^™^, DNA was transformed into NEB 5-alpha cells (New England Biolabs) and plated on LB plates with carbenicillin for selection purposes and incubated overnight at 37 °C. Isolated colonies were selected and grown in LB liquid media supplemented with carbenicillin. DNA extraction was performed using a Qiagen miniprep kit and eluted in H_2_O before being sent to Azenta for sequence verification.

### Cell culture

HEK-293T (ATCC-CRL-3216^™^), U2OS (ATCC – HTB-96), and HeLa (GenTarget Inc, SC031-Puro) cells were cultured at 37 °C in a 5% CO_2_ incubator in T75 or T175 flasks with DMEM high glucose GlutaMAX^™^ media (ThermoFisher Scientific) supplemented with 10% FBS (Gibco) and Penicillin-Streptomycin (Gibco). Cells were passaged at 80% confluency and used until passage 20.

### Transfection

For transfection, cells were counted using trypan blue and Countess II FL cell counter (ThermoFisher Scientific) and seeded into 48 well plates at 55,000 cells per well (HEK-293T), 35,000 cells per well (U2OS), and 25,000 cells per well (HeLa). All cells were seeded one day prior to transfection in 250 μL of DMEM media with 10% FBS. Lipofectamine LTX (Invitrogen), TransIT-HelaMonster (Mirus) and MirusIT 2020 (Mirus) were used to transfect HEK293T, HeLa, and U2OS cells, respectively. All cell types were transfected with 500 ng of total DNA. In two plasmid transfection experiments the DNA ratio of protein to guide plasmid was 3:1. In two protein plus guide transfections, the DNA ratio was 2:1:1. Seventy two hours after transfection, media was removed and cells were lysed in 10 mM Tris-HCl (Invitrogen) 0.05% SDS (Invitrogen) with Protein Kinase K (NEB) and incubated at 37 °C for 1 h. Cell lysate was incubated at 95 °C for 15 min then spun down at 4000 rpm for 15 min. Cell lysate was then diluted at a 1:1 ratio with dH_2_O to be later used for high throughput sequencing.

### High throughput DNA sequencing of genomic DNA samples

Cell lysate was PCR amplified with genomic primers flanked by Illumina adaptor overhang to generate approximately 300 base products. The products were amplified with another round of PCR and pooled together to generate Illumina barcoded library. The library was cleaned up using Sera-Mag Select (Cytiva) and quantified by SpectraMax^®^ Quant^™^ dsDNA Assay Kit (Molecular Devices). Pooled amplicon libraries were sequenced using Illumina MiSeq 2×250 kit.

### Data analysis

Raw Illumina MiSeq paired-end reads were trimmed, merged, and aligned using CRISPResso2 (Ref^28^). Trimmomatic^29^ was used to trim Nextera paired-end adapters from each read-pair (“ILLUMINACLIP:NexteraPE-PE.fa:0:90:10:0:true”) and then reads were filtered with an average base quality < 20. Next, read pairs were merged using FLASH^30^ requiring a minimum overlap of 20 bases and a maximum overlap of 260 bases. Finally, reads were aligned with a modified Needleman-Wunsch algorithm in CRISPResso2 and sequences were kept only for those with a score ≥ 60. Edit quantification was done using the base editor mode and a quantification window of 25 bases upstream and downstream of the predicted cleavage site. To ensure enough coverage to capture rare events, samples had at least 5,000x coverage per target amplicon. To minimize false positives, edit events with < 15 supporting reads or an edited read percentage < 0.1% were discounted. To identify expected REDRAW edits, RTT sequence was aligned to each amplicon, using Needleman-Wunsch alignment algorithm, and these variants were treated as the expected edit during analysis.

### Protein purification

All chemicals were purchased from Millipore Sigma unless otherwise noted. pET28a LbCas12a and LbCas12a (R1138A) possessing a C-terminal 6x Histidine tag were transformed into Rosetta 2 (DE3) cells. For each protein, 500 mL of LB media with kanamycin and chloramphenicol was inoculated with 1 mL of overnight starter culture and grown until OD600 of 0.4-0.6. Protein expression was induced by addition of 0.2 mM IPTG and cells were allowed to express for 18-20 h at 18 °C with 2255 rpm shaking. Cell pellets were harvested by centrifugation and stored at −80 °C. The cell pellet was resuspended in 10 mL lysis buffer: 50 mM HEPES-KOH pH 7.5, 0.5 M NaCl, 10% glycerol, 5 mM TCEP (Pierce), 1X Protease Inhibitor Cocktail III, 5 U/mL DNase I (Pierce 89836) and 10 mM Imidazole. The cells were lysed by sonication 5×10 s bursts with 30 s rest at 50% amplitude. Clarified supernatant was passed over a ~3 mL-Ni-NTA column (HisPur^™^ resin, Pierce) equilibrated in buffer A1: 20 mM HEPES pH 7.5, 500 mM NaCl, 10% glycerol, 2 mM TCEP, and 10 mM imidazole. The column was washed in buffer A1 for 4 cv. A second wash step of 3 cv in Buffer A2: 20 mM HEPES pH 7.5, 500 mM NaCl, 10% glycerol, 2 mM TCEP, and 20 mM imidazole pH 7.5 was performed. Protein was eluted from the column in elution buffer: 20 mM HEPES pH 7.5, 150 mM NaCl, 10% glycerol, 0.5 mM TCEP, and 350 mM Imidazole pH 7.5. A second purification step utilizing cation exchange chromatography was performed. Briefly, the peak fractions from the Ni^2+^ were pooled, and loaded onto a 1 mL MonoS column using AKTA Purifier (Cytiva). The column was equilibrated in Buffer A3: 20 mM HEPES-KOH pH 7.5, 150 mM NaCl, 10% glycerol, 1 mM MgCl_2_, and 0.5 mM TCEP. After sample loading, the column was washed 5 CV in buffer A3 followed by elution of the protein over a linear gradient 15 cv up to 100% Buffer B1: 20 mM HEPES-KOH pH 7.5, 1 M NaCl, 10% glycerol, 1 mM MgCl2, and 0.5 mM TCEP. Peak fractions were identified concentrated by Amicon 50K MWCO and brought up to 50% glycerol for final storage at −20 °C.

### In vitro nickase assay

Plasmid containing DMNT1 target region was ordered from Twist Biosciences. Plasmid was purified using QIAprep Spin Miniprep kit (Qiagen) according to manufacturer’s instructions. Ten LbCas12a crRNAs were designed targeting DNMT1. One crRNA served as the cutting control with no mismatches in the spacer sequence. The other cRNAs contained base mismatches at various positions within the spacer sequence (converting C to A, T to G, A to C, and G to T). RNA guides were ordered from Synthego and rehydrated with nuclease free water to a final concentration of 300 nM. Plasmid nicking assays were carried out in 27 μL reactions containing NEB2.1, LbCas12a (1 μM), nuclease free water, and crRNA (300 nM). The nickase control contained an unmodified guide with Cas12a (R1138A) instead of LbCas12a. Samples were incubated at room temperature for 10 min then 3 μL of target site plasmid (50 ng/μL) was added to each sample. Samples were incubated for 15 min at 37 °C then heat inactivated at 80 °C for 20 min. To each sample, 6 μL of 6x purple loading dye was added and 20 μL of total sample was loaded into a 1% agarose gel then run for 1 h at 120 V. Gel was imaged using the Azure C600 gel imager.

## Authorship contributions

Y.B.K. conceived, designed, directed, conducted experiments, and performed analyses. E.B.P., M.B., E.S.P., B.A.P., and G.M.R. cloned plasmids, conducted experiments, and performed analyses. D.N. conducted experiments. E.B.P. performed in vitro nickase assays. D.G.S. supervised and designed experiments. D.S. and S.J. performed high throughput sequencing. J.F. and Z.C. provided high throughput sequencing data analysis. J.E.H. performed protein purifications. Y.B.K., J.M.W., and A.H. supervised research and wrote the manuscript.

