## Supplemental Information for "A novel mechanistic framework for precise sequence replacement using reverse transcriptase and diverse CRISPR-Cas systems"

### SUPPLEMENTARY FIGURES

a

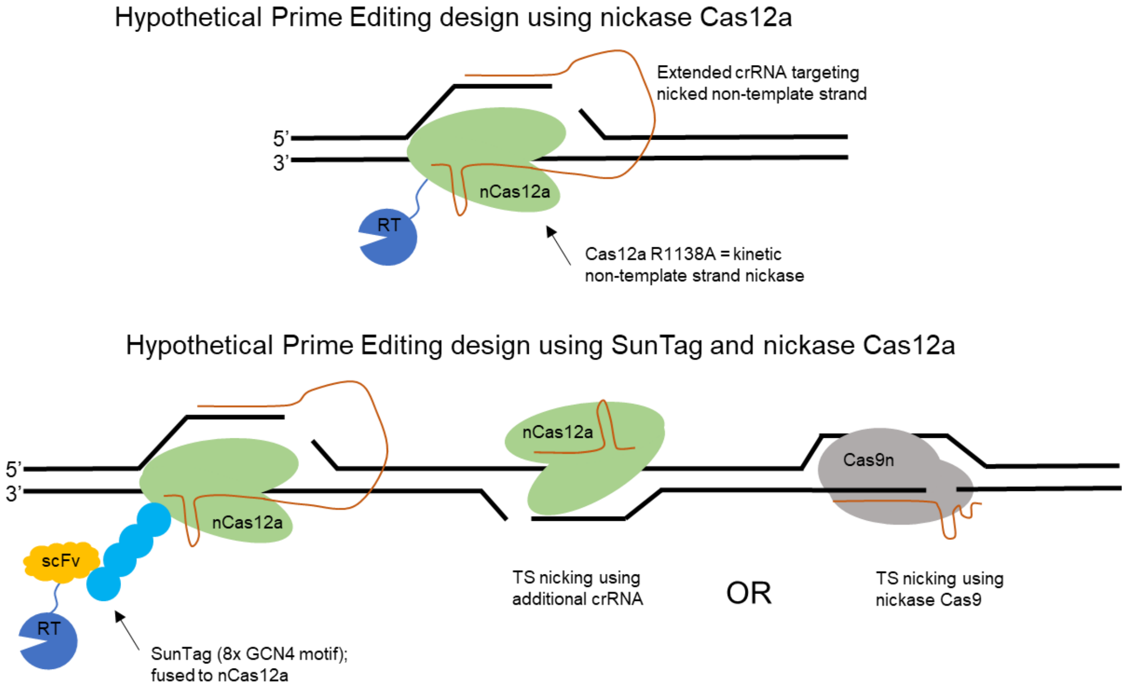
Hypothetical Prime Editing design using nickase Cas12a
Extended crRNA targeting nicked non-template strand
5'
3'
nCas12a
RT
Cas12a R1138A = kinetic non-template strand nickase
Hypothetical Prime Editing design using SunTag and nickase Cas12a
nCas12a
scFv
RT
SunTag (8x GCN4 motif); fused to nCas12a
TS nicking using additional crRNA
OR
Cas9n
TS nicking using nickase Cas9

b

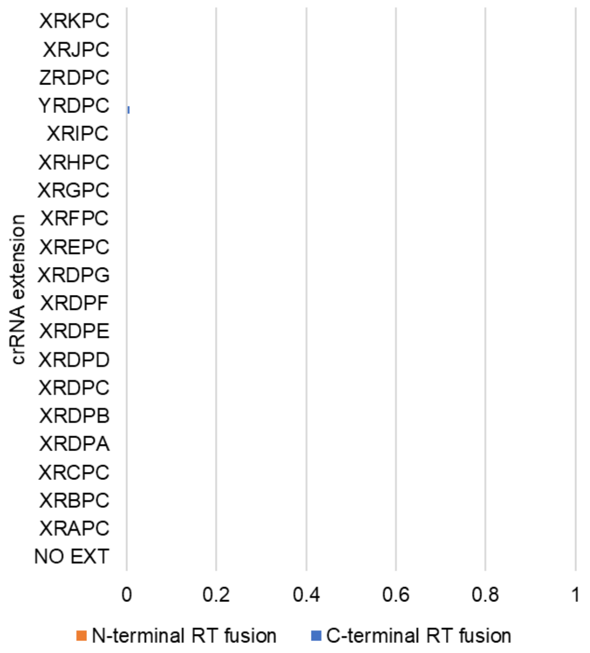
crRNA extension
XRKPC
XRJPC
ZRDPC
YRDPC
XRIPC
XRHPC
XRGPC
XRFPC
XREPC
XRDPG
XRDPF
XRDPE
XRDPD
XRDPC
XRDPB
XRDPA
XRCPC
XRBPC
XRAPC
NO EXT
0
0.2
0.4
0.6
0.8
1
N-terminal RT fusion
C-terminal RT fusion

809

c

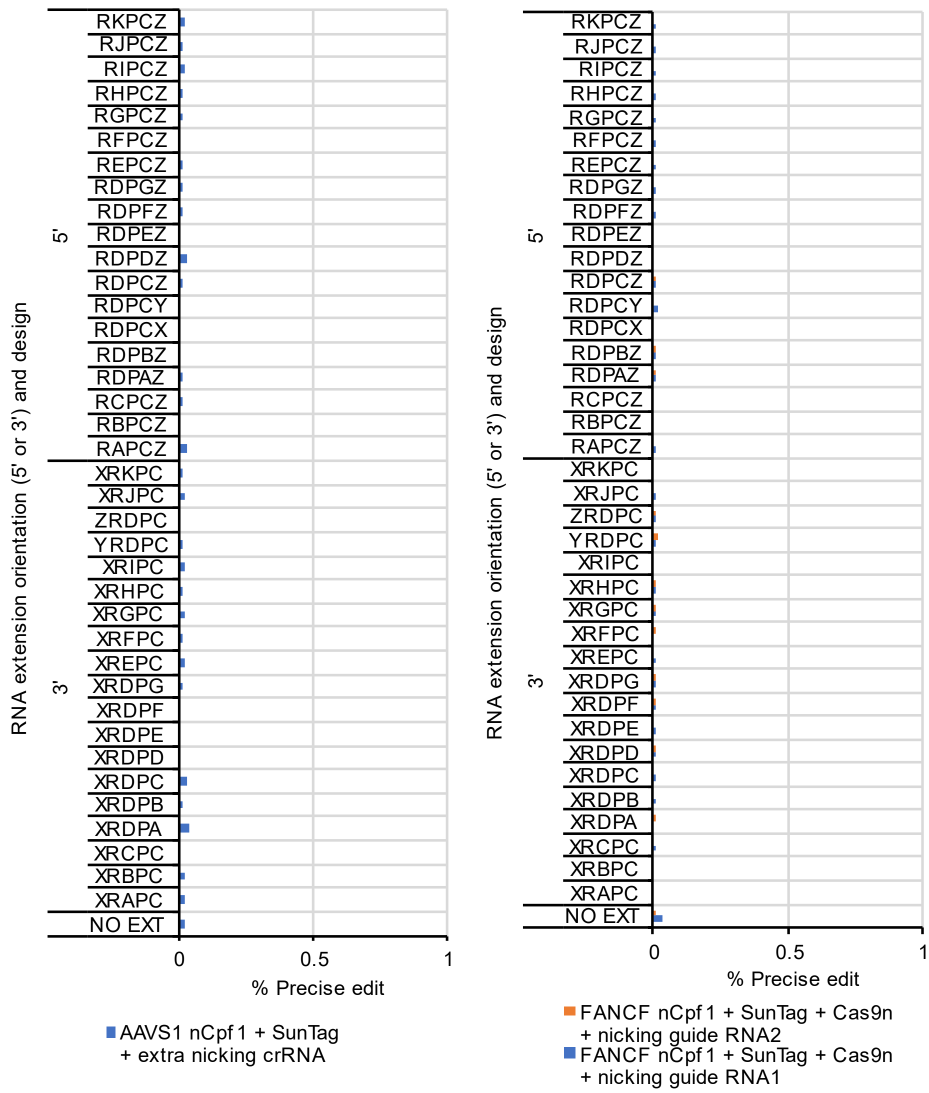
RKPCZ
RJPCZ
RIPCZ
RHPCZ
RGPCZ
RFPCZ
REPCZ
RDPGZ
RDPFZ
RDPEZ
RDPDZ
RDPCZ
RDPCY
RDPCX
RDPBZ
RDPAZ
RCPCZ
RBPCZ
RAPCZ
XRKPC
XRJPC
ZRDPC
YRDPC
XRIPC
XRHPC
XRGPC
XRFPC
XREPC
XRDPG
XRDPF
XRDPE
XRDPD
XRDPC
XRDPB
XRDPA
XRCPC
XRBPC
XRAPC
NO EXT
5'
3'
RNA extension orientation (5' or 3') and design
0
0.5
1
% Precise edit
AAVS1 nCpf1 + SunTag + extra nicking crRNA
FANCF nCpf1 + SunTag + Cas9n + nicking guide RNA2
FANCF nCpf1 + SunTag + Cas9n + nicking guide RNA1

d

| Extension code | extended RNA sequence |
|---|---|
| XRDPA | TCTGTCCCCTCCACCCCACAGTGCTAGTGGCCACACTGTGGGGTGG |
| XRDPB | TCTGTCCCCTCCACCCCACAGTGCTAGTGGCCACACTGTGGGGTGGAG |
| XRDPC | TCTGTCCCCTCCACCCCACAGTGCTAGTGGCCACACTGTGGGGTGGAGGG |
| XRDPD | TCTGTCCCCTCCACCCCACAGTGCTAGTGGCCACACTGTGGGGTGGAGGGGA |
| XRDPE | TCTGTCCCCTCCACCCCACAGTGCTAGTGGCCACACTGTGGGGTGGAGGGGA |
| XRDPF | TCTGTCCCCTCCACCCCACAGTGCTAGTGGCCACACTGTGGGGTGGAGGGGA<br>GA |
| XRDPG | TCTGTCCCCTCCACCCCACAGTGCTAGTGGCCACACTGTGGGGTGGAGGGGA<br>GATA |
| XRAPC | TCTGTCCCCTCCACCCCACAGTGGCCACACTGTGGGGTGGAGGG |
| XRBPC | TCTGTCCCCTCCACCCCACAGTGTGGCCACACTGTGGGGTGGAGGG |
| XRCPC | TCTGTCCCCTCCACCCCACAGTGAGTGGCCACACTGTGGGGTGGAGGG |
| XREPC | TCTGTCCCCTCCACCCCACAGTGCCCTAGTGGCCACACTGTGGGGTGGAGGG |
| XRFPC | TCTGTCCCCTCCACCCCACAGTGGTCCCTAGTGGCCACACTGTGGGGTGGAG |
| XRGPC | TCTGTCCCCTCCACCCCACAGTGCTGTCCCTAGTGGCCACACTGTGGGGTGGA<br>GG |
| XRHPC | TCTGTCCCCTCCACCCCACAGTGTCCTGTCCCTAGTGGCCACACTGTGGGGTG<br>AGGG |
| XRIPC | TCTGTCCCCTCCACCCCACAGTGAATCCTGTCCCTAGTGGCCACACTGTGGGG<br>GGAGGG |
| YRDPC | TCTGTCCCCTCCACCCCACAGTGCACCCTAGTGGCCACACTGTGGGGTGGAG |
| ZRDPC | TCTGTCCCCTCCACCCCACAGTGCACCCACCCACCCTAGTGGCCACACTGTGGG<br>GTGGAGGG |
| XRJPC | TCTGTCCCCTCCACCCCACAGTGCTAGTGGCCTCACTGTGGGGTGGAGGG |
| XRKPC | TCTGTCCCCTCCACCCCACAGTGCTAGTGGCCGCACTGTGGGGTGGAGGG |
| RDPCZ | CTAGTGGCCACACTGTGGGGTGGAGGGCACCCACCCACC |
| RDPCX | CTAGTGGCCACACTGTGGGGTGGAGGG |
| RDPCY | CTAGTGGCCACACTGTGGGGTGGAGGGCACC |
| RDPAZ | CTAGTGGCCACACTGTGGGGTGGCACCCACCCACC |
| RDPBZ | CTAGTGGCCACACTGTGGGGTGGAGCACCCACCCACC |
| RDPDZ | CTAGTGGCCACACTGTGGGGTGGAGGGGACACCCACCCACC |
| RDPEZ | CTAGTGGCCACACTGTGGGGTGGAGGGGACACACCCACCCACC |
| RDPFZ | CTAGTGGCCACACTGTGGGGTGGAGGGGACAGACACCCACCCACC |
| RDPGZ | CTAGTGGCCACACTGTGGGGTGGAGGGGACAGATACACCCACCCACC |
| RAPCZ | GCCACACTGTGGGGTGGAGGGCACCCACCCACC |
| RBPCZ | TGGCCACACTGTGGGGTGGAGGGCACCCACCCACC |
| RCPCZ | AGTGGCCACACTGTGGGGTGGAGGGCACCCACCCACC |
| REPCZ | CCCTAGTGGCCACACTGTGGGGTGGAGGGCACCCACCCACC |
| RFPCZ | GTCCCTAGTGGCCACACTGTGGGGTGGAGGGCACCCACCCACC |
| RGPCZ | CTGTCCCTAGTGGCCACACTGTGGGGTGGAGGGCACCCACCCACC |
| RHPCZ | TCCTGTCCCTAGTGGCCACACTGTGGGGTGGAGGGCACCCACCCACC |
| RIPCZ | AATCCTGTCCCTAGTGGCCACACTGTGGGGTGGAGGGCACCCACCCACC |
| RJPCZ | CTAGTGGCCTCACTGTGGGGTGGAGGGCACCCACCCACC |

| RTT edit | CODE | Length | RTT sequence |
|---|---|---|---|
| 1GtoT | RA | 8 | GCCACACT |
| 1GtoT | RB | 10 | TGGCCACACT |
| 1GtoT | RC | 12 | AGTGGCCACACT |
| 1GtoT | RD | 14 | CTAGTGGCCACACT |
| 1GtoT | RE | 16 | CCCTAGTGGCCACACT |
| 1GtoT | RF | 18 | GTCCCTAGTGGCCACACT |
| 1GtoT | RG | 20 | CTGTCCCTAGTGGCCACACT |
| 1GtoT | RH | 22 | TCCTGTCCCTAGTGGCCACACT |
| 1GtoT | RI | 24 | AATCCTGTCCCTAGTGGCCACACT |
| 1GtoA | RJ | 14 | CTAGTGGCCTCACT |
| 1GtoC | RK | 14 | CTAGTGGCCGCACT |
| 1GtoT | RL | 40 | GGGCTTTTCTGTCACCAATCCTGTCCCTAGTGGCCA<br>ACT |
| 1GtoT | RM | 50 | CCTAAGGATGGGGCTTTTCTGTCACCAATCCTGTCCC<br>TAGTGGCCACACT |
| 1GtoT | RN | 60 | AAGGAGGAGGCCTAAGGATGGGGCTTTTCTGTCACC<br>AATCCTGTCCCTAGTGGCCACACT |
| 1GtoT | RO | 70 | GGAGACTAGGAAGGAGGAGGCCTAAGGATGGGGCTT<br>TTCTGTCACCAATCCTGTCCCTAGTGGCCACACT |
| 1GtoT | RP | 80 | CCCAATATCAGGAGACTAGGAAGGAGGAGGCCTAAG<br>GATGGGGCTTTTCTGTCACCAATCCTGTCCCTAGTGG<br>CCACACT |

| | CODE | length | PBS sequence |
|---|---|---|---|
| | PA | 9 | GTGGGGTGG |
| | PB | 11 | GTGGGGTGGAG |
| | PC | 13 | GTGGGGTGGAGGG |
| | PD | 15 | GTGGGGTGGAGGGGA |
| | PE | 17 | GTGGGGTGGAGGGGACA |
| | PF | 19 | GTGGGGTGGAGGGGACAGA |
| | PG | 21 | GTGGGGTGGAGGGGACAGATA |
| | PH | 23 | GTGGGGTGGAGGGGACAGATAAA |
| | PI | 25 | GTGGGGTGGAGGGGACAGATAAAAG |
| | PJ | 27 | GTGGGGTGGAGGGGACAGATAAAAGTA |
| | PK | 29 | GTGGGGTGGAGGGGACAGATAAAAGTACC |
| | PL | 39 | GTGGGGTGGAGGGGACAGATAAAAGTACCCAGAACC<br>AGA |
| | PM | 49 | GTGGGGTGGAGGGGACAGATAAAAGTACCCAGAACC<br>AGAGCCACATTAA |
| | PN | 59 | GTGGGGTGGAGGGGACAGATAAAAGTACCCAGAACC<br>AGAGCCACATTAACCGGCCCTGG |
| | PO | 69 | GTGGGGTGGAGGGGACAGATAAAAGTACCCAGAACC<br>AGAGCCACATTAACCGGCCCTGGGAATATAAGG |
| | PP | 79 | GTGGGGTGGAGGGGACAGATAAAAGTACCCAGAACC<br>AGAGCCACATTAACCGGCCCTGGGAATATAAGGTGGT<br>CCCAGC |

| | | CODE | linker sequence |
|---|---|---|---|
| | | X | |
| | | Y | CACC |
| | | Z | CACCCACCCACC |

**Supplementary Figure 1. Prime editing architecture is not compatible with Cas12a.** (a) In a hypothetical Cas12a Prime editing scenario, non-target strand nicking Cas12a (R1138A) is used with RT(5M). The RTT and PBS sequences are designed to be complementary to the non-target strand DNA. Prime Editing configurations using both fusion and SunTag recruitment strategies were explored, with an additional nickase module in some scenarios. Extensive testing of (b) various lengths of RTT and PBS using N-terminal and C-terminal RT(5M) fusions and (c) antibody (SunTag) recruitment of RT(5M) in the presence of secondary nicking guide RNA failed to detect significant precise editing. (d) Extended RNA sequences used in this figure.

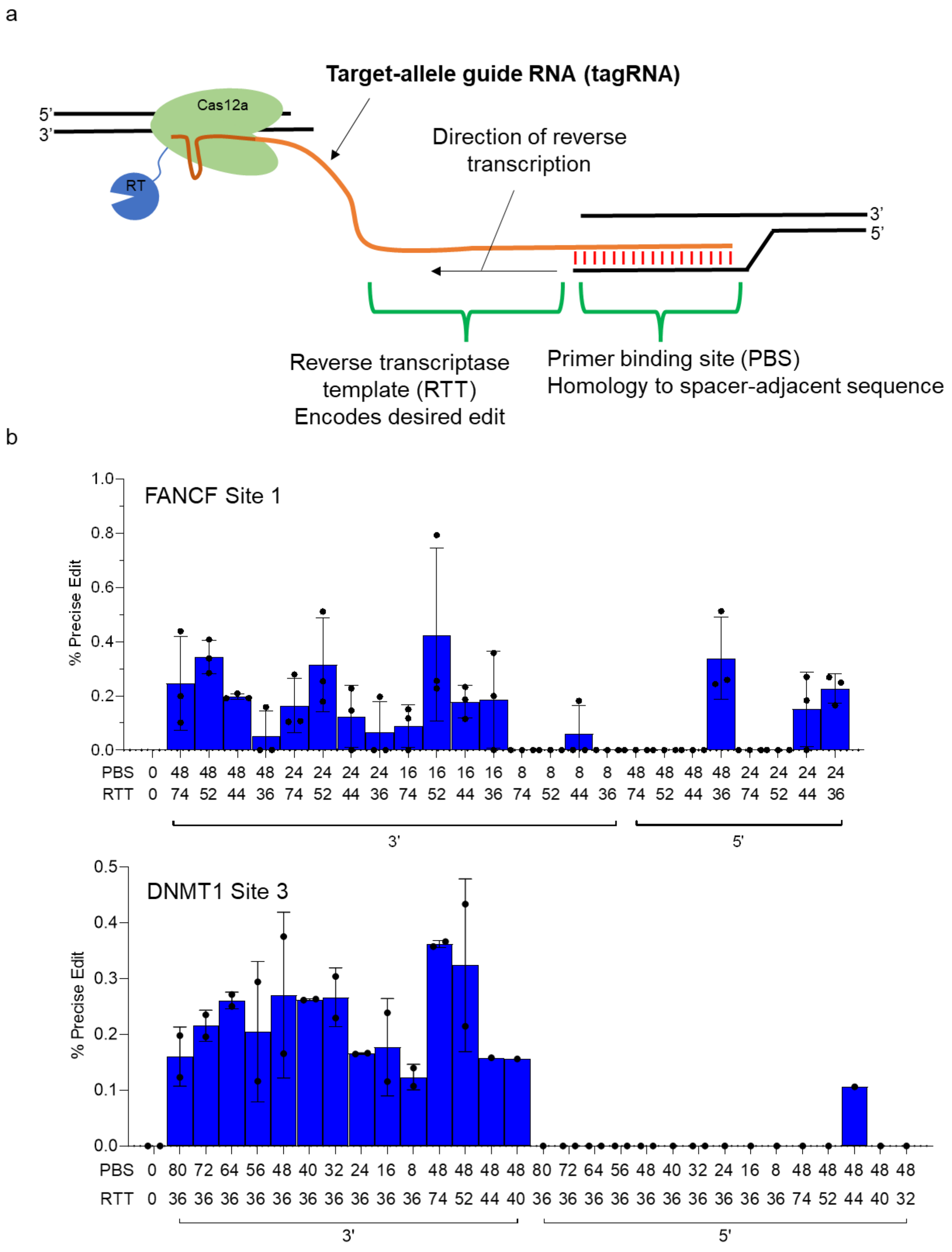
a
Target-allele guide RNA (tagRNA)
Cas12a
RT
5'
3'
3'
5'
Direction of reverse transcription
Reverse transcriptase template (RTT)
Encodes desired edit
Primer binding site (PBS)
Homology to spacer-adjacent sequence
b
FANCF Site 1
% Precise Edit
PBS 0 48 48 48 48 24 24 24 24 16 16 16 16 8 8 8 8 48 48 48 48 24 24 24 24
RTT 0 74 52 44 36 74 52 44 36 74 52 44 36 74 52 44 36 74 52 44 36 74 52 44 36
3'
5'
DNMT1 Site 3
% Precise Edit
PBS 0 80 72 64 56 48 40 32 24 16 8 48 48 48 48 80 72 64 56 48 40 32 24 16 8 48 48 48 48 48
RTT 0 36 36 36 36 36 36 36 36 36 36 74 52 44 40 36 36 36 36 36 36 36 36 36 36 74 52 44 40 32
3'
5'

**Supplementary Figure 2. Features of REDRAW tagRNA.** (a) In REDRAW, the extended crRNA containing an RT template (RTT) and primer binding site (PBS) is referred to as target-allele guide RNA (tagRNA). PBS hybridizes to the target strand DNA and RTT is used by the RT(5M) to encode the desired edit into DNA. (b) Optimal PBS and RTT lengths are identified by screening tagRNAs in HEK293T cells.

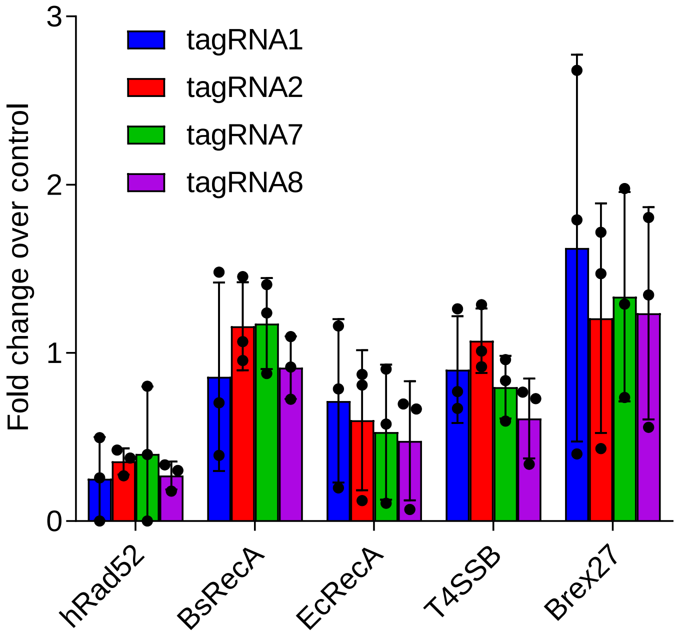


**Supplementary Figure 3. Brex27 peptide fusion at C-terminus of RE1.** Across 4 different tagRNAs tested, C-terminal Brex27 fusion resulted in on average 1.3-fold increase in efficiency over RE1.

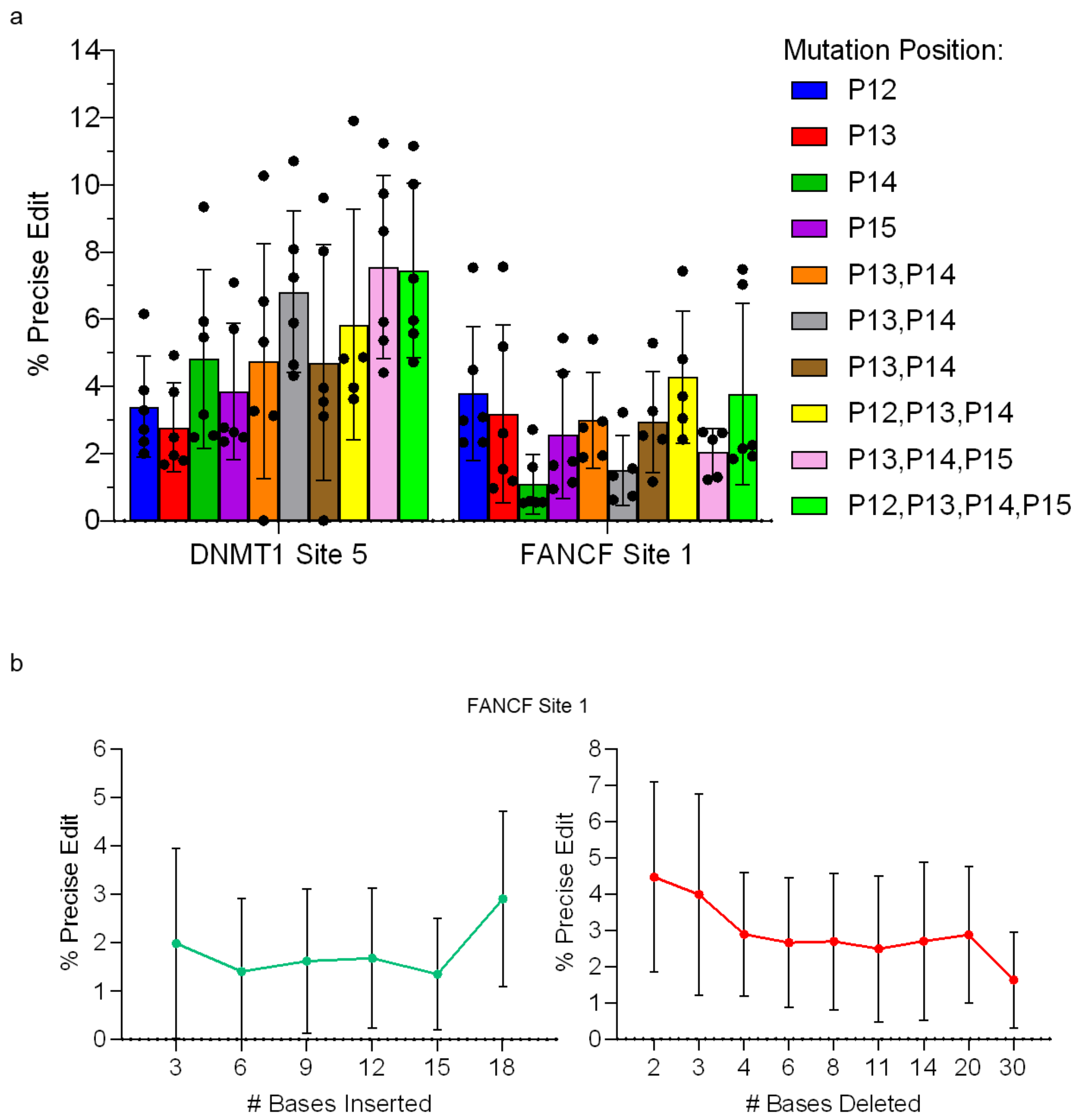


**Supplementary Figure 4. Characterization of mutational scope of REDRAW.** (a) REDRAW can generate single, double, triple, and quadruple mutations at 2 genomic sites. (b) Insertions and deletions mediated by RE2 at a second locus.

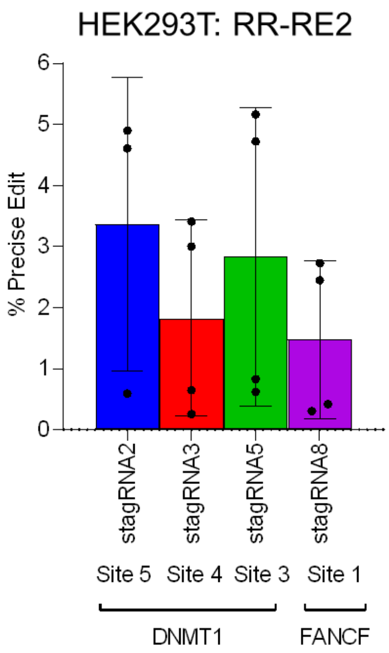


**Supplementary Figure 5. REDRAW with RR-LbCas12a**. RR-RE2 is constructed by replacing LbCas12a in RE2 with RR-LbCas12a, which has two arginine mutations that expands its PAM specificity. RR-RE2 mediates efficient precise editing at four different loci in HEK293T cells.

a

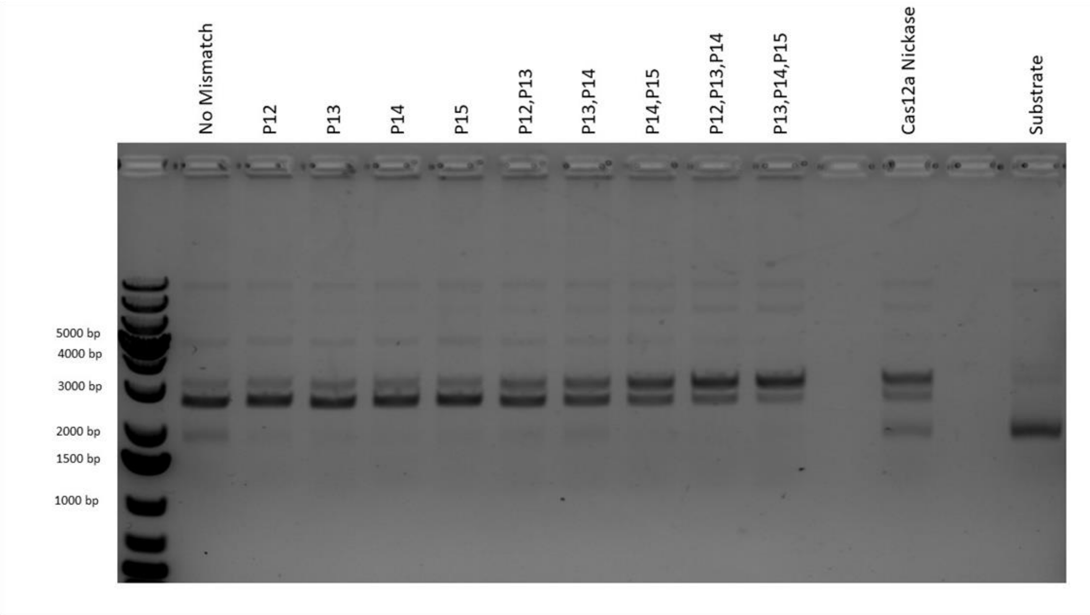


b

| | tagRNA15 | tagRNA17 |
|---|---|---|
| No nicking crRNA | 0.69 | 0.51 |
| crRNA; full complementarity | 0.00 | 0.00 |
| crRNA; mismatch at P12 | 0.00 | 4.62 |
| crRNA; mismatch at P13 | 0.00 | 0.00 |
| crRNA; mismatch at P14 | 0.00 | 0.00 |
| crRNA; mismatch at P15 | 0.00 | 0.36 |
| crRNA; mismatch at P12, 13 | 3.77 | 4.46 |
| crRNA; mismatch at P13, 14 | 0.15 | 1.08 |
| crRNA; mismatch at P12, 13, 14 | 0.68 | 0.98 |
| crRNA; mismatch at P13, 14, 15 | 0.00 | 0.74 |

**Supplementary Figure 6. Enhancing REDRAW by providing a nick on the DNA strand opposite to edit-containing strand.** (a) In vitro supercoiled plasmid nicking assay. Nuclease-active Cas12a can create single strand nicks by providing crRNAs with mismatches in position 12, 13, 14, or 15. Positions are indicated as P, followed by a numeral. crRNAs with no mismatches (control), single mismatches at positions 12-15, double consecutive mutations at positions 12-15, and triple consecutive mutations at positions 12-15 were tested. Cas12a nickase (R1138A) with fully matched crRNA is used as a positive control for creating a nicked plasmid. (b) Secondary mismatch containing crRNAs increase precise editing efficiency in HEK293T cells. In this target site, mismatches at position 12 and 13 led to the most significant fold improvement.

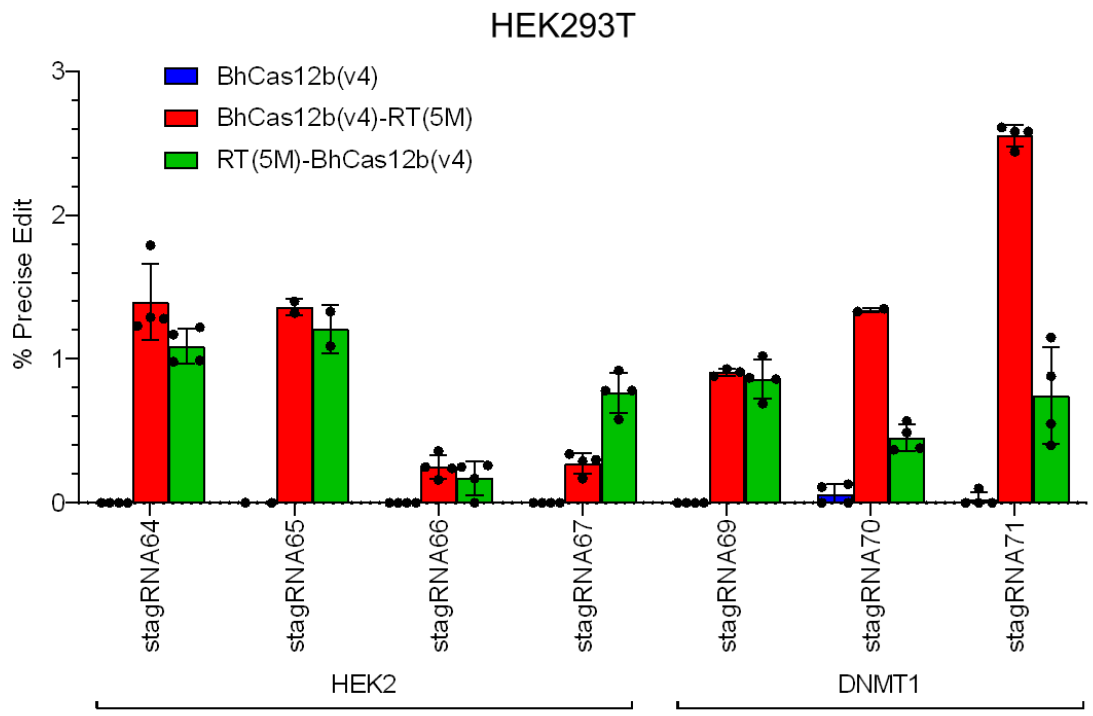


**Supplementary Figure 7. REDRAW with BhCas12b.** BhCas12b is also compatible with REDRAW and generates detectable precise editing using both N-terminal and C-terminal RT(5M) fusion to BhCas12b v4, which is a BhCas12b engineered for higher activity in human cells.

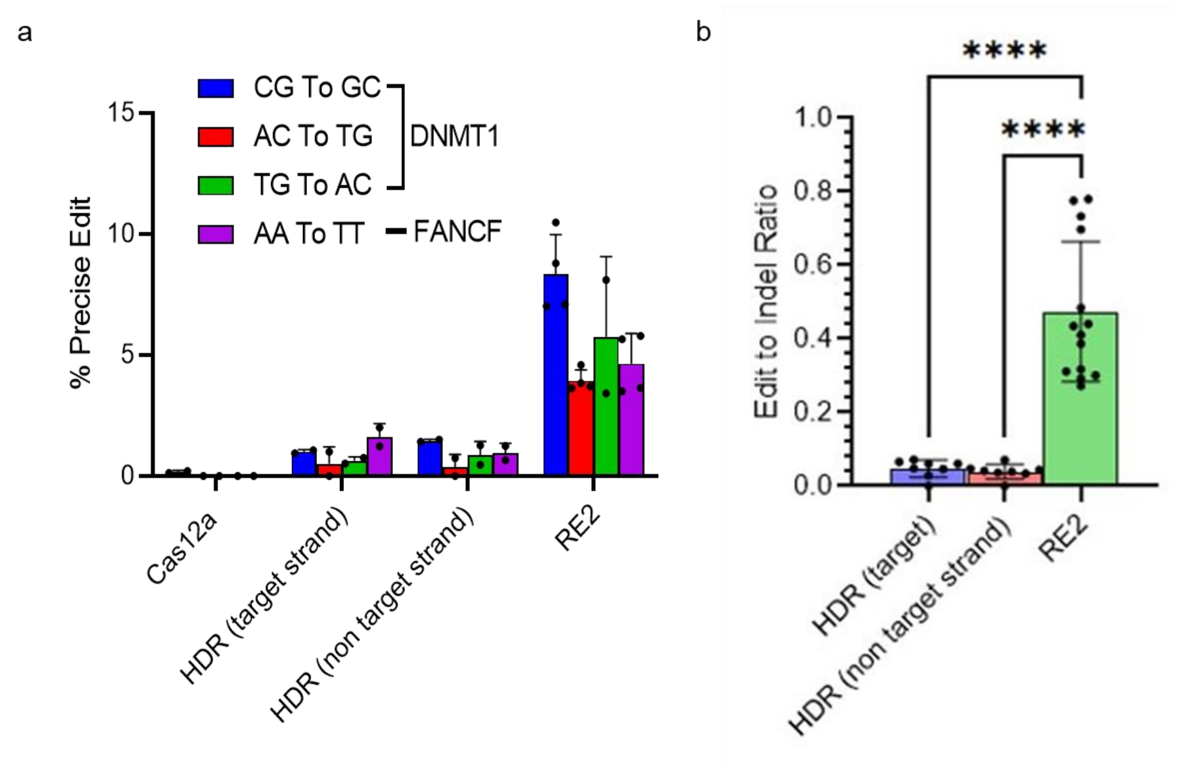


**Supplementary Figure 8. REDRAW comparison with HDR.** (a) LbCas12a nuclease without donor DNA, Cas12a HDR with both target and non-target strand DNA donor, and RE2 were compared for their ability to introduce four different double transversion mutations at the DNMT1 and FANCF sites. (b) RE2 displays an average 9.8-fold increase in edit-to-indel ratio compared to that of Cas12a-mediated HDR.

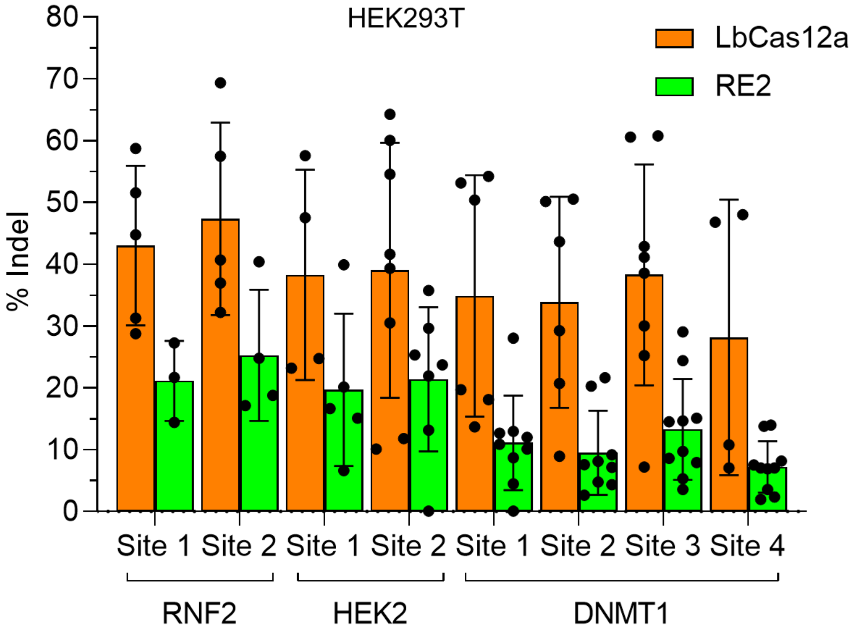


**Supplementary Figure 9. Random insertions and deletions generated during REDRAW.** REDRAW creates indel byproducts in HEK293T cells at all sites tested, but at an average frequency of 2.1-fold less than the indels caused by LbCas12a.

a

Site: DNMT1
On target spacer = GCTCAGCAGGCACCTGCCTCAGC
LbCas12a GUIDE-Seq reads: 297 (Chen, et al.), 946 (Kleinstiver, et al.)

Off-target site 1
Spacer: GCTCAGCAGGCACCTGCCCATGG
LbCas12a GUIDE-Seq reads: 394 (Chen, et al.), 683 (Kleinstiver, et al.)

Off-target site 2
Spacer: GCTCAGCTGACACCTGCCCACCA
LbCas12a GUIDE-Seq reads: 20 (Chen, et al.)

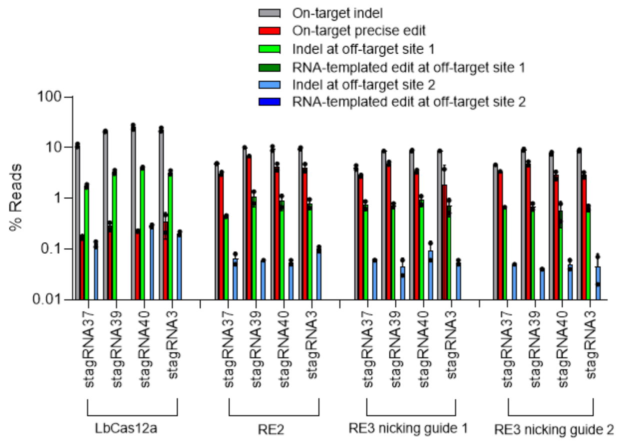


b

Site: DNMT1
On target spacer = CTGATGGTCCATGTCTGTTACTC
LbCas12a GUIDE-Seq reads: 982 (Kleinstiver, et al.)

Off-target site 1
Spacer: CTGATGGTCCATGTCTGAATTAG
LbCas12a GUIDE-Seq reads: 206 (Kleinstiver, et al.)

Off-target site 2
Spacer: CTGATGGTCCACGCCTGTTAACA
LbCas12a GUIDE-Seq reads: 5 (Kleinstiver, et al.)

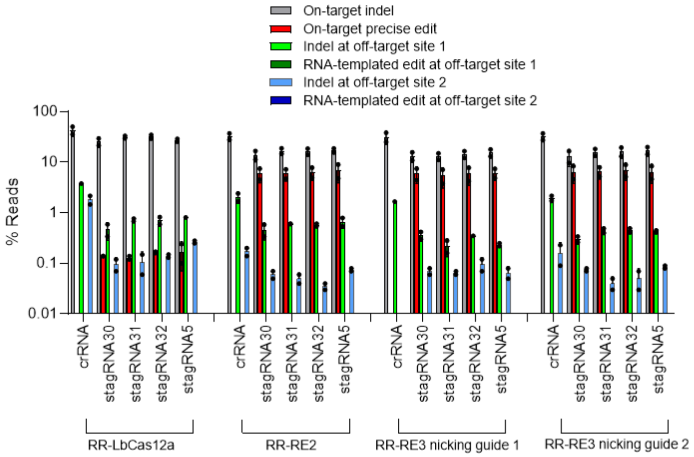


**Supplementary Figure 10. Off-target activity of REDRAW editors.** On-target and RNA-guided off-target activity is measured for RE2, RR-RE2, RE3, RR-RE3 constructs using four stagRNAs targeting previously identified Cas12a off-target sites in the DNMT1 locus. Two different nicking crRNAs are used for RE3 applications. (a) Off-target activity measured for RE2 and RE3. (b) Off-target activity measured for RR-RE2 and RR-RE3.

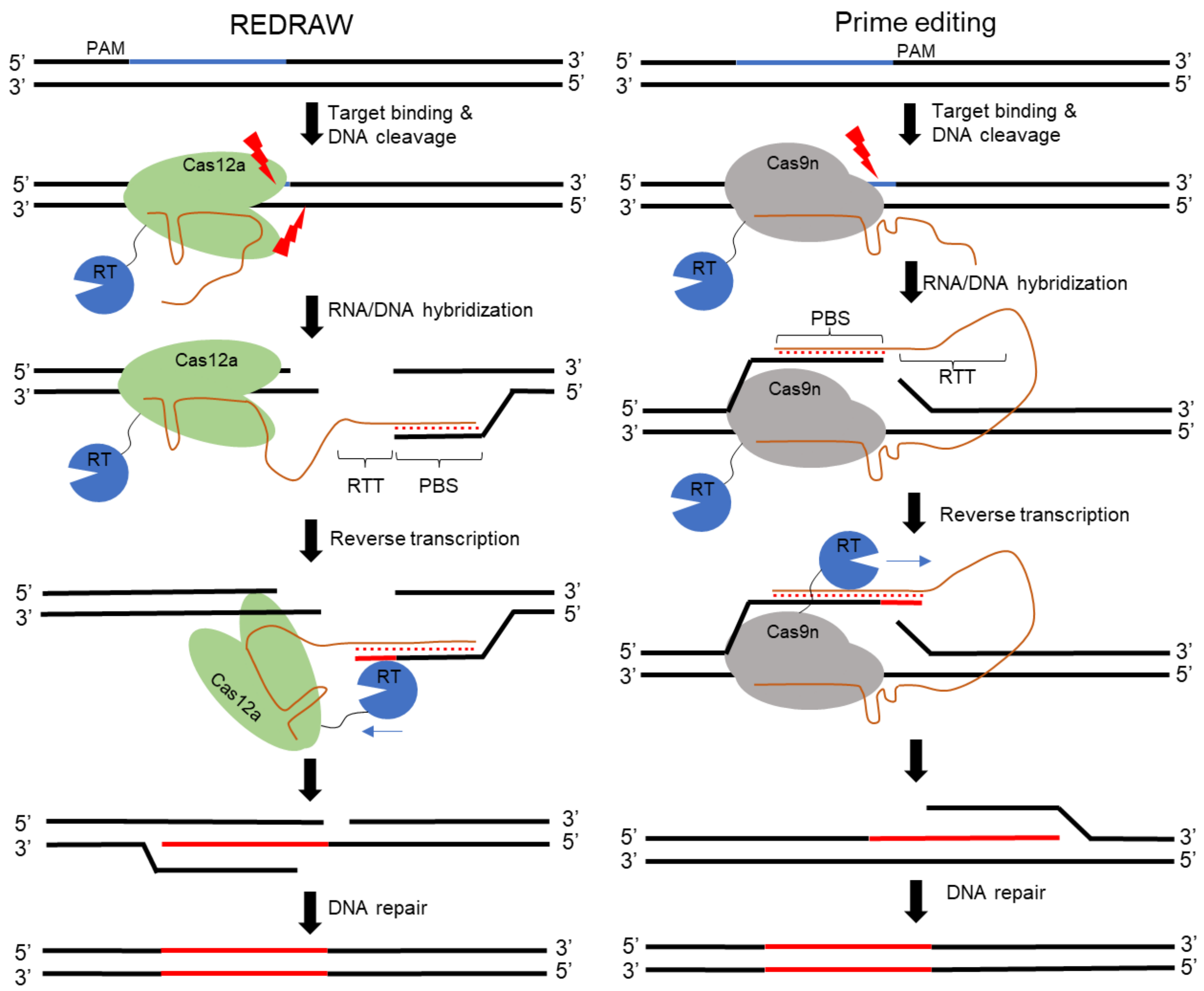


**Supplementary Figure 11. Comparison between REDRAW and Prime editing mechanisms.** Several key differences between REDRAW and Prime editing are shown, including PBS binding location and reverse transcription direction. Blue segments denote Cas12a or Cas9 binding sites. Red segments denote newly synthesized DNA strands that contain desired mutations. The lightning bolt symbol denotes a DNA cleavage event.

**Supplementary Note 1**

Engineered crRNAs used in this study:

| stagRNA ID | Locus | Enzyme | Spacer Sequence | Extension Sequence | 5' or 3' | PAM |
|---|---|---|---|---|---|---|
| stagRNA1 | DNMT1 | LbCas12a | CCTCACTCCTGCTCGGTGAATTT | ACAGCAGGCCTTTGGTCAGGTTGGCTGCTGGGCTGGCCCTGGGGCCGTTTCCCTCACTCCTGGACGGTGAATTTGGCTCAGCAGGCACCTGCCTCAGCTGCTCACTTGAGCCTCTGGGTCTA | 3' | TTTC |
| stagRNA2 | DNMT1 | LbCas12a | CCTCACTCCTGCTCGGTGAATTT | ACAGCAGGCCTTTGGTCAGGTTGGCTGCTGGGCTGGCCCTGGGGCCGTTTCCCTCACTCCTGCTGCGTGAATTTGGCTCAGCAGGCACCTGCCTCAGCTGCTCACTTGAGCCTCTGGGTCTA | 3' | TTTC |
| stagRNA3 | DNMT1 | LbCas12a | GCTCAGCAGGCACCTGCCTCAGC | GTTTCCCTCACTCCTGCTCGGTGAATTTGGCTCAGCAGGCTGCTGCCTCAGCTGCTCACTTGAGCCTCTGGGTCTAGAACCCTCTGGGGACCGTTTGAGG | 3' | TTTG |
| stagRNA4 | DNMT1 | LbCas12a | GCTCAGCAGGCACCTGCCTCAGC | GTTTCCCTCACTCCTGCTCGGTGAATTTGGCTCAGCAGGCACGAGCCTCAGCTGCTCACTTGAGCCTCTGGGTCTAGAACCCTCTGGGGACCGTTTGAGG | 3' | TTTG |
| stagRNA5 | DNMT1 | LbCas12a | CTGATGGTCCATGTCTGTTACTC | AGCTGTTAACATCAGTACGTTAATGTTTCCTGATGGTCCAACTCTGTTACTCGCCTGTCAAGTGGCGTGACACCGGGCGTGTTCCCCAGAGTGACTTTTC | 3' | TTTC |
| stagRNA6 | DNMT1 | LbCas12a | CTGATGGTCCATGTCTGTTACTC | AGCTGTTAACATCAGTACGTTAATGTTTCCTGATGGTCCATGAGTGTTACTCGCCTGTCAAGTGGCGTGACACCGGGCGTGTTCCCCAGAGTGACTTTTC | 3' | TTTC |
| stagRNA7 | FANCF | LbCas12a | GCGGATGTTCCAATCAGTACGCA | AATAGCATTGCAGAGAGGCGTATCATTTCGCGGATGTTGGAATCAGTACGCAGAGAGTCGCCGTCTCCAAGGTGAAAGCGGAAGTAGGGCCTTCGCGCAC | 3' | TTTC |
| stagRNA8 | FANCF | LbCas12a | GCGGATGTTCCAATCAGTACGCA | AATAGCATTGCAGAGAGGCGTATCATTTCGCGGATGTTCCTTTCAGTACGCAGAGAGTCGCCGTCTCCAAGGTGAAAGCGGAAGTAGGGCCTTCGCGCAC | 3' | TTTC |
| stagRNA9 | RNF2 | LbCas12a | TATGAGTTACAACGAACACCTCA | CATTAAGCAAAACATGGGAACTCAGTTTATATGAGTTACAtgGAACACCTCAGGTAATGACTAAGATGACTGCCAAGGGGCATATGAGACGTGTAAACTG | 3' | TTTA |
| stagRNA10 | RNF2 | LbCas12a | TATGAGTTACAACGAACACCTCA | CATTAAGCAAAACATGGGAACTCAGTTTATATGAGTTACAgtGAACACCTCAGGTAATGACTAAGATGACTGCCAAGGGGCATATGAGACGTGTAAACTG | 3' | TTTA |
| stagRNA11 | RNF2 | LbCas12a | TATGAGTTACAACGAACACCTCA | CATTAAGCAAAACATGGGAACTCAGTTTATATGAGTTACAACACCTCAGGTAATGACTAAGATGACTGCCAAGGGGCATATGAGACGTGTAAACTG | 3' | TTTA |
| stagRNA12 | RNF2 | LbCas12a | CACGTCTCATATGCCCCTTGGCA | TTCATGTTCTAAAAATGTATCCCAGTTTACACGTCTCATAacCCCCTTGGCAGTCATCTTAGTCATTACCTGAGGTGTTCGTTGTAACTCATATAAACTG | 3' | TTTA |
| stagRNA13 | RNF2 | LbCas12a | CACGTCTCATATGCCCCTTGGCA | TTCATGTTCTAAAAATGTATCCCAGTTTACACGTCTCATAcaCCCCTTGGCAGTCATCTTAGTCATTACCTGAGGTGTTCGTTGTAACTCATATAAACTG | 3' | TTTA |
| stagRNA14 | RNF2 | LbCas12a | CACGTCTCATATGCCCCTTGGCA | TTCATGTTCTAAAAATGTATCCCAGTTTACACGTCTCATACCTTGGCAGTCATCTTAGTCATTACCTGAGGTGTTCGTTGTAACTCATATAAACTG | 3' | TTTA |
| stagRNA15 | HEK2 | LbCas12a | CAGCCCGCTGGCCCTGTAAAGGA | GCTAACTGTGACAGCATGTGGTAATTTTCCAGCCCGCTGGggCTGTAAAGGAAACTGGAACACAAAGCATAGACTGCGGGGCGGGCCAGCCTGAATAGCT | 3' | TTTC |
| stagRNA16 | HEK2 | LbCas12a | CAGCCCGCTGGCCCTGTAAAGGA | GCTAACTGTGACAGCATGTGGTAATTTTCCAGCCCGCTGGttCTGTAAAGGAAACTGGAACACAAAGCATAGACTGCGGGGCGGGCCAGCCTGAATAGCT | 3' | TTTC |
| stagRNA17 | HEK2 | LbCas12a | CAGCCCGCTGGCCCTGTAAAGGA | GCTAACTGTGACAGCATGTGGTAATTTTCCAGCCCGCTGGGTAAAGGAAACTGGAACACAAAGCATAGACTGCGGGGCGGGCCAGCCTGAATAGCT | 3' | TTTC |
| stagRNA18 | HEK2 | LbCas12a | CAGCTATTCAGGCTGGCCCGCCC | GACATCATCAGATATTCTGCACTTGTTTGCAGCTATTCAGcgTGGCCCGCCCCGCAGTCTATGCTTTGTGTTCCAGTTTCCTTTACAGGGCCAGCGGGCT | 3' | TTTG |
| stagRNA19 | HEK2 | LbCas12a | CAGCTATTCAGGCTGGCCCGCCC | GACATCATCAGATATTCTGCACTTGTTTGCAGCTATTCAGatTGGCCCGCCCCGCAGTCTATGCTTTGTGTTCCAGTTTCCTTTACAGGGCCAGCGGGCT | 3' | TTTG |
| stagRNA20 | HEK2 | LbCas12a | CAGCTATTCAGGCTGGCCCGCCC | GACATCATCAGATATTCTGCACTTGTTTGCAGCTATTCAGGCCCGCCCCGCAGTCTATGCTTTGTGTTCCAGTTTCCTTTACAGGGCCAGCGGGCT | 3' | TTTG |
| stagRNA21 | DNMT1 | LbCas12a | CCTTCAGCTAAAATAAAGGAGGA | TTCCCCAGAGTGACTTTTCCTTTTATTTCCCTTCAGCTAAttTAAAGGAGGAGGAAGCTGCTAAGGACTAGTTCTGCCCTCCCGTCACCCCTGTTTCTGG | 3' | TTTC |
| stagRNA22 | DNMT1 | LbCas12a | CCTTCAGCTAAAATAAAGGAGGA | TTCCCCAGAGTGACTTTTCCTTTTATTTCCCTTCAGCTAAggTAAAGGAGGAGGAAGCTGCTAAGGACTAGTTCTGCCCTCCCGTCACCCCTGTTTCTGG | 3' | TTTC |
| stagRNA23 | DNMT1 | LbCas12a | CCTTCAGCTAAAATAAAGGAGGA | TTCCCCAGAGTGACTTTTCCTTTTATTTCCCTTCAGCTAAAAGGAGGAGGAAGCTGCTAAGGACTAGTTCTGCCCTCCCGTCACCCCTGTTTCTGG | 3' | TTTC |
| stagRNA24 | DNMT1 | LbCas12a | TTTCCCTTCAGCTAAAATAAAGG | CGTGTTCCCCAGAGTGACTTTTCCTTTTATTTCCCTTCAGgaAAAATAAAGGAGGAGGAAGCTGCTAAGGACTAGTTCTGCCCTCCCGTCACCCCTGTTT | 3' | TTTA |
| stagRNA25 | DNMT1 | LbCas12a | TTTCCCTTCAGCTAAAATAAAGG | CGTGTTCCCCAGAGTGACTTTTCCTTTTATTTCCCTTCAGtcAAAATAAAGGAGGAGGAAGCTGCTAAGGACTAGTTCTGCCCTCCCGTCACCCCTGTTT | 3' | TTTA |
| stagRNA26 | DNMT1 | LbCas12a | TTTCCCTTCAGCTAAAATAAAGG | CGTGTTCCCCAGAGTGACTTTTCCTTTTATTTCCCTTCAGAATAAAGGAGGAGGAAGCTGCTAAGGACTAGTTCTGCCCTCCCGTCACCCCTGTTT | 3' | TTTA |
| stagRNA27 | DNMT1 | LbCas12a | CTGATGGTCCATGTCTGTTACTC | AGCTGTTAACATCAGTACGTTAATGTTTCCTGATGGTCCAAGTCTGTTACTCGCCTGTCAAGTGGCGTGACACCGGGCGTGTTCCCCAGAGTGACTTTTC | 3' | TTTC |
| stagRNA28 | DNMT1 | LbCas12a | CTGATGGTCCATGTCTGTTACTC | AGCTGTTAACATCAGTACGTTAATGTTTCCTGATGGTCCATCTCTGTTACTCGCCTGTCAAGTGGCGTGACACCGGGCGTGTTCCCCAGAGTGACTTTTC | 3' | TTTC |
| stagRNA29 | DNMT1 | LbCas12a | CTGATGGTCCATGTCTGTTACTC | AGCTGTTAACATCAGTACGTTAATGTTTCCTGATGGTCCATGACTGTTACTCGCCTGTCAAGTGGCGTGACACCGGGCGTGTTCCCCAGAGTGACTTTTC | 3' | TTTC |
| stagRNA30 | DNMT1 | LbCas12a | CTGATGGTCCATGTCTGTTACTC | AGCTGTTAACATCAGTACGTTAATGTTTCCTGATGGTCCATGTGTGTTACTCGCCTGTCAAGTGGCGTGACACCGGGCGTGTTCCCCAGAGTGACTTTTC | 3' | TTTC |

| | | | | | | |
|---|---|---|---|---|---|---|
| stagRNA31 | DNMT1 | LbCas12a | CTGATGGTCCATGTCTGTTACTC | AGCTGTTAACATCAGTACGTTAATGTTTCCTGATGGTCCATCACTGTTACTCGCCTGTCAAGTGGCGTGACACCGGGCGTGTTCCCCAGAGTGACTTTTC | 3' | TTTC |
| stagRNA32 | DNMT1 | LbCas12a | CTGATGGTCCATGTCTGTTACTC | AGCTGTTAACATCAGTACGTTAATGTTTCCTGATGGTCCAACAGTGTTACTCGCCTGTCAAGTGGCGTGACACCGGGCGTGTTCCCCAGAGTGACTTTTC | 3' | TTTC |
| stagRNA35 | DNMT1 | LbCas12a | GCTCAGCAGGCACCTGCCTCAGC | GTTTCCCTCACTCCTGCTCGGTGAATTTGGCTCAGCAGGCTCCTGCCTCAGCTGCTCACTTGAGCCTCTGGGTCTAGAACCCTCTGGGGACCGTTTGAGG | 3' | TTTG |
| stagRNA36 | DNMT1 | LbCas12a | GCTCAGCAGGCACCTGCCTCAGC | GTTTCCCTCACTCCTGCTCGGTGAATTTGGCTCAGCAGGCAGCTGCCTCAGCTGCTCACTTGAGCCTCTGGGTCTAGAACCCTCTGGGGACCGTTTGAGG | 3' | TTTG |
| stagRNA37 | DNMT1 | LbCas12a | GCTCAGCAGGCACCTGCCTCAGC | GTTTCCCTCACTCCTGCTCGGTGAATTTGGCTCAGCAGGCACGTGCCTCAGCTGCTCACTTGAGCCTCTGGGTCTAGAACCCTCTGGGGACCGTTTGAGG | 3' | TTTG |
| stagRNA38 | DNMT1 | LbCas12a | GCTCAGCAGGCACCTGCCTCAGC | GTTTCCCTCACTCCTGCTCGGTGAATTTGGCTCAGCAGGCACCAGCCTCAGCTGCTCACTTGAGCCTCTGGGTCTAGAACCCTCTGGGGACCGTTTGAGG | 3' | TTTG |
| stagRNA39 | DNMT1 | LbCas12a | GCTCAGCAGGCACCTGCCTCAGC | GTTTCCCTCACTCCTGCTCGGTGAATTTGGCTCAGCAGGCAGGTGCCTCAGCTGCTCACTTGAGCCTCTGGGTCTAGAACCCTCTGGGGACCGTTTGAGG | 3' | TTTG |
| stagRNA40 | DNMT1 | LbCas12a | GCTCAGCAGGCACCTGCCTCAGC | GTTTCCCTCACTCCTGCTCGGTGAATTTGGCTCAGCAGGCTGCAGCCTCAGCTGCTCACTTGAGCCTCTGGGTCTAGAACCCTCTGGGGACCGTTTGAGG | 3' | TTTG |
| stagRNA41 | DNMT1 | enAsCas12a | CCTCACTCCTGCTCGGTGAATTT | ACAGCAGGCCTTTGGTCAGGTTGGCTGCTGGGCTGGCCCTGGGGCCGTTTCCCTCACTCCTGGACGGTGAATTTGGCTCAGCAGGCACCTGCCTCAGCTGCTCACTTGAGCCTCTGGGTCTA | 3' | TTTC |
| stagRNA42 | DNMT1 | enAsCas12a | CCTCACTCCTGCTCGGTGAATTT | GGCTGCTGGGCTGGCCCTGGGGCCGTTTCCCTCACTCCTGGACGGTGAATTTGGCTCAGCAGGCACCTGCCTCAGCTGCTCACTTGAGCCTCTGGGTCTA | 3' | TTTC |
| stagRNA43 | DNMT1 | enAsCas12a | CCTCACTCCTGCTCGGTGAATTT | ACAGCAGGCCTTTGGTCAGGTTGGCTGCTGGGCTGGCCCTGGGGCCGTTTCCCTCACTCCTGGAGCGTGAATTTGGCTCAGCAGGCACCTGCCTCAGCTGCTCACTTGAGCCTCTGGGTCTA | 3' | TTTC |
| stagRNA44 | DNMT1 | enAsCas12a | CCTCACTCCTGCTCGGTGAATTT | GGCTGCTGGGCTGGCCCTGGGGCCGTTTCCCTCACTCCTGGAGCGTGAATTTGGCTCAGCAGGCACCTGCCTCAGCTGCTCACTTGAGCCTCTGGGTCTA | 3' | TTTC |
| stagRNA45 | DNMT1 | enAsCas12a | GCTCAGCAGGCACCTGCCTCAGC | CTGCTGGGCTGGCCCTGGGGCCGTTTCCCTCACTCCTGCTCGGTGAATTTGGCTCAGCAGGCTGCTGCCTCAGCTGCTCACTTGAGCCTCTGGGTCTAGAACCCTCTGGGGACCGTTTGAGG | 3' | TTTG |
| stagRNA46 | DNMT1 | enAsCas12a | GCTCAGCAGGCACCTGCCTCAGC | GTTTCCCTCACTCCTGCTCGGTGAATTTGGCTCAGCAGGCTGCTGCCTCAGCTGCTCACTTGAGCCTCTGGGTCTAGAACCCTCTGGGGACCGTTTGAGG | 3' | TTTG |
| stagRNA47 | DNMT1 | enAsCas12a | GCTCAGCAGGCACCTGCCTCAGC | CTGCTGGGCTGGCCCTGGGGCCGTTTCCCTCACTCCTGCTCGGTGAATTTGGCTCAGCAGGCTGGAGCCTCAGCTGCTCACTTGAGCCTCTGGGTCTAGAACCCTCTGGGGACCGTTTGAGG | 3' | TTTG |
| stagRNA48 | DNMT1 | enAsCas12a | GCTCAGCAGGCACCTGCCTCAGC | GTTTCCCTCACTCCTGCTCGGTGAATTTGGCTCAGCAGGCTGGAGCCTCAGCTGCTCACTTGAGCCTCTGGGTCTAGAACCCTCTGGGGACCGTTTGAGG | 3' | TTTG |
| stagRNA49 | RNF2 | enAsCas12a | AAGCATTCCTGACTTCTGTATGT | TTAGAACATGAAAACTTAAATAGAACATAGAGTAATAATAAATTTATATTCAAGCATTCCTGTGTTCTGTATGTTGGCTAACACTTTATTCATTGATTGAACAAATAATCTGAATACTCTAT | 3' | ATTC |
| stagRNA50 | RNF2 | enAsCas12a | AAGCATTCCTGACTTCTGTATGT | GAACATAGAGTAATAATAAATTTATATTCAAGCATTCCTGTGTTCTGTATGTTGGCTAACACTTTATTCATTGATTGAACAAATAATCTGAATACTCTAT | 3' | ATTC |
| stagRNA51 | RNF2 | enAsCas12a | AAGCATTCCTGACTTCTGTATGT | TTAGAACATGAAAACTTAAATAGAACATAGAGTAATAATAAATTTATATTCAAGCATTCCTGTGAACTGTATGTTGGCTAACACTTTATTCATTGATTGAACAAATAATCTGAATACTCTAT | 3' | ATTC |
| stagRNA52 | RNF2 | enAsCas12a | TGAGTTACAACGAACACCTCAGG | GCAGACAAACGGAACTCAACCATTAAGCAAAACATGGGAACTCAGTTTATATGAGTTACAACCTACACCTCAGGTAATGACTAAGATGACTGCCAAGGGGCATATGAGACGTGTAAACTGGG | 3' | TATA |
| stagRNA53 | RNF2 | enAsCas12a | TGAGTTACAACGAACACCTCAGG | TTAAGCAAAACATGGGAACTCAGTTTATATGAGTTACAACCTACACCTCAGGTAATGACTAAGATGACTGCCAAGGGGCATATGAGACGTGTAAACTGGG | 3' | TATA |
| stagRNA54 | RNF2 | enAsCas12a | TGAGTTACAACGAACACCTCAGG | GCAGACAAACGGAACTCAACCATTAAGCAAAACATGGGAACTCAGTTTATATGAGTTACAACCTTGACCTCAGGTAATGACTAAGATGACTGCCAAGGGGCATATGAGACGTGTAAACTGGG | 3' | TATA |
| stagRNA55 | RNF2 | enAsCas12a | TGAGTTACAACGAACACCTCAGG | TTAAGCAAAACATGGGAACTCAGTTTATATGAGTTACAACCTTGACCTCAGGTAATGACTAAGATGACTGCCAAGGGGCATATGAGACGTGTAAACTGGG | 3' | TATA |
| stagRNA56 | RNF2 | enAsCas12a | TTCAAGCATTCCTGACTTCTGTA | TTTTTAGAACATGAAAACTTAAATAGAACATAGAGTAATAATAAATTTATATTCAAGCATTCGAGACTTCTGTATGTTGGCTAACACTTTATTCATTGATTGAACAAATAATCTGAATACTC | 3' | TATA |
| stagRNA57 | RNF2 | enAsCas12a | TTCAAGCATTCCTGACTTCTGTA | ATAGAACATAGAGTAATAATAAATTTATATTCAAGCATTCGAGACTTCTGTATGTTGGCTAACACTTTATTCATTGATTGAACAAATAATCTGAATACTC | 3' | TATA |
| stagRNA58 | RNF2 | enAsCas12a | TTCAAGCATTCCTGACTTCTGTA | TTTTTAGAACATGAAAACTTAAATAGAACATAGAGTAATAATAAATTTATATTCAAGCATTCGACTCTTCTGTATGTTGGCTAACACTTTATTCATTGATTGAACAAATAATCTGAATACTC | 3' | TATA |
| stagRNA59 | RNF2 | enAsCas12a | TTCAAGCATTCCTGACTTCTGTA | ATAGAACATAGAGTAATAATAAATTTATATTCAAGCATTCGACTCTTCTGTATGTTGGCTAACACTTTATTCATTGATTGAACAAATAATCTGAATACTC | 3' | TATA |
| stagRNA60 | FANCF | enAsCas12a | ACTGGTTGTGCAGCCGCCGCTCC | AGTTCGCTAATCCCGGAACTGGACCCCGCCCAAAGCCGCCCTCTTGCCTCCACTGGTTGTGCTCCCGCCGCTCCAGAGCCGTGCGAATGGGGCCATGCCGACCAAAGCGCCGATGGATGTGG | 3' | CTCC |

| | | | | | | |
|---|---|---|---|---|---|---|
| stagRNA61 | FANCF | enAsCas12a | ACTGGTTGTGCAGCCGCCGCTCC | ACCCCGCCCAAAGCCGCCCTCTTGCCTCCACTGGTTGTGCTCCCGCCGCTCCAGAGCCGTGCGAATGGGGCCATGCCGACCAAAGCGCCGATGGATGTGG | 3' | CTCC |
| stagRNA62 | FANCF | enAsCas12a | ACTGGTTGTGCAGCCGCCGCTCC | AGTTCGCTAATCCCGGAACTGGACCCCGCCCAAAGCCGCCCTCTTGCCTCCACTGGTTGTGCTCGGGCCGCTCCAGAGCCGTGCGAATGGGGCCATGCCGACCAAAGCGCCGATGGATGTGG | 3' | CTCC |
| stagRNA63 | FANCF | enAsCas12a | ACTGGTTGTGCAGCCGCCGCTCC | ACCCCGCCCAAAGCCGCCCTCTTGCCTCCACTGGTTGTGCTCGGGCCGCTCCAGAGCCGTGCGAATGGGGCCATGCCGACCAAAGCGCCGATGGATGTGG | 3' | CTCC |
| stagRNA64 | HEK2 | BhCas12b | TCCAGCCCGCTGGCCCTGTAAA | AACAATGATAACAAGACCTGGCTGAGCTAACTGTGACAGCATGTGGTAATTTTCCAGCCCGCTGCGCCTGTAAAGGAAACTGGAACACAAAGCATAGACTGCGGGGCGGGCCAGCCTGAATA | 3' | ATTT |
| stagRNA65 | HEK2 | BhCas12b | TCCAGCCCGCTGGCCCTGTAAA | TGAGCTAACTGTGACAGCATGTGGTAATTTTCCAGCCCGCTGCGCCTGTAAAGGAAACTGGAACACAAAGCATAGACTGCGGGGCGGGCCAGCCTGAATA | 3' | ATTT |
| stagRNA66 | HEK2 | BhCas12b | AGGCTGGCCCGCCCCGCAGTCT | CAAACTGTGCGTATGACATCATCAGATATTCTGCACTTGTTTGCAGCTATTCAGGCTGGCCCGGGCCGCAGTCTATGCTTTGTGTTCCAGTTTCCTTTACAGGGCCAGCGGGCTGGAAAATT | 3' | ATTC |
| stagRNA67 | HEK2 | BhCas12b | AGGCTGGCCCGCCCCGCAGTCT | CAGATATTCTGCACTTGTTTGCAGCTATTCAGGCTGGCCCGGGCCGCAGTCTATGCTTTGTGTTCCAGTTTCCTTTACAGGGCCAGCGGGCTGGAAAATT | 3' | ATTC |
| stagRNA68 | DNMT1 | BhCas12b | GGTCAGCTGTTAACATCAGTAC | TGAGGAGTGTTCAGTCTCCGTGAACGTTCCCTTAGCACTCTGCCACTTATTGGGTCAGCTGTTCGCATCAGTACGTTAATGTTTCCTGATGGTCCATGTCTGTTACTCGCCTGTCAAGTGGC | 3' | ATTG |
| stagRNA69 | DNMT1 | BhCas12b | GGTCAGCTGTTAACATCAGTAC | AACGTTCCCTTAGCACTCTGCCACTTATTGGGTCAGCTGTTCGCATCAGTACGTTAATGTTTCCTGATGGTCCATGTCTGTTACTCGCCTGTCAAGTGGC | 3' | ATTG |
| stagRNA70 | DNMT1 | BhCas12b | ACGTACTGATGTTAACAGCTGA | TGTCACGCCACTTGACAGGCGAGTAACAGACATGGACCATCAGGAAACATTAACGTACTGATGACAACAGCTGACCCAATAAGTGGCAGAGTGCTAAGGGAACGTTCACGGAGACTGAACAC | 3' | ATTA |
| stagRNA71 | DNMT1 | BhCas12b | ACGTACTGATGTTAACAGCTGA | GTAACAGACATGGACCATCAGGAAACATTAACGTACTGATGACAACAGCTGACCCAATAAGTGGCAGAGTGCTAAGGGAACGTTCACGGAGACTGAACAC | 3' | ATTA |
| tagRNA1 | DNMT1 | LbCas12a | CCTCACTCCTGCTCGGTGAATTT | ACAGCAGGCCTTTGGTCAGGTTGGCTGCTGGGCTGGCCCTGGGGCCGTTTCCCTCACTCCTGGACGGTGAATTTGGCTCAGCAGGCACCTGCCTCAGCTGCTCACTTGAGCCTCTGGGTCTA | 3' | TTTC |
| tagRNA2 | DNMT1 | LbCas12a | CCTCACTCCTGCTCGGTGAATTT | ACAGCAGGCCTTTGGTCAGGTTGGCTGCTGGGCTGGCCCTGGGGCCGTTTCCCTCACTCCTGCTGCGTGAATTTGGCTCAGCAGGCACCTGCCTCAGCTGCTCACTTGAGCCTCTGGGTCTA | 3' | TTTC |
| tagRNA3 | DNMT1 | LbCas12a | GCTCAGCAGGCACCTGCCTCAGC | GTTTCCCTCACTCCTGCTCGGTGAATTTGGCTCAGCAGGCTGCTGCCTCAGCTGCTCACTTGAGCCTCTGGGTCTAGAACCCTCTGGGGACCGTTTGAGG | 3' | TTTG |
| tagRNA4 | DNMT1 | LbCas12a | GCTCAGCAGGCACCTGCCTCAGC | GTTTCCCTCACTCCTGCTCGGTGAATTTGGCTCAGCAGGCACGAGCCTCAGCTGCTCACTTGAGCCTCTGGGTCTAGAACCCTCTGGGGACCGTTTGAGG | 3' | TTTG |
| tagRNA5 | DNMT1 | LbCas12a | CTGATGGTCCATGTCTGTTACTC | AGCTGTTAACATCAGTACGTTAATGTTTCCTGATGGTCCAACTCTGTTACTCGCCTGTCAAGTGGCGTGACACCGGGCGTGTTCCCCAGAGTGACTTTTC | 3' | TTTC |
| tagRNA6 | DNMT1 | LbCas12a | CTGATGGTCCATGTCTGTTACTC | AGCTGTTAACATCAGTACGTTAATGTTTCCTGATGGTCCATGAGTGTTACTCGCCTGTCAAGTGGCGTGACACCGGGCGTGTTCCCCAGAGTGACTTTTC | 3' | TTTC |
| tagRNA7 | FANCF | LbCas12a | GCGGATGTTCCAATCAGTACGCA | AATAGCATTGCAGAGAGGCGTATCATTTCGCGGATGTTGGAATCAGTACGCAGAGAGTCGCCGTCTCCAAGGTGAAAGCGGAAGTAGGGCCTTCGCGCAC | 3' | TTTC |
| tagRNA8 | FANCF | LbCas12a | GCGGATGTTCCAATCAGTACGCA | AATAGCATTGCAGAGAGGCGTATCATTTCGCGGATGTTCCTTTCAGTACGCAGAGAGTCGCCGTCTCCAAGGTGAAAGCGGAAGTAGGGCCTTCGCGCAC | 3' | TTTC |
| tagRNA9 | EMX1 | Cas9 | GAGTCCGAGCAGAAGAAGAA | AGTACAAACGGCAGAAGCTGGAGGAGGAAGGGCCTGAGTCCGAGCAGAACTTGAAGGGCTCCCATCACATCAACCGGTGGCGCATTGCCACGAAGCAGGC | 3' | GGG |
| tagRNA10 | EMX1 | Cas9 | GAGTCCGAGCAGAAGAAGAA | AGTACAAACGGCAGAAGCTGGAGGAGGAAGGGCCTGAGTCCGAGCAGAAAGGGAAGGGCTCCCATCACATCAACCGGTGGCGCATTGCCACGAAGCAGGC | 3' | GGG |
| tagRNA11 | FANCF | Cas9 | GGAATCCCTTCTGCAGCACC | GGTGAAAGCGGAAGTAGGGCCTTCGCGCACCTCATGGAATCCCTTCTGCTCGACCTGGATCGCTTTTCCGAGCTTCTGGCGGTCTCAAGCACTACCTACG | 3' | TGG |
| tagRNA12 | FANCF | Cas9 | GGAATCCCTTCTGCAGCACC | GGTGAAAGCGGAAGTAGGGCCTTCGCGCACCTCATGGAATCCCTTCTGCGATACCTGGATCGCTTTTCCGAGCTTCTGGCGGTCTCAAGCACTACCTACG | 3' | TGG |
| tagRNA13 | RNF2 | Cas9 | GTCATCTTAGTCATTACCTG | TATCCCAGTTTACACGTCTCATATGCCCCTTGGCAGTCATCTTAGTCATACGCTGAGGTGTTCGTTGTAACTCATATAAACTGAGTTCCCATGTTTTGCT | 3' | AGG |
| tagRNA14 | RNF2 | Cas9 | GTCATCTTAGTCATTACCTG | TATCCCAGTTTACACGTCTCATATGCCCCTTGGCAGTCATCTTAGTCATCGTCTGAGGTGTTCGTTGTAACTCATATAAACTGAGTTCCCATGTTTTGCT | 3' | AGG |
| tagRNA15 | DNMT1 | LbCas12a | CCTCACTCCTGCTCGGTGAATTT | ACAGCAGGCCTTTGGTCAGGTTGGCTGCTGGGCTGGCCCTGGGGCCGTAACCCTCACTCCTGCTCGGTGAATTTGGCTCAGCAGGCACCTGCCTCAGCTGCTCACTTGAGCCTCTGGGTCTA | 3' | TTTC |
| tagRNA16 | DNMT1 | LbCas12a | CCTCACTCCTGCTCGGTGAATTT | GGCTGCTGGGCTGGCCCTGGGGCCGTAACCCTCACTCCTGCTCGGTGAATTTGGCTCAGCAGGCACCTGCCTCAGCTGCTCACTTGAGCCTCTGGGTCTA | 3' | TTTC |
| tagRNA17 | DNMT1 | LbCas12a | CCTCACTCCTGCTCGGTGAATTT | GGCTGGCCCTGGGGCCGTAACCCTCACTCCTGCTCGGTGAATTTGGCTCAGCAGGCACCTGCCTCAGCTGCTCACTTGAGCCTCTGGGTCTA | 3' | TTTC |
| tagRNA18 | DNMT1 | LbCas12a | CCTCACTCCTGCTCGGTGAATTT | CTGGGGCCGTAACCCTCACTCCTGCTCGGTGAATTTGGCTCAGCAGGCACCTGCCTCAGCTGCTCACTTGAGCCTCTGGGTCTA | 3' | TTTC |
| tagRNA19 | DNMT1 | LbCas12a | CCTCACTCCTGCTCGGTGAATTT | CTGGGGCCGTAACCCTCACTCCTGCTCGGTGAATTTGGCTCAGCAGGCACCTGCCTCAGCTGCTCACTTGAGCCTCTGGGTCTA | 5' | TTTC |
| tagRNA20 | DNMT1 | LbCas12a | CCTCACTCCTGCTCGGTGAATTT | CTGGGGCCGTAACCCTCACTCCTGCTCGGTGAATTTGGCTCAGCAGGCACCTGCCTCAGC | 5' | TTTC |

Figure 1b

| PBS | RTT | 5' or 3' | PAM | Locus | Enzyme | Spacer Sequence | Extension Sequence |
|---|---|---|---|---|---|---|---|
| 0 | 0 | N/A | TTTC | DNMT1 | LbCas12a | CCTCACTCCTGCTCGGTGAATTT | N/A |
| 48 | 74 | 3' | TTTC | DNMT1 | LbCas12a | CCTCACTCCTGCTCGGTGAATTT | ACAGCAGGCCTTTGGTCAGGTTGGCTGCTGGGCTGGCCCTGGGGCCGTAACCCTCACTCCTGCTCGGTGAATTTGGCTCAGCAGGCACCTGCCTCAGCTGCTCACTTGAGCCTCTGGGTCTA |
| 48 | 52 | 3' | TTTC | DNMT1 | LbCas12a | CCTCACTCCTGCTCGGTGAATTT | GGCTGCTGGGCTGGCCCTGGGGCCGTAACCCTCACTCCTGCTCGGTGAATTTGGCTCAGCAGGCACCTGCCTCAGCTGCTCACTTGAGCCTCTGGGTCTA |
| 48 | 44 | 3' | TTTC | DNMT1 | LbCas12a | CCTCACTCCTGCTCGGTGAATTT | GGCTGGCCCTGGGGCCGTAACCCTCACTCCTGCTCGGTGAATTTGGCTCAGCAGGCACCTGCCTCAGCTGCTCACTTGAGCCTCTGGGTCTA |
| 48 | 36 | 3' | TTTC | DNMT1 | LbCas12a | CCTCACTCCTGCTCGGTGAATTT | CTGGGGCCGTAACCCTCACTCCTGCTCGGTGAATTTGGCTCAGCAGGCACCTGCCTCAGCTGCTCACTTGAGCCTCTGGGTCTA |
| 24 | 74 | 3' | TTTC | DNMT1 | LbCas12a | CCTCACTCCTGCTCGGTGAATTT | ACAGCAGGCCTTTGGTCAGGTTGGCTGCTGGGCTGGCCCTGGGGCCGTAACCCTCACTCCTGCTCGGTGAATTTGGCTCAGCAGGCACCTGCCTCAGC |
| 24 | 52 | 3' | TTTC | DNMT1 | LbCas12a | CCTCACTCCTGCTCGGTGAATTT | GGCTGCTGGGCTGGCCCTGGGGCCGTAACCCTCACTCCTGCTCGGTGAATTTGGCTCAGCAGGCACCTGCCTCAGC |
| 24 | 44 | 3' | TTTC | DNMT1 | LbCas12a | CCTCACTCCTGCTCGGTGAATTT | GGCTGGCCCTGGGGCCGTAACCCTCACTCCTGCTCGGTGAATTTGGCTCAGCAGGCACCTGCCTCAGC |
| 24 | 36 | 3' | TTTC | DNMT1 | LbCas12a | CCTCACTCCTGCTCGGTGAATTT | CTGGGGCCGTAACCCTCACTCCTGCTCGGTGAATTTGGCTCAGCAGGCACCTGCCTCAGC |
| 16 | 74 | 3' | TTTC | DNMT1 | LbCas12a | CCTCACTCCTGCTCGGTGAATTT | ACAGCAGGCCTTTGGTCAGGTTGGCTGCTGGGCTGGCCCTGGGGCCGTAACCCTCACTCCTGCTCGGTGAATTTGGCTCAGCAGGCACCT |
| 16 | 52 | 3' | TTTC | DNMT1 | LbCas12a | CCTCACTCCTGCTCGGTGAATTT | GGCTGCTGGGCTGGCCCTGGGGCCGTAACCCTCACTCCTGCTCGGTGAATTTGGCTCAGCAGGCACCT |
| 16 | 44 | 3' | TTTC | DNMT1 | LbCas12a | CCTCACTCCTGCTCGGTGAATTT | GGCTGGCCCTGGGGCCGTAACCCTCACTCCTGCTCGGTGAATTTGGCTCAGCAGGCACCT |
| 16 | 36 | 3' | TTTC | DNMT1 | LbCas12a | CCTCACTCCTGCTCGGTGAATTT | CTGGGGCCGTAACCCTCACTCCTGCTCGGTGAATTTGGCTCAGCAGGCACCT |
| 8 | 74 | 3' | TTTC | DNMT1 | LbCas12a | CCTCACTCCTGCTCGGTGAATTT | ACAGCAGGCCTTTGGTCAGGTTGGCTGCTGGGCTGGCCCTGGGGCCGTAACCCTCACTCCTGCTCGGTGAATTTGGCTCAGC |
| 8 | 52 | 3' | TTTC | DNMT1 | LbCas12a | CCTCACTCCTGCTCGGTGAATTT | GGCTGCTGGGCTGGCCCTGGGGCCGTAACCCTCACTCCTGCTCGGTGAATTTGGCTCAGC |
| 8 | 44 | 3' | TTTC | DNMT1 | LbCas12a | CCTCACTCCTGCTCGGTGAATTT | GGCTGGCCCTGGGGCCGTAACCCTCACTCCTGCTCGGTGAATTTGGCTCAGC |
| 8 | 36 | 3' | TTTC | DNMT1 | LbCas12a | CCTCACTCCTGCTCGGTGAATTT | CTGGGGCCGTAACCCTCACTCCTGCTCGGTGAATTTGGCTCAGC |
| 48 | 74 | 5' | TTTC | DNMT1 | LbCas12a | CCTCACTCCTGCTCGGTGAATTT | ACAGCAGGCCTTTGGTCAGGTTGGCTGCTGGGCTGGCCCTGGGGCCGTAACCCTCACTCCTGCTCGGTGAATTTGGCTCAGCAGGCACCTGCCTCAGCTGCTCACTTGAGCCTCTGGGTCTA |
| 48 | 52 | 5' | TTTC | DNMT1 | LbCas12a | CCTCACTCCTGCTCGGTGAATTT | GGCTGCTGGGCTGGCCCTGGGGCCGTAACCCTCACTCCTGCTCGGTGAATTTGGCTCAGCAGGCACCTGCCTCAGCTGCTCACTTGAGCCTCTGGGTCTA |
| 48 | 44 | 5' | TTTC | DNMT1 | LbCas12a | CCTCACTCCTGCTCGGTGAATTT | GGCTGGCCCTGGGGCCGTAACCCTCACTCCTGCTCGGTGAATTTGGCTCAGCAGGCACCTGCCTCAGCTGCTCACTTGAGCCTCTGGGTCTA |
| 48 | 36 | 5' | TTTC | DNMT1 | LbCas12a | CCTCACTCCTGCTCGGTGAATTT | CTGGGGCCGTAACCCTCACTCCTGCTCGGTGAATTTGGCTCAGCAGGCACCTGCCTCAGCTGCTCACTTGAGCCTCTGGGTCTA |
| 24 | 74 | 5' | TTTC | DNMT1 | LbCas12a | CCTCACTCCTGCTCGGTGAATTT | ACAGCAGGCCTTTGGTCAGGTTGGCTGCTGGGCTGGCCCTGGGGCCGTAACCCTCACTCCTGCTCGGTGAATTTGGCTCAGCAGGCACCTGCCTCAGC |
| 24 | 52 | 5' | TTTC | DNMT1 | LbCas12a | CCTCACTCCTGCTCGGTGAATTT | GGCTGCTGGGCTGGCCCTGGGGCCGTAACCCTCACTCCTGCTCGGTGAATTTGGCTCAGCAGGCACCTGCCTCAGC |
| 24 | 44 | 5' | TTTC | DNMT1 | LbCas12a | CCTCACTCCTGCTCGGTGAATTT | GGCTGGCCCTGGGGCCGTAACCCTCACTCCTGCTCGGTGAATTTGGCTCAGCAGGCACCTGCCTCAGC |
| 24 | 36 | 5' | TTTC | DNMT1 | LbCas12a | CCTCACTCCTGCTCGGTGAATTT | CTGGGGCCGTAACCCTCACTCCTGCTCGGTGAATTTGGCTCAGCAGGCACCTGCCTCAGC |

Figure 4a

| Transversion Position | PAM | Locus | Enzyme | Spacer Sequence | Extension Sequence |
|---|---|---|---|---|---|
| P12 | TTTC | DNMT1 | LbCas12a | CTGATGGTCCATGTCTGTTACTC | AGCTGTTAACATCAGTACGTTAATGTTTCCTGATGGTCCAAGTCTGTTACTCGCCTGTCAAGTGGCGTGACACCGGGCGTGTTCCCCAGAGTGACTTTTC |
| P13 | TTTC | DNMT1 | LbCas12a | CTGATGGTCCATGTCTGTTACTC | AGCTGTTAACATCAGTACGTTAATGTTTCCTGATGGTCCATCTCTGTTACTCGCCTGTCAAGTGGCGTGACACCGGGCGTGTTCCCCAGAGTGACTTTTC |
| P14 | TTTC | DNMT1 | LbCas12a | CTGATGGTCCATGTCTGTTACTC | AGCTGTTAACATCAGTACGTTAATGTTTCCTGATGGTCCATGACTGTTACTCGCCTGTCAAGTGGCGTGACACCGGGCGTGTTCCCCAGAGTGACTTTTC |
| P15 | TTTC | DNMT1 | LbCas12a | CTGATGGTCCATGTCTGTTACTC | AGCTGTTAACATCAGTACGTTAATGTTTCCTGATGGTCCATGTGTGTTACTCGCCTGTCAAGTGGCGTGACACCGGGCGTGTTCCCCAGAGTGACTTTTC |
| P12,P13 | TTTC | DNMT1 | LbCas12a | CTGATGGTCCATGTCTGTTACTC | AGCTGTTAACATCAGTACGTTAATGTTTCCTGATGGTCCAACTCTGTTACTCGCCTGTCAAGTGGCGTGACACCGGGCGTGTTCCCCAGAGTGACTTTTC |
| P13,P14 | TTTC | DNMT1 | LbCas12a | CTGATGGTCCATGTCTGTTACTC | AGCTGTTAACATCAGTACGTTAATGTTTCCTGATGGTCCATCACTGTTACTCGCCTGTCAAGTGGCGTGACACCGGGCGTGTTCCCCAGAGTGACTTTTC |
| P14,P15 | TTTC | DNMT1 | LbCas12a | CTGATGGTCCATGTCTGTTACTC | AGCTGTTAACATCAGTACGTTAATGTTTCCTGATGGTCCATGAGTGTTACTCGCCTGTCAAGTGGCGTGACACCGGGCGTGTTCCCCAGAGTGACTTTTC |
| P12,P13,P14,P15 | TTTC | DNMT1 | LbCas12a | CTGATGGTCCATGTCTGTTACTC | AGCTGTTAACATCAGTACGTTAATGTTTCCTGATGGTCCAACAGTGTTACTCGCCTGTCAAGTGGCGTGACACCGGGCGTGTTCCCCAGAGTGACTTTTC |
| P12 | TTTG | DNMT1 | LbCas12a | GCTCAGCAGGCACCTGCCTCAGC | GTTTCCCTCACTCCTGCTCGGTGAATTTGGCTCAGCAGGCTCCTGCCTCAGCTGCTCACTTGAGCCTCTGGGTCTAGAACCCTCTGGGGACCGTTTGAGG |
| P13 | TTTG | DNMT1 | LbCas12a | GCTCAGCAGGCACCTGCCTCAGC | GTTTCCCTCACTCCTGCTCGGTGAATTTGGCTCAGCAGGCAGCTGCCTCAGCTGCTCACTTGAGCCTCTGGGTCTAGAACCCTCTGGGGACCGTTTGAGG |
| P14 | TTTG | DNMT1 | LbCas12a | GCTCAGCAGGCACCTGCCTCAGC | GTTTCCCTCACTCCTGCTCGGTGAATTTGGCTCAGCAGGCACGTGCCTCAGCTGCTCACTTGAGCCTCTGGGTCTAGAACCCTCTGGGGACCGTTTGAGG |
| P15 | TTTG | DNMT1 | LbCas12a | GCTCAGCAGGCACCTGCCTCAGC | GTTTCCCTCACTCCTGCTCGGTGAATTTGGCTCAGCAGGCACCAGCCTCAGCTGCTCACTTGAGCCTCTGGGTCTAGAACCCTCTGGGGACCGTTTGAGG |
| P12,P13 | TTTG | DNMT1 | LbCas12a | GCTCAGCAGGCACCTGCCTCAGC | GTTTCCCTCACTCCTGCTCGGTGAATTTGGCTCAGCAGGCTGCTGCCTCAGCTGCTCACTTGAGCCTCTGGGTCTAGAACCCTCTGGGGACCGTTTGAGG |
| P13,P14 | TTTG | DNMT1 | LbCas12a | GCTCAGCAGGCACCTGCCTCAGC | GTTTCCCTCACTCCTGCTCGGTGAATTTGGCTCAGCAGGCAGGTGCCTCAGCTGCTCACTTGAGCCTCTGGGTCTAGAACCCTCTGGGGACCGTTTGAGG |
| P14,P15 | TTTG | DNMT1 | LbCas12a | GCTCAGCAGGCACCTGCCTCAGC | GTTTCCCTCACTCCTGCTCGGTGAATTTGGCTCAGCAGGCACGAGCCTCAGCTGCTCACTTGAGCCTCTGGGTCTAGAACCCTCTGGGGACCGTTTGAGG |
| P12,P13,P14,P15 | TTTG | DNMT1 | LbCas12a | GCTCAGCAGGCACCTGCCTCAGC | GTTTCCCTCACTCCTGCTCGGTGAATTTGGCTCAGCAGGCTGCAGCCTCAGCTGCTCACTTGAGCCTCTGGGTCTAGAACCCTCTGGGGACCGTTTGAGG |

Figure 4c

| Target Codon | PAM | Locus | Enzyme | Spacer Sequence | Extension Sequence |
|---|---|---|---|---|---|
| GCC | TTTC | DNMT1 | LbCas12a | CCTCACTCCTGCTCGGTGAATTT | ACAGCAGGCCTTTGGTCAGGTTGGCTGCTGGGCTGGCCCTGGGGCCGTTTCCCTCACTCCTGC<br>TGCCTGAATTTGGCTCAGCAGGCACCTGCCTCAGCTGCTCACTTGAGCCTCTGGGTCTA |
| ATT | TTTC | DNMT1 | LbCas12a | CCTCACTCCTGCTCGGTGAATTT | ACAGCAGGCCTTTGGTCAGGTTGGCTGCTGGGCTGGCCCTGGGGCCGTTTCCCTCACTCCTGC<br>TATTTGAATTTGGCTCAGCAGGCACCTGCCTCAGCTGCTCACTTGAGCCTCTGGGTCTA |
| TCC | TTTC | DNMT1 | LbCas12a | CCTCACTCCTGCTCGGTGAATTT | ACAGCAGGCCTTTGGTCAGGTTGGCTGCTGGGCTGGCCCTGGGGCCGTTTCCCTCACTCCTGC<br>TTCCTGAATTTGGCTCAGCAGGCACCTGCCTCAGCTGCTCACTTGAGCCTCTGGGTCTA |
| TTT | TTTC | DNMT1 | LbCas12a | CCTCACTCCTGCTCGGTGAATTT | ACAGCAGGCCTTTGGTCAGGTTGGCTGCTGGGCTGGCCCTGGGGCCGTTTCCCTCACTCCTGC<br>TTTTTGAATTTGGCTCAGCAGGCACCTGCCTCAGCTGCTCACTTGAGCCTCTGGGTCTA |
| GAC | TTTC | DNMT1 | LbCas12a | CCTCACTCCTGCTCGGTGAATTT | ACAGCAGGCCTTTGGTCAGGTTGGCTGCTGGGCTGGCCCTGGGGCCGTTTCCCTCACTCCTGC<br>TGACTGAATTTGGCTCAGCAGGCACCTGCCTCAGCTGCTCACTTGAGCCTCTGGGTCTA |
| AAT | TTTC | DNMT1 | LbCas12a | CCTCACTCCTGCTCGGTGAATTT | ACAGCAGGCCTTTGGTCAGGTTGGCTGCTGGGCTGGCCCTGGGGCCGTTTCCCTCACTCCTGC<br>TAATTGAATTTGGCTCAGCAGGCACCTGCCTCAGCTGCTCACTTGAGCCTCTGGGTCTA |
| GCA | TTTC | DNMT1 | LbCas12a | CCTCACTCCTGCTCGGTGAATTT | ACAGCAGGCCTTTGGTCAGGTTGGCTGCTGGGCTGGCCCTGGGGCCGTTTCCCTCACTCCTGC<br>TGCATGAATTTGGCTCAGCAGGCACCTGCCTCAGCTGCTCACTTGAGCCTCTGGGTCTA |
| ATA | TTTC | DNMT1 | LbCas12a | CCTCACTCCTGCTCGGTGAATTT | ACAGCAGGCCTTTGGTCAGGTTGGCTGCTGGGCTGGCCCTGGGGCCGTTTCCCTCACTCCTGC<br>TATATGAATTTGGCTCAGCAGGCACCTGCCTCAGCTGCTCACTTGAGCCTCTGGGTCTA |
| TAC | TTTC | DNMT1 | LbCas12a | CCTCACTCCTGCTCGGTGAATTT | ACAGCAGGCCTTTGGTCAGGTTGGCTGCTGGGCTGGCCCTGGGGCCGTTTCCCTCACTCCTGC<br>TTACTGAATTTGGCTCAGCAGGCACCTGCCTCAGCTGCTCACTTGAGCCTCTGGGTCTA |
| TAG | TTTC | DNMT1 | LbCas12a | CCTCACTCCTGCTCGGTGAATTT | ACAGCAGGCCTTTGGTCAGGTTGGCTGCTGGGCTGGCCCTGGGGCCGTTTCCCTCACTCCTGC<br>TTAGTGAATTTGGCTCAGCAGGCACCTGCCTCAGCTGCTCACTTGAGCCTCTGGGTCTA |
| GAA | TTTC | DNMT1 | LbCas12a | CCTCACTCCTGCTCGGTGAATTT | ACAGCAGGCCTTTGGTCAGGTTGGCTGCTGGGCTGGCCCTGGGGCCGTTTCCCTCACTCCTGC<br>TGAATGAATTTGGCTCAGCAGGCACCTGCCTCAGCTGCTCACTTGAGCCTCTGGGTCTA |
| AAA | TTTC | DNMT1 | LbCas12a | CCTCACTCCTGCTCGGTGAATTT | ACAGCAGGCCTTTGGTCAGGTTGGCTGCTGGGCTGGCCCTGGGGCCGTTTCCCTCACTCCTGC<br>TAAATGAATTTGGCTCAGCAGGCACCTGCCTCAGCTGCTCACTTGAGCCTCTGGGTCTA |
| TCA | TTTC | DNMT1 | LbCas12a | CCTCACTCCTGCTCGGTGAATTT | ACAGCAGGCCTTTGGTCAGGTTGGCTGCTGGGCTGGCCCTGGGGCCGTTTCCCTCACTCCTGC<br>TTCATGAATTTGGCTCAGCAGGCACCTGCCTCAGCTGCTCACTTGAGCCTCTGGGTCTA |
| TTA | TTTC | DNMT1 | LbCas12a | CCTCACTCCTGCTCGGTGAATTT | ACAGCAGGCCTTTGGTCAGGTTGGCTGCTGGGCTGGCCCTGGGGCCGTTTCCCTCACTCCTGC<br>TTTATGAATTTGGCTCAGCAGGCACCTGCCTCAGCTGCTCACTTGAGCCTCTGGGTCTA |
| TAA | TTTC | DNMT1 | LbCas12a | CCTCACTCCTGCTCGGTGAATTT | ACAGCAGGCCTTTGGTCAGGTTGGCTGCTGGGCTGGCCCTGGGGCCGTTTCCCTCACTCCTGC<br>TTAATGAATTTGGCTCAGCAGGCACCTGCCTCAGCTGCTCACTTGAGCCTCTGGGTCTA |
| ACA | TTTC | DNMT1 | LbCas12a | CTGATGGTCCATGTCTGTTACTC | AGCTGTTAACATCAGTACGTTAATGTTTCCTGATGGTCCAACACTGTTACTCGCCTGTCAAGTG<br>GCGTGACACCGGGCGTGTTCCCCAGAGTGACTTTTC |
| GTG | TTTC | DNMT1 | LbCas12a | CTGATGGTCCATGTCTGTTACTC | AGCTGTTAACATCAGTACGTTAATGTTTCCTGATGGTCCAGTGCTGTTACTCGCCTGTCAAGTG<br>GCGTGACACCGGGCGTGTTCCCCAGAGTGACTTTTC |
| CCA | TTTC | DNMT1 | LbCas12a | CTGATGGTCCATGTCTGTTACTC | AGCTGTTAACATCAGTACGTTAATGTTTCCTGATGGTCCACCACTGTTACTCGCCTGTCAAGTG<br>GCGTGACACCGGGCGTGTTCCCCAGAGTGACTTTTC |
| CTG | TTTC | DNMT1 | LbCas12a | CTGATGGTCCATGTCTGTTACTC | AGCTGTTAACATCAGTACGTTAATGTTTCCTGATGGTCCACTGCTGTTACTCGCCTGTCAAGTG<br>GCGTGACACCGGGCGTGTTCCCCAGAGTGACTTTTC |
| AAA | TTTC | DNMT1 | LbCas12a | CTGATGGTCCATGTCTGTTACTC | AGCTGTTAACATCAGTACGTTAATGTTTCCTGATGGTCCAAAACTGTTACTCGCCTGTCAAGTG<br>GCGTGACACCGGGCGTGTTCCCCAGAGTGACTTTTC |
| GAG | TTTC | DNMT1 | LbCas12a | CTGATGGTCCATGTCTGTTACTC | AGCTGTTAACATCAGTACGTTAATGTTTCCTGATGGTCCAGAGCTGTTACTCGCCTGTCAAGTG<br>GCGTGACACCGGGCGTGTTCCCCAGAGTGACTTTTC |
| ACC | TTTC | DNMT1 | LbCas12a | CTGATGGTCCATGTCTGTTACTC | AGCTGTTAACATCAGTACGTTAATGTTTCCTGATGGTCCAACCCTGTTACTCGCCTGTCAAGTG<br>GCGTGACACCGGGCGTGTTCCCCAGAGTGACTTTTC |
| GTC | TTTC | DNMT1 | LbCas12a | CTGATGGTCCATGTCTGTTACTC | AGCTGTTAACATCAGTACGTTAATGTTTCCTGATGGTCCAGTCCTGTTACTCGCCTGTCAAGTG<br>GCGTGACACCGGGCGTGTTCCCCAGAGTGACTTTTC |
| CAA | TTTC | DNMT1 | LbCas12a | CTGATGGTCCATGTCTGTTACTC | AGCTGTTAACATCAGTACGTTAATGTTTCCTGATGGTCCACAACTGTTACTCGCCTGTCAAGTG<br>GCGTGACACCGGGCGTGTTCCCCAGAGTGACTTTTC |
| CAG | TTTC | DNMT1 | LbCas12a | CTGATGGTCCATGTCTGTTACTC | AGCTGTTAACATCAGTACGTTAATGTTTCCTGATGGTCCACAGCTGTTACTCGCCTGTCAAGTG<br>GCGTGACACCGGGCGTGTTCCCCAGAGTGACTTTTC |
| AAC | TTTC | DNMT1 | LbCas12a | CTGATGGTCCATGTCTGTTACTC | AGCTGTTAACATCAGTACGTTAATGTTTCCTGATGGTCCAAACCTGTTACTCGCCTGTCAAGTG<br>GCGTGACACCGGGCGTGTTCCCCAGAGTGACTTTTC |
| GAC | TTTC | DNMT1 | LbCas12a | CTGATGGTCCATGTCTGTTACTC | AGCTGTTAACATCAGTACGTTAATGTTTCCTGATGGTCCAGACCTGTTACTCGCCTGTCAAGTG<br>GCGTGACACCGGGCGTGTTCCCCAGAGTGACTTTTC |
| CCC | TTTC | DNMT1 | LbCas12a | CTGATGGTCCATGTCTGTTACTC | AGCTGTTAACATCAGTACGTTAATGTTTCCTGATGGTCCACCCCTGTTACTCGCCTGTCAAGTG<br>GCGTGACACCGGGCGTGTTCCCCAGAGTGACTTTTC |
| CTC | TTTC | DNMT1 | LbCas12a | CTGATGGTCCATGTCTGTTACTC | AGCTGTTAACATCAGTACGTTAATGTTTCCTGATGGTCCACTCCTGTTACTCGCCTGTCAAGTG<br>GCGTGACACCGGGCGTGTTCCCCAGAGTGACTTTTC |
| CAC | TTTC | DNMT1 | LbCas12a | CTGATGGTCCATGTCTGTTACTC | AGCTGTTAACATCAGTACGTTAATGTTTCCTGATGGTCCACACCTGTTACTCGCCTGTCAAGTG<br>GCGTGACACCGGGCGTGTTCCCCAGAGTGACTTTTC |

| Figure 4d | | | | | | |
|---|---|---|---|---|---|---|
| **# Bases** | **Insertion/Deletion** | **PAM** | **Locus** | **Enzyme** | **Spacer Sequence** | **Extension Sequence** |
| 2 | Deletion | TTTC | DNMT1 | LbCas12a | CCTCACTCCTGCTCGGTGAATTT | ACAGCAGGCCTTTGGTCAGGTTGGCTGCTGGGCTGGCCCTGGGGCCGTTTCCCTCACTCCTGCGGTGAATTTGGCTCAGCAGGCACCTGCCTCAGCTGCTCACTTGAGCCTCTGGGTCTA |
| 3 | Deletion | TTTC | DNMT1 | LbCas12a | CCTCACTCCTGCTCGGTGAATTT | ACAGCAGGCCTTTGGTCAGGTTGGCTGCTGGGCTGGCCCTGGGGCCGTTTCCCTCACTCCTCGGTGAATTTGGCTCAGCAGGCACCTGCCTCAGCTGCTCACTTGAGCCTCTGGGTCTA |
| 4 | Deletion | TTTC | DNMT1 | LbCas12a | CCTCACTCCTGCTCGGTGAATTT | ACAGCAGGCCTTTGGTCAGGTTGGCTGCTGGGCTGGCCCTGGGGCCGTTTCCCTCACTCCTGGTGAATTTGGCTCAGCAGGCACCTGCCTCAGCTGCTCACTTGAGCCTCTGGGTCTA |
| 6 | Deletion | TTTC | DNMT1 | LbCas12a | CCTCACTCCTGCTCGGTGAATTT | ACAGCAGGCCTTTGGTCAGGTTGGCTGCTGGGCTGGCCCTGGGGCCGTTTCCCTCACTCCGTGAATTTGGCTCAGCAGGCACCTGCCTCAGCTGCTCACTTGAGCCTCTGGGTCTA |
| 8 | Deletion | TTTC | DNMT1 | LbCas12a | CCTCACTCCTGCTCGGTGAATTT | ACAGCAGGCCTTTGGTCAGGTTGGCTGCTGGGCTGGCCCTGGGGCCGTTTCCCTCACTCTGAATTTGGCTCAGCAGGCACCTGCCTCAGCTGCTCACTTGAGCCTCTGGGTCTA |
| 11 | Deletion | TTTC | DNMT1 | LbCas12a | CCTCACTCCTGCTCGGTGAATTT | ACAGCAGGCCTTTGGTCAGGTTGGCTGCTGGGCTGGCCCTGGGGCCGTTTCCCTCACTCCTTTGGCTCAGCAGGCACCTGCCTCAGCTGCTCACTTGAGCCTCTGGGTCTA |
| 14 | Deletion | TTTC | DNMT1 | LbCas12a | CCTCACTCCTGCTCGGTGAATTT | ACAGCAGGCCTTTGGTCAGGTTGGCTGCTGGGCTGGCCCTGGGGCCGTTTCCCTCACTCCGGCTCAGCAGGCACCTGCCTCAGCTGCTCACTTGAGCCTCTGGGTCTA |
| 20 | Deletion | TTTC | DNMT1 | LbCas12a | CCTCACTCCTGCTCGGTGAATTT | ACAGCAGGCCTTTGGTCAGGTTGGCTGCTGGGCTGGCCCTGGGGCCGTTTCCCTGGCTCAGCAGGCACCTGCCTCAGCTGCTCACTTGAGCCTCTGGGTCTA |
| 30 | Deletion | TTTC | DNMT1 | LbCas12a | CCTCACTCCTGCTCGGTGAATTT | ACAGCAGGCCTTTGGTCAGGTTGGCTGCTGGGCTGGCCCTGGGGGGCTCAGCAGGCACCTGCCTCAGCTGCTCACTTGAGCCTCTGGGTCTA |
| 3 | Insertion | TTTC | DNMT1 | LbCas12a | CCTCACTCCTGCTCGGTGAATTT | ACAGCAGGCCTTTGGTCAGGTTGGCTGCTGGGCTGGCCCTGGGGCCGTTTCCCTCACTCCTGCTCGACGGTGAATTTGGCTCAGCAGGCACCTGCCTCAGCTGCTCACTTGAGCCTCTGGGTCTA |
| 6 | Insertion | TTTC | DNMT1 | LbCas12a | CCTCACTCCTGCTCGGTGAATTT | ACAGCAGGCCTTTGGTCAGGTTGGCTGCTGGGCTGGCCCTGGGGCCGTTTCCCTCACTCCTGCTCGACTACGGTGAATTTGGCTCAGCAGGCACCTGCCTCAGCTGCTCACTTGAGCCTCTGGGTCTA |
| 9 | Insertion | TTTC | DNMT1 | LbCas12a | CCTCACTCCTGCTCGGTGAATTT | ACAGCAGGCCTTTGGTCAGGTTGGCTGCTGGGCTGGCCCTGGGGCCGTTTCCCTCACTCCTGCTCGACTACAAAGGTGAATTTGGCTCAGCAGGCACCTGCCTCAGCTGCTCACTTGAGCCTCTGGGTCTA |
| 12 | Insertion | TTTC | DNMT1 | LbCas12a | CCTCACTCCTGCTCGGTGAATTT | ACAGCAGGCCTTTGGTCAGGTTGGCTGCTGGGCTGGCCCTGGGGCCGTTTCCCTCACTCCTGCTCGACTACAAAGACGGTGAATTTGGCTCAGCAGGCACCTGCCTCAGCTGCTCACTTGAGCCTCTGGGTCTA |
| 15 | Insertion | TTTC | DNMT1 | LbCas12a | CCTCACTCCTGCTCGGTGAATTT | ACAGCAGGCCTTTGGTCAGGTTGGCTGCTGGGCTGGCCCTGGGGCCGTTTCCCTCACTCCTGCTCGACTACAAAGACGATGGTGAATTTGGCTCAGCAGGCACCTGCCTCAGCTGCTCACTTGAGCCTCTGGGTCTA |
| 18 | Insertion | TTTC | DNMT1 | LbCas12a | CCTCACTCCTGCTCGGTGAATTT | ACAGCAGGCCTTTGGTCAGGTTGGCTGCTGGGCTGGCCCTGGGGCCGTTTCCCTCACTCCTGCTCGACTACAAAGACGATGACGGTGAATTTGGCTCAGCAGGCACCTGCCTCAGCTGCTCACTTGAGCCTCTGGGTCTA |
| 21 | Insertion | TTTC | DNMT1 | LbCas12a | CCTCACTCCTGCTCGGTGAATTT | ACAGCAGGCCTTTGGTCAGGTTGGCTGCTGGGCTGGCCCTGGGGCCGTTTCCCTCACTCCTGCTCGACTACAAAGACGATGACGACGGTGAATTTGGCTCAGCAGGCACCTGCCTCAGCTGCTCACTTGAGCCTCTGGGTCTA |
| 24 | Insertion | TTTC | DNMT1 | LbCas12a | CCTCACTCCTGCTCGGTGAATTT | ACAGCAGGCCTTTGGTCAGGTTGGCTGCTGGGCTGGCCCTGGGGCCGTTTCCCTCACTCCTGCTCGACTACAAAGACGATGACGACAAGGGTGAATTTGGCTCAGCAGGCACCTGCCTCAGCTGCTCACTTGAGCCTCTGGGTCTA |

| Figure 6a | | | | |
|---|---|---|---|---|
| **PAM** | **Locus** | **Enzyme** | **Spacer Sequence** | **Extension Sequence** |
| GGG | EMX1 | Cas9 | GAGTCCGAGCAGAAGAAGAA | GTGATGGGAGCACTTCTTCTTCTGCTCGGA |
| TGG | FANCF | Cas9 | GGAATCCCTTCTGCAGCACC | GGAAAAGCGATCAAGGTGCTGCAGAAGGGA |
| AGG | RNF2 | Cas9 | GTCATCTTAGTCATTACCTG | AACGAACACCTCATGTAATGACTAAGATG |

| Figure 5b | | | | |
|---|---|---|---|---|
| **Modification (position)** | **Spacer Sequence** | **PAM** | **Locus** | **Enzyme** |
| N/A | CCTCACTCCTGCTCGGTGAATTT | TTTC | DNMT1 | LbCas12a |
| P12,P13 | CCTCACTCCTGAGCGGTGAATTT | TTTC | DNMT1 | LbCas12a |
| P13,P14 | CCTCACTCCTGCGAGGTGAATTT | TTTC | DNMT1 | LbCas12a |
| P14,P15 | CCTCACTCCTGCTATGTGAATTT | TTTC | DNMT1 | LbCas12a |
| N/A | GCTCAGCAGGCACCTGCCTCAGC | TTTG | DNMT1 | LbCas12a |
| P12,P13 | GCTCAGCAGGCCACTGCCTCAGC | TTTG | DNMT1 | LbCas12a |
| P13,P14 | GCTCAGCAGGCAAATGCCTCAGC | TTTG | DNMT1 | LbCas12a |
| P14,P15 | GCTCAGCAGGCACAGGCCTCAGC | TTTG | DNMT1 | LbCas12a |
| N/A | GACCAATAGCATTGCAGAGAGGC | TTTC | FANCF | LbCas12a |
| P12,P13 | GACCAATAGCAGGGCAGAGAGGC | TTTC | FANCF | LbCas12a |
| P13,P14 | GACCAATAGCATGTCAGAGAGGC | TTTC | FANCF | LbCas12a |
| P14,P15 | GACCAATAGCATTTAAGAGAGGC | TTTC | FANCF | LbCas12a |

Supplementary Figure 2b

| PBS | RTT | 5' or 3' | PAM | Locus | Enzyme | Spacer Sequence | Extension Sequence |
|---|---|---|---|---|---|---|---|
| 48 | 74 | 3' | TTTC | FANCF | LbCas12a | GCGGATGTTCCAATCAGTACGCA | GAGATTGACATGCATTTCGACCAATAGCATTGCAGAGAGGCGTATCATAACGCGGATGTTCCAATCAGTACGCAGAGAGTCGCCGTCTCCAAGGTGAAAGCGGAAGTAGGGCCTTCGCGCAC |
| 48 | 52 | 3' | TTTC | FANCF | LbCas12a | GCGGATGTTCCAATCAGTACGCA | AATAGCATTGCAGAGAGGCGTATCATAACGCGGATGTTCCAATCAGTACGCAGAGAGTCGCCGTCTCCAAGGTGAAAGCGGAAGTAGGGCCTTCGCGCAC |
| 48 | 44 | 3' | TTTC | FANCF | LbCas12a | GCGGATGTTCCAATCAGTACGCA | TGCAGAGAGGCGTATCATAACGCGGATGTTCCAATCAGTACGCAGAGAGTCGCCGTCTCCAAGGTGAAAGCGGAAGTAGGGCCTTCGCGCAC |
| 48 | 36 | 3' | TTTC | FANCF | LbCas12a | GCGGATGTTCCAATCAGTACGCA | GGCGTATCATAACGCGGATGTTCCAATCAGTACGCAGAGAGTCGCCGTCTCCAAGGTGAAAGCGGAAGTAGGGCCTTCGCGCAC |
| 24 | 74 | 3' | TTTC | FANCF | LbCas12a | GCGGATGTTCCAATCAGTACGCA | GAGATTGACATGCATTTCGACCAATAGCATTGCAGAGAGGCGTATCATAACGCGGATGTTCCAATCAGTACGCAGAGAGTCGCCGTCTCCAAGGTGAA |
| 24 | 52 | 3' | TTTC | FANCF | LbCas12a | GCGGATGTTCCAATCAGTACGCA | AATAGCATTGCAGAGAGGCGTATCATAACGCGGATGTTCCAATCAGTACGCAGAGAGTCGCCGTCTCCAAGGTGAA |
| 24 | 44 | 3' | TTTC | FANCF | LbCas12a | GCGGATGTTCCAATCAGTACGCA | TGCAGAGAGGCGTATCATAACGCGGATGTTCCAATCAGTACGCAGAGAGTCGCCGTCTCCAAGGTGAA |
| 24 | 36 | 3' | TTTC | FANCF | LbCas12a | GCGGATGTTCCAATCAGTACGCA | GGCGTATCATAACGCGGATGTTCCAATCAGTACGCAGAGAGTCGCCGTCTCCAAGGTGAA |
| 16 | 74 | 3' | TTTC | FANCF | LbCas12a | GCGGATGTTCCAATCAGTACGCA | GAGATTGACATGCATTTCGACCAATAGCATTGCAGAGAGGCGTATCATAACGCGGATGTTCCAATCAGTACGCAGAGAGTCGCCGTCTCC |
| 16 | 52 | 3' | TTTC | FANCF | LbCas12a | GCGGATGTTCCAATCAGTACGCA | AATAGCATTGCAGAGAGGCGTATCATAACGCGGATGTTCCAATCAGTACGCAGAGAGTCGCCGTCTCC |
| 16 | 44 | 3' | TTTC | FANCF | LbCas12a | GCGGATGTTCCAATCAGTACGCA | TGCAGAGAGGCGTATCATAACGCGGATGTTCCAATCAGTACGCAGAGAGTCGCCGTCTCC |
| 16 | 36 | 3' | TTTC | FANCF | LbCas12a | GCGGATGTTCCAATCAGTACGCA | GGCGTATCATAACGCGGATGTTCCAATCAGTACGCAGAGAGTCGCCGTCTCC |
| 8 | 74 | 3' | TTTC | FANCF | LbCas12a | GCGGATGTTCCAATCAGTACGCA | GAGATTGACATGCATTTCGACCAATAGCATTGCAGAGAGGCGTATCATAACGCGGATGTTCCAATCAGTACGCAGAGAGTCG |
| 8 | 52 | 3' | TTTC | FANCF | LbCas12a | GCGGATGTTCCAATCAGTACGCA | AATAGCATTGCAGAGAGGCGTATCATAACGCGGATGTTCCAATCAGTACGCAGAGAGTCG |
| 8 | 44 | 3' | TTTC | FANCF | LbCas12a | GCGGATGTTCCAATCAGTACGCA | TGCAGAGAGGCGTATCATAACGCGGATGTTCCAATCAGTACGCAGAGAGTCG |
| 8 | 36 | 3' | TTTC | FANCF | LbCas12a | GCGGATGTTCCAATCAGTACGCA | GGCGTATCATAACGCGGATGTTCCAATCAGTACGCAGAGAGTCG |
| 48 | 74 | 5' | TTTC | FANCF | LbCas12a | GCGGATGTTCCAATCAGTACGCA | GAGATTGACATGCATTTCGACCAATAGCATTGCAGAGAGGCGTATCATAACGCGGATGTTCCAATCAGTACGCAGAGAGTCGCCGTCTCCAAGGTGAAAGCGGAAGTAGGGCCTTCGCGCAC |
| 48 | 52 | 5' | TTTC | FANCF | LbCas12a | GCGGATGTTCCAATCAGTACGCA | AATAGCATTGCAGAGAGGCGTATCATAACGCGGATGTTCCAATCAGTACGCAGAGAGTCGCCGTCTCCAAGGTGAAAGCGGAAGTAGGGCCTTCGCGCAC |
| 48 | 44 | 5' | TTTC | FANCF | LbCas12a | GCGGATGTTCCAATCAGTACGCA | TGCAGAGAGGCGTATCATAACGCGGATGTTCCAATCAGTACGCAGAGAGTCGCCGTCTCCAAGGTGAAAGCGGAAGTAGGGCCTTCGCGCAC |
| 48 | 36 | 5' | TTTC | FANCF | LbCas12a | GCGGATGTTCCAATCAGTACGCA | GGCGTATCATAACGCGGATGTTCCAATCAGTACGCAGAGAGTCGCCGTCTCCAAGGTGAAAGCGGAAGTAGGGCCTTCGCGCAC |
| 24 | 74 | 5' | TTTC | FANCF | LbCas12a | GCGGATGTTCCAATCAGTACGCA | GAGATTGACATGCATTTCGACCAATAGCATTGCAGAGAGGCGTATCATAACGCGGATGTTCCAATCAGTACGCAGAGAGTCGCCGTCTCCAAGGTGAA |
| 24 | 52 | 5' | TTTC | FANCF | LbCas12a | GCGGATGTTCCAATCAGTACGCA | AATAGCATTGCAGAGAGGCGTATCATAACGCGGATGTTCCAATCAGTACGCAGAGAGTCGCCGTCTCCAAGGTGAA |
| 24 | 44 | 5' | TTTC | FANCF | LbCas12a | GCGGATGTTCCAATCAGTACGCA | TGCAGAGAGGCGTATCATAACGCGGATGTTCCAATCAGTACGCAGAGAGTCGCCGTCTCCAAGGTGAA |
| 24 | 36 | 5' | TTTC | FANCF | LbCas12a | GCGGATGTTCCAATCAGTACGCA | GGCGTATCATAACGCGGATGTTCCAATCAGTACGCAGAGAGTCGCCGTCTCCAAGGTGAA |
| 0 | 0 | N/A | TTTC | DNMT1 | LbCas12a | CTGATGGTCCATGTCTGTTACTC | N/A |
| 80 | 36 | 3' | TTTC | DNMT1 | LbCas12a | CTGATGGTCCATGTCTGTTACTC | ACGTTAATGTAACCTGATGGTCCATGTCTGTTACTCGCCTGTCAAGTGGCGTGACACCGGGCGTGTTCCCCAGAGTGACTTTTCCTTTTATTTCCCTTCAGCTAAAATAAAGGAGG |
| 72 | 36 | 3' | TTTC | DNMT1 | LbCas12a | CTGATGGTCCATGTCTGTTACTC | ACGTTAATGTAACCTGATGGTCCATGTCTGTTACTCGCCTGTCAAGTGGCGTGACACCGGGCGTGTTCCCCAGAGTGACTTTTCCTTTTATTTCCCTTCAGCTAAAAT |
| 64 | 36 | 3' | TTTC | DNMT1 | LbCas12a | CTGATGGTCCATGTCTGTTACTC | ACGTTAATGTAACCTGATGGTCCATGTCTGTTACTCGCCTGTCAAGTGGCGTGACACCGGGCGTGTTCCCCAGAGTGACTTTTCCTTTTATTTCCCTTCA |
| 56 | 36 | 3' | TTTC | DNMT1 | LbCas12a | CTGATGGTCCATGTCTGTTACTC | ACGTTAATGTAACCTGATGGTCCATGTCTGTTACTCGCCTGTCAAGTGGCGTGACACCGGGCGTGTTCCCCAGAGTGACTTTTCCTTTTATT |
| 48 | 36 | 3' | TTTC | DNMT1 | LbCas12a | CTGATGGTCCATGTCTGTTACTC | ACGTTAATGTAACCTGATGGTCCATGTCTGTTACTCGCCTGTCAAGTGGCGTGACACCGGGCGTGTTCCCCAGAGTGACTTTTC |
| 40 | 36 | 3' | TTTC | DNMT1 | LbCas12a | CTGATGGTCCATGTCTGTTACTC | ACGTTAATGTAACCTGATGGTCCATGTCTGTTACTCGCCTGTCAAGTGGCGTGACACCGGGCGTGTTCCCCAGAGT |
| 32 | 36 | 3' | TTTC | DNMT1 | LbCas12a | CTGATGGTCCATGTCTGTTACTC | ACGTTAATGTAACCTGATGGTCCATGTCTGTTACTCGCCTGTCAAGTGGCGTGACACCGGGCGTGTTC |
| 24 | 36 | 3' | TTTC | DNMT1 | LbCas12a | CTGATGGTCCATGTCTGTTACTC | ACGTTAATGTAACCTGATGGTCCATGTCTGTTACTCGCCTGTCAAGTGGCGTGACACCGG |
| 16 | 36 | 3' | TTTC | DNMT1 | LbCas12a | CTGATGGTCCATGTCTGTTACTC | ACGTTAATGTAACCTGATGGTCCATGTCTGTTACTCGCCTGTCAAGTGGCGT |
| 8 | 36 | 3' | TTTC | DNMT1 | LbCas12a | CTGATGGTCCATGTCTGTTACTC | ACGTTAATGTAACCTGATGGTCCATGTCTGTTACTCGCCTGTCA |
| 48 | 74 | 3' | TTTC | DNMT1 | LbCas12a | CTGATGGTCCATGTCTGTTACTC | GCACTCTGCCACTTATTGGGTCAGCTGTTAACATCAGTACGTTAATGTAACCTGATGGTCCATGTCTGTTACTCGCCTGTCAAGTGGCGTGACACCGGGCGTGTTCCCCAGAGTGACTTTTC |
| 48 | 52 | 3' | TTTC | DNMT1 | LbCas12a | CTGATGGTCCATGTCTGTTACTC | AGCTGTTAACATCAGTACGTTAATGTAACCTGATGGTCCATGTCTGTTACTCGCCTGTCAAGTGGCGTGACACCGGGCGTGTTCCCCAGAGTGACTTTTC |
| 48 | 44 | 3' | TTTC | DNMT1 | LbCas12a | CTGATGGTCCATGTCTGTTACTC | ACATCAGTACGTTAATGTAACCTGATGGTCCATGTCTGTTACTCGCCTGTCAAGTGGCGTGACACCGGGCGTGTTCCCCAGAGTGACTTTTC |
| 48 | 40 | 3' | TTTC | DNMT1 | LbCas12a | CTGATGGTCCATGTCTGTTACTC | CAGTACGTTAATGTAACCTGATGGTCCATGTCTGTTACTCGCCTGTCAAGTGGCGTGACACCGGGCGTGTTCCCCAGAGTGACTTTTC |
| 48 | 32 | 3' | TTTC | DNMT1 | LbCas12a | CTGATGGTCCATGTCTGTTACTC | TAATGTAACCTGATGGTCCATGTCTGTTACTCGCCTGTCAAGTGGCGTGACACCGGGCGTGTTCCCCAGAGTGACTTTTC |
| 48 | 28 | 3' | TTTC | DNMT1 | LbCas12a | CTGATGGTCCATGTCTGTTACTC | GTAACCTGATGGTCCATGTCTGTTACTCGCCTGTCAAGTGGCGTGACACCGGGCGTGTTCCCCAGAGTGACTTTTC |
| 80 | 36 | 5' | TTTC | DNMT1 | LbCas12a | CTGATGGTCCATGTCTGTTACTC | ACGTTAATGTAACCTGATGGTCCATGTCTGTTACTCGCCTGTCAAGTGGCGTGACACCGGGCGTGTTCCCCAGAGTGACTTTTCCTTTTATTTCCCTTCAGCTAAAATAAAGGAGG |
| 72 | 36 | 5' | TTTC | DNMT1 | LbCas12a | CTGATGGTCCATGTCTGTTACTC | ACGTTAATGTAACCTGATGGTCCATGTCTGTTACTCGCCTGTCAAGTGGCGTGACACCGGGCGTGTTCCCCAGAGTGACTTTTCCTTTTATTTCCCTTCAGCTAAAAT |
| 64 | 36 | 5' | TTTC | DNMT1 | LbCas12a | CTGATGGTCCATGTCTGTTACTC | ACGTTAATGTAACCTGATGGTCCATGTCTGTTACTCGCCTGTCAAGTGGCGTGACACCGGGCGTGTTCCCCAGAGTGACTTTTCCTTTTATTTCCCTTCA |
| 56 | 36 | 5' | TTTC | DNMT1 | LbCas12a | CTGATGGTCCATGTCTGTTACTC | ACGTTAATGTAACCTGATGGTCCATGTCTGTTACTCGCCTGTCAAGTGGCGTGACACCGGGCGTGTTCCCCAGAGTGACTTTTCCTTTTATT |
| 48 | 36 | 5' | TTTC | DNMT1 | LbCas12a | CTGATGGTCCATGTCTGTTACTC | ACGTTAATGTAACCTGATGGTCCATGTCTGTTACTCGCCTGTCAAGTGGCGTGACACCGGGCGTGTTCCCCAGAGTGACTTTTC |
| 40 | 36 | 5' | TTTC | DNMT1 | LbCas12a | CTGATGGTCCATGTCTGTTACTC | ACGTTAATGTAACCTGATGGTCCATGTCTGTTACTCGCCTGTCAAGTGGCGTGACACCGGGCGTGTTCCCCAGAGT |
| 32 | 36 | 5' | TTTC | DNMT1 | LbCas12a | CTGATGGTCCATGTCTGTTACTC | ACGTTAATGTAACCTGATGGTCCATGTCTGTTACTCGCCTGTCAAGTGGCGTGACACCGGGCGTGTTC |
| 24 | 36 | 5' | TTTC | DNMT1 | LbCas12a | CTGATGGTCCATGTCTGTTACTC | ACGTTAATGTAACCTGATGGTCCATGTCTGTTACTCGCCTGTCAAGTGGCGTGACACCGG |
| 16 | 36 | 5' | TTTC | DNMT1 | LbCas12a | CTGATGGTCCATGTCTGTTACTC | ACGTTAATGTAACCTGATGGTCCATGTCTGTTACTCGCCTGTCAAGTGGCGT |
| 8 | 36 | 5' | TTTC | DNMT1 | LbCas12a | CTGATGGTCCATGTCTGTTACTC | ACGTTAATGTAACCTGATGGTCCATGTCTGTTACTCGCCTGTCA |
| 48 | 74 | 5' | TTTC | DNMT1 | LbCas12a | CTGATGGTCCATGTCTGTTACTC | GCACTCTGCCACTTATTGGGTCAGCTGTTAACATCAGTACGTTAATGTAACCTGATGGTCCATGTCTGTTACTCGCCTGTCAAGTGGCGTGACACCGGGCGTGTTCCCCAGAGTGACTTTTC |
| 48 | 52 | 5' | TTTC | DNMT1 | LbCas12a | CTGATGGTCCATGTCTGTTACTC | AGCTGTTAACATCAGTACGTTAATGTAACCTGATGGTCCATGTCTGTTACTCGCCTGTCAAGTGGCGTGACACCGGGCGTGTTCCCCAGAGTGACTTTTC |
| 48 | 44 | 5' | TTTC | DNMT1 | LbCas12a | CTGATGGTCCATGTCTGTTACTC | ACATCAGTACGTTAATGTAACCTGATGGTCCATGTCTGTTACTCGCCTGTCAAGTGGCGTGACACCGGGCGTGTTCCCCAGAGTGACTTTTC |
| 48 | 40 | 5' | TTTC | DNMT1 | LbCas12a | CTGATGGTCCATGTCTGTTACTC | CAGTACGTTAATGTAACCTGATGGTCCATGTCTGTTACTCGCCTGTCAAGTGGCGTGACACCGGGCGTGTTCCCCAGAGTGACTTTTC |
| 48 | 32 | 5' | TTTC | DNMT1 | LbCas12a | CTGATGGTCCATGTCTGTTACTC | TAATGTAACCTGATGGTCCATGTCTGTTACTCGCCTGTCAAGTGGCGTGACACCGGGCGTGTTCCCCAGAGTGACTTTTC |
| 48 | 28 | 5' | TTTC | DNMT1 | LbCas12a | CTGATGGTCCATGTCTGTTACTC | GTAACCTGATGGTCCATGTCTGTTACTCGCCTGTCAAGTGGCGTGACACCGGGCGTGTTCCCCAGAGTGACTTTTC |

Supplementary Figure 4a

| Mutation Position | PAM | Locus | Enzyme | Spacer Sequence | Extension Sequence |
|---|---|---|---|---|---|
| P13,P14 | TTTC | DNMT1 | LbCas12a | CCTCACTCCTGCTCGGTGAATTT | ACAGCAGGCCTTTGGTCAGGTTGGCTGCTGGGCTGGCCCTGGGGCCGTTTCCCTCACTCCTGCAGGGTGAATTTGGC<br>TCAGCAGGCACCTGCCTCAGCTGCTCACTTGAGCCTCTGGGTCTA |
| P13,P14 | TTTC | DNMT1 | LbCas12a | CCTCACTCCTGCTCGGTGAATTT | ACAGCAGGCCTTTGGTCAGGTTGGCTGCTGGGCTGGCCCTGGGGCCGTTTCCCTCACTCCTGCCTGGTGAATTTGGC<br>TCAGCAGGCACCTGCCTCAGCTGCTCACTTGAGCCTCTGGGTCTA |
| P13,P14 | TTTC | DNMT1 | LbCas12a | CCTCACTCCTGCTCGGTGAATTT | ACAGCAGGCCTTTGGTCAGGTTGGCTGCTGGGCTGGCCCTGGGGCCGTTTCCCTCACTCCTGCGAGGTGAATTTGGC<br>TCAGCAGGCACCTGCCTCAGCTGCTCACTTGAGCCTCTGGGTCTA |
| P12,P13,P14 | TTTC | DNMT1 | LbCas12a | CCTCACTCCTGCTCGGTGAATTT | ACAGCAGGCCTTTGGTCAGGTTGGCTGCTGGGCTGGCCCTGGGGCCGTTTCCCTCACTCCTGGAGGGTGAATTTGG<br>CTCAGCAGGCACCTGCCTCAGCTGCTCACTTGAGCCTCTGGGTCTA |
| P13,P14,P15 | TTTC | DNMT1 | LbCas12a | CCTCACTCCTGCTCGGTGAATTT | ACAGCAGGCCTTTGGTCAGGTTGGCTGCTGGGCTGGCCCTGGGGCCGTTTCCCTCACTCCTGCAGCGTGAATTTGGC<br>TCAGCAGGCACCTGCCTCAGCTGCTCACTTGAGCCTCTGGGTCTA |
| P12,P13,P14,P15 | TTTC | DNMT1 | LbCas12a | CCTCACTCCTGCTCGGTGAATTT | ACAGCAGGCCTTTGGTCAGGTTGGCTGCTGGGCTGGCCCTGGGGCCGTTTCCCTCACTCCTGGAGCGTGAATTTGGC<br>TCAGCAGGCACCTGCCTCAGCTGCTCACTTGAGCCTCTGGGTCTA |
| P12 | TTTC | DNMT1 | LbCas12a | CCTCACTCCTGCTCGGTGAATTT | ACAGCAGGCCTTTGGTCAGGTTGGCTGCTGGGCTGGCCCTGGGGCCGTTTCCCTCACTCCTGGTCGGTGAATTTGGC<br>TCAGCAGGCACCTGCCTCAGCTGCTCACTTGAGCCTCTGGGTCTA |
| P13 | TTTC | DNMT1 | LbCas12a | CCTCACTCCTGCTCGGTGAATTT | ACAGCAGGCCTTTGGTCAGGTTGGCTGCTGGGCTGGCCCTGGGGCCGTTTCCCTCACTCCTGCACGGTGAATTTGGC<br>TCAGCAGGCACCTGCCTCAGCTGCTCACTTGAGCCTCTGGGTCTA |
| P14 | TTTC | DNMT1 | LbCas12a | CCTCACTCCTGCTCGGTGAATTT | ACAGCAGGCCTTTGGTCAGGTTGGCTGCTGGGCTGGCCCTGGGGCCGTTTCCCTCACTCCTGCTGGGTGAATTTGGC<br>TCAGCAGGCACCTGCCTCAGCTGCTCACTTGAGCCTCTGGGTCTA |
| P15 | TTTC | DNMT1 | LbCas12a | CCTCACTCCTGCTCGGTGAATTT | ACAGCAGGCCTTTGGTCAGGTTGGCTGCTGGGCTGGCCCTGGGGCCGTTTCCCTCACTCCTGCTCCGTGAATTTGGC<br>TCAGCAGGCACCTGCCTCAGCTGCTCACTTGAGCCTCTGGGTCTA |
| P13,P14 | TTTC | FANCF | LbCas12a | GCGGATGTTCCAATCAGTACGCA | AATAGCATTGCAGAGAGGCGTATCATTTCGCGGATGTTCCATACAGTACGCAGAGAGTCGCCGTCTCCAAGGTGAA<br>AGCGGAAGTAGGGCCTTCGCGCAC |
| P13,P14 | TTTC | FANCF | LbCas12a | GCGGATGTTCCAATCAGTACGCA | AATAGCATTGCAGAGAGGCGTATCATTTCGCGGATGTTCCAGCCAGTACGCAGAGAGTCGCCGTCTCCAAGGTGAA<br>AGCGGAAGTAGGGCCTTCGCGCAC |
| P13,P14 | TTTC | FANCF | LbCas12a | GCGGATGTTCCAATCAGTACGCA | AATAGCATTGCAGAGAGGCGTATCATTTCGCGGATGTTCCACGCAGTACGCAGAGAGTCGCCGTCTCCAAGGTGAA<br>AGCGGAAGTAGGGCCTTCGCGCAC |
| P12,P13,P14 | TTTC | FANCF | LbCas12a | GCGGATGTTCCAATCAGTACGCA | AATAGCATTGCAGAGAGGCGTATCATTTCGCGGATGTTCCTTACAGTACGCAGAGAGTCGCCGTCTCCAAGGTGAAA<br>GCGGAAGTAGGGCCTTCGCGCAC |
| P13,P14,P15 | TTTC | FANCF | LbCas12a | GCGGATGTTCCAATCAGTACGCA | AATAGCATTGCAGAGAGGCGTATCATTTCGCGGATGTTCCATAGAGTACGCAGAGAGTCGCCGTCTCCAAGGTGAA<br>AGCGGAAGTAGGGCCTTCGCGCAC |
| P12,P13,P14,P15 | TTTC | FANCF | LbCas12a | GCGGATGTTCCAATCAGTACGCA | AATAGCATTGCAGAGAGGCGTATCATTTCGCGGATGTTCCTTAGAGTACGCAGAGAGTCGCCGTCTCCAAGGTGAA<br>AGCGGAAGTAGGGCCTTCGCGCAC |
| P12 | TTTC | FANCF | LbCas12a | GCGGATGTTCCAATCAGTACGCA | AATAGCATTGCAGAGAGGCGTATCATTTCGCGGATGTTCCTATCAGTACGCAGAGAGTCGCCGTCTCCAAGGTGAAA<br>GCGGAAGTAGGGCCTTCGCGCAC |
| P13 | TTTC | FANCF | LbCas12a | GCGGATGTTCCAATCAGTACGCA | AATAGCATTGCAGAGAGGCGTATCATTTCGCGGATGTTCCATTCAGTACGCAGAGAGTCGCCGTCTCCAAGGTGAAA<br>GCGGAAGTAGGGCCTTCGCGCAC |
| P14 | TTTC | FANCF | LbCas12a | GCGGATGTTCCAATCAGTACGCA | AATAGCATTGCAGAGAGGCGTATCATTTCGCGGATGTTCCAAACAGTACGCAGAGAGTCGCCGTCTCCAAGGTGAA<br>AGCGGAAGTAGGGCCTTCGCGCAC |
| P15 | TTTC | FANCF | LbCas12a | GCGGATGTTCCAATCAGTACGCA | AATAGCATTGCAGAGAGGCGTATCATTTCGCGGATGTTCCAATGAGTACGCAGAGAGTCGCCGTCTCCAAGGTGAA<br>AGCGGAAGTAGGGCCTTCGCGCAC |

Supplementary Figure 4b

| # Bases | Insertion/Deletion | PAM | Locus | Enzyme | Spacer Sequence | Extension Sequence |
|---|---|---|---|---|---|---|
| 2 | Deletion | TTTC | FANCF | LbCas12a | GCGGATGTTCCAATCAGTACGCA | AATAGCATTGCAGAGAGGCGTATCATTTCGCGGATGTTCCACAGTACGCAGAGAGTCGCCGTCTCCAAG<br>GTGAAAGCGGAAGTAGGGCCTTCGCGCAC |
| 3 | Deletion | TTTC | FANCF | LbCas12a | GCGGATGTTCCAATCAGTACGCA | AATAGCATTGCAGAGAGGCGTATCATTTCGCGGATGTTCCAAGTACGCAGAGAGTCGCCGTCTCCAAGGT<br>GAAAGCGGAAGTAGGGCCTTCGCGCAC |
| 4 | Deletion | TTTC | FANCF | LbCas12a | GCGGATGTTCCAATCAGTACGCA | AATAGCATTGCAGAGAGGCGTATCATTTCGCGGATGTTCCAGTACGCAGAGAGTCGCCGTCTCCAAGGT<br>GAAAGCGGAAGTAGGGCCTTCGCGCAC |
| 6 | Deletion | TTTC | FANCF | LbCas12a | GCGGATGTTCCAATCAGTACGCA | AATAGCATTGCAGAGAGGCGTATCATTTCGCGGATGTTCGTACGCAGAGAGTCGCCGTCTCCAAGGTGA<br>AAGCGGAAGTAGGGCCTTCGCGCAC |
| 8 | Deletion | TTTC | FANCF | LbCas12a | GCGGATGTTCCAATCAGTACGCA | AATAGCATTGCAGAGAGGCGTATCATTTCGCGGATGTTTACGCAGAGAGTCGCCGTCTCCAAGGTGAAA<br>GCGGAAGTAGGGCCTTCGCGCAC |
| 11 | Deletion | TTTC | FANCF | LbCas12a | GCGGATGTTCCAATCAGTACGCA | AATAGCATTGCAGAGAGGCGTATCATTTCGCGGATGTTGCAGAGAGTCGCCGTCTCCAAGGTGAAAGCG<br>GAAGTAGGGCCTTCGCGCAC |
| 14 | Deletion | TTTC | FANCF | LbCas12a | GCGGATGTTCCAATCAGTACGCA | AATAGCATTGCAGAGAGGCGTATCATTTCGCGGATGTTGAGAGTCGCCGTCTCCAAGGTGAAAGCGGAA<br>GTAGGGCCTTCGCGCAC |
| 20 | Deletion | TTTC | FANCF | LbCas12a | GCGGATGTTCCAATCAGTACGCA | AATAGCATTGCAGAGAGGCGTATCATTTCGCGGAGAGTCGCCGTCTCCAAGGTGAAAGCGGAAGTAGGG<br>CCTTCGCGCAC |
| 30 | Deletion | TTTC | FANCF | LbCas12a | GCGGATGTTCCAATCAGTACGCA | AATAGCATTGCAGAGAGGCGTAGAGAGTCGCCGTCTCCAAGGTGAAAGCGGAAGTAGGGCCTTCGCGC<br>AC |
| 3 | Insertion | TTTC | FANCF | LbCas12a | GCGGATGTTCCAATCAGTACGCA | AATAGCATTGCAGAGAGGCGTATCATTTCGCGGATGTTCCAAGACTCAGTACGCAGAGAGTCGCCGTCTC<br>CAAGGTGAAAGCGGAAGTAGGGCCTTCGCGCAC |
| 6 | Insertion | TTTC | FANCF | LbCas12a | GCGGATGTTCCAATCAGTACGCA | AATAGCATTGCAGAGAGGCGTATCATTTCGCGGATGTTCCAAGACTACTCAGTACGCAGAGAGTCGCCGT<br>CTCCAAGGTGAAAGCGGAAGTAGGGCCTTCGCGCAC |
| 9 | Insertion | TTTC | FANCF | LbCas12a | GCGGATGTTCCAATCAGTACGCA | AATAGCATTGCAGAGAGGCGTATCATTTCGCGGATGTTCCAAGACTACAAATCAGTACGCAGAGAGTCGC<br>CGTCTCCAAGGTGAAAGCGGAAGTAGGGCCTTCGCGCAC |
| 12 | Insertion | TTTC | FANCF | LbCas12a | GCGGATGTTCCAATCAGTACGCA | AATAGCATTGCAGAGAGGCGTATCATTTCGCGGATGTTCCAAGACTACAAAGACTCAGTACGCAGAGAG<br>TCGCCGTCTCCAAGGTGAAAGCGGAAGTAGGGCCTTCGCGCAC |
| 15 | Insertion | TTTC | FANCF | LbCas12a | GCGGATGTTCCAATCAGTACGCA | AATAGCATTGCAGAGAGGCGTATCATTTCGCGGATGTTCCAAGACTACAAAGACGATTCAGTACGCAGA<br>GAGTCGCCGTCTCCAAGGTGAAAGCGGAAGTAGGGCCTTCGCGCAC |
| 18 | Insertion | TTTC | FANCF | LbCas12a | GCGGATGTTCCAATCAGTACGCA | AATAGCATTGCAGAGAGGCGTATCATTTCGCGGATGTTCCAAGACTACAAAGACGATGACTCAGTACGCA<br>GAGAGTCGCCGTCTCCAAGGTGAAAGCGGAAGTAGGGCCTTCGCGCAC |
| 21 | Insertion | TTTC | FANCF | LbCas12a | GCGGATGTTCCAATCAGTACGCA | AATAGCATTGCAGAGAGGCGTATCATTTCGCGGATGTTCCAAGACTACAAAGACGATGACGACTCAGTAC<br>GCAGAGAGTCGCCGTCTCCAAGGTGAAAGCGGAAGTAGGGCCTTCGCGCAC |
| 24 | Insertion | TTTC | FANCF | LbCas12a | GCGGATGTTCCAATCAGTACGCA | AATAGCATTGCAGAGAGGCGTATCATTTCGCGGATGTTCCAAGACTACAAAGACGATGACGACAAGTCA<br>GTACGCAGAGAGTCGCCGTCTCCAAGGTGAAAGCGGAAGTAGGGCCTTCGCGCAC |

Supplementary Figure 6a

| Modification (position) | Spacer Sequence | PAM | Locus | Enzyme |
|---|---|---|---|---|
| No Mismatch | CCTCACTCCTGCTCGGTGAATAA | TTTC | DNMT1 | LbCas12a |
| P12 | CCTCACTCCTGATCGGTGAATAA | TTTC | DNMT1 | LbCas12a |
| P13 | CCTCACTCCTGCGCGGTGAATAA | TTTC | DNMT1 | LbCas12a |
| P14 | CCTCACTCCTGCTAGGTGAATAA | TTTC | DNMT1 | LbCas12a |
| P15 | CCTCACTCCTGCTCTGTGAATAA | TTTC | DNMT1 | LbCas12a |
| P12,P13 | CCTCACTCCTGAGCGGTGAATAA | TTTC | DNMT1 | LbCas12a |
| P13,P14 | CCTCACTCCTGCGAGGTGAATAA | TTTC | DNMT1 | LbCas12a |
| P14,P15 | CCTCACTCCTGCTATGTGAATAA | TTTC | DNMT1 | LbCas12a |
| P12,P13,P14 | CCTCACTCCTGAGAGGTGAATAA | TTTC | DNMT1 | LbCas12a |
| P13,P14,P15 | CCTCACTCCTGCGATGTGAATAA | TTTC | DNMT1 | LbCas12a |

Supplementary Figure 10 (nicking guides used in the figure)

| Spacer Sequence | PAM | Locus | Enzyme |
|---|---|---|---|
| ACATCAGTACGAAAATGTTTCCT | TTTC | DNMT1 | RR-RE2 |
| CTCGCCTGTCATCTGGCGTGACA | TTTC | DNMT1 | RR-RE2 |
| CCTCACTCCTGCGAGGTGAATTT | TTTC | DNMT1 | RE2 |
| CCTCACTCCTGCTATGTGAATTT | TTTC | DNMT1 | RE2 |

**Supplementary Note 2**

High-throughput sequencing primers used in this study (shown without the overhang sequences used for Illumina barcoding PCR)

| Target Gene | Direction | Primer Sequence |
|---|---|---|
| DNMT1-primer pair 1 | Forward | CATGTTAAAAACACAACATCAGTG |
| DNMT1-primer pair 1 | Reverse | GTTTGAGGAGTGTTCAGTCT |
| DNMT1-primer pair 2 | Forward | AGGAAACATTAACGTACTGATGT |
| DNMT1-primer pair 2 | Reverse | CTGTGAGGATTGAGTGAGTTG |
| FANCF-primer pair 1 | Forward | GAGTCAAGGAACACGGATAAA |
| FANCF-primer pair 1 | Reverse | AAAGCGCCGATGGATGTG |
| FANCF-primer pair 2 | Forward | TCTCAAGCACTACCTACGTC |
| FANCF-primer pair 2 | Reverse | GTGGTAACGAGCTGCATC |
| RNF2 | Forward | CTTTATTTCCAGCAATGTCTCAG |
| RNF2 | Reverse | GTACCTACCACAAAAGCTTGTA |
| HEK2 | Forward | GACGTCTGCCCAATATGTAA |
| HEK2 | Reverse | TGAACTTCCCAAGTGAGAAG |
| EMX1-primer pair 1 | Forward | CAGCTCAGCCTGAGTGTTG |
| EMX1-primer pair 1 | Reverse | CGTGGGTTTGTGGTTGC |
| EMX1-primer pair 2 | Forward | AGAAGCTGGAGGAGGAAG |
| EMX1-primer pair 2 | Reverse | CTTGTCCCTCTGTCAATGG |
| DNMT1 off-target 1 (RE2) | Forward | TACGCCATGGGTGATAGTG |
| DNMT1 off-target 1 (RE2) | Reverse | CTGATACCAACAGAGTTGTG |
| DNMT1 off-target 2 (RE2) | Forward | AATCTGTGCACTCGGAGTG |
| DNMT1 off-target 2 (RE2) | Reverse | CCTTGTGTCCTTGACTGC |
| DNMT1 off-target 1 (RR-RE2) | Forward | AGAAATGCCAGGTTCAGC |
| DNMT1 off-target 1 (RR-RE2) | Reverse | CTTTGGAATTGAGCACGTG |
| DNMT1 off-target 2 (RR-RE2) | Forward | ATTTCTGAGAGCATGGTCTC |
| DNMT1 off-target 2 (RR-RE2) | Reverse | ACCATGAACACAGCAGATAG |
